# Suitable habitats of fish species in the Barents Sea

**DOI:** 10.1101/2020.01.20.912816

**Authors:** Bérengère Husson, Gregoire Certain, Anatoly Filin, Benjamin Planque

## Abstract

Many marine species are shifting their distribution poleward in response to climate change. The Barents Sea, as a doorstep to the fast-warming Arctic, is experiencing large scale changes in its environment and its communities. This paper aims at understanding what environmental predictors limit fish species habitats in the Barents Sea and discuss their possible evolution in response to the warming of the Arctic.

Species distribution models usually aim at predicting the probability of presence or the average abundance of a species, conditional on environmental drivers. A complementary approach is to determine suitable habitats by modelling the upper limit of a species’ response to environmental factors. Using quantile regressions, we model the upper limit of biomass for 33 fish species in the Barents Sea in response to 10 environmental predictors. Boreal species are mainly limited by temperatures and most of them are expected to be able to expand their distribution in the Barents Sea when new thermally suitable habitats become available, in the limit of bathymetric constraints. Artic species are often limited by several predictors, mainly depth, bottom and surface temperature and ice cover, and future habitats are hard to predict qualitatively. Widespread species like the Atlantic cod are not strongly limited by the selected variables at the scale of the study, and current and future suitable habitats are harder to predict. These models can be used as input to integrative tools like end-to-end models on the habitat preference and tolerance at the species scale to inform resource management and conservation.

## 1 INTRODUCTION

There is growing evidence of spatial shifts of species distribution correlated with climate change (Chen et al., 2011; Hickling et al., 2006; Parmesan and Yohe, 2003; Thomas, 2010). The Arctic is warming faster than any other ocean in the world (IPCC, 2014). Cheung et al. (2009) investigated the potential geographical changes in marine biodiversity worldwide in response to warming and suggested a general increase of species richness in Arctic waters due to northward migrations of species. The region would experience higher species turnover rates due to invasion and local extinction than anywhere on the globe. As such, describing distribution patterns, understanding drivers and projecting potential changes at the species scale is of crucial importance for conservation and management purposes (Marzloff et al., 2016; Pecl et al., 2017; Porfirio et al., 2014).

The Barents Sea is situated on the border of the Arctic. It is under the influence of two water masses outflowed from the warm, saline Atlantic in the south west and the cold, less saline Arctic in the north east (Loeng, 1991). Those two masses separate two main communities, arctic and boreal, that differ, among other thing, in fish species composition (e.g. Fossheim et al., 2015; Johannesen et al., 2012), traits (Frainer et al., 2017) and trophic structure (Kortsch et al., 2015). Important commercial fish (e.g. Atlantic cod, *Gadus morhua*, Atlanto-scandic herring, *Clupea harengus*, and capelin, *Mallotus villosus*) are present in both types of communities and respond to climatic signals (Chambault et al., 2018; Hamre, 1994; Matishov et al., 2012). During past decades, the Barents Sea has been experiencing an increase in Atlantic water inflow and coinciding heat content in the water column, as well as loss of sea ice in the northeast (Årthun et al., 2012; Dalpadado et al., 2012; Lind et al., 2018). In the meantime, the distribution of demersal fish has been altered with a general displacement of boreal communities towards the northeast (Fossheim et al., 2015). Unfortunately, studies in Arctic and subarctic waters have sometimes suffered from a lack of appropriate data to provide robust conclusions about changes in individual species biogeography (Ingvaldsen et al., 2015). Hollowed et al. (2013) estimated, based on expert knowledge, the potential of 17 species of the sub-arctic regions to shift their distribution northward, following the increased production in newly ice-free areas. This set of species was quite evenly divided into species with high, medium and low potential to change their distribution. However, changes in spatial distribution at the scale of individual species remain uncertain. There is a need for empirical studies that would help better identify the main drivers of changes in species distributions, which species are likely to respond most, and to which degree it is possible to make prediction about future geographical distributions based on currently available information.

Because environment is generally assumed to be the main driver of species distribution (Pearson & Dawson, 2003), most spatial distribution models (SDM) rely solely on environmental covariates. As such, they predict the suitability of a habitat to host a species based on its mean response (in presence/absence or in abundances) to environmental conditions. An alternative approach is to explicitly focus on how those factors may limit species habitats by predicting the upper limit of the species response, i.e. a high quantile (e.g. >0.9) instead of the mean. The statistical method of quantile regression (QR) (Cade et al., 1999; Cade & Noon, 2003) provides a useful framework for assessing limiting factors from observational data. It predicts the expected response for a given quantile *q*. With *q* = 0.5, QR predicts the median response. With high *q*’s (>0.9), QR predicts the upper limit of the response. For a set of environment conditions, it is possible to determine the most limiting factor by considering several single-covariate models and identifying the one that predicts the lowest response (Austin, 2007). This approach inherits from the Sprengel-Liebig law of the minimum (van der Ploeg et al., 1999), which considers that a response variable can only be as high as allowed by the most limiting factor.

Quantile regression originated in economics (Koenker and Bassett, 1978) and has been used in ecology to investigate prey-size-predator size relationship (Bethea et al., 2004), DNA variation across environmental gradients (Knight and Ackerly, 2002), response to metal concentrations (Schmidt et al., 2012), and fish recruitment-environment relationship (Planque and Buffaz, 2008). Review papers have highlighted its utility for the prediction of suitable habitats (Austin, 2007; Elith and Leathwick, 2009; Hegel et al., 2010), with some applications for terrestrial (Cade et al., 1999; Carrascal et al., 2016; Jarema et al., 2009; Schröder et al., 2005) and aquatic species (Ateweberhan et al., 2018; Cozzoli et al., 2013; Dunham et al., 2002; Lancaster and Belyea, 2006; Lauria et al., 2011; Vaz et al., 2008). Most of the ecological applications of quantile regression have assumed linear relationships between the biological response and the predictors. Based on theoretical considerations, the species response to an environmental factor is expected to be bell-shaped (Hutchinson, 1957; Whittaker, 1967) and recent studies have applied non-linear quantile regression models to allow for this (Anderson, 2008; Cozzoli et al., 2013; Dunham et al., 2002; Halkos, 2011; Schröder et al., 2005).

The aim of the present work is to (i) quantify the limiting effect of the environmental factors that impact on the spatial distribution of fish in the Barents Sea, (ii) assess the predictability of future geographical distributions based on currently available information and (iii) identify which species are most likely to respond to future environmental changes. For this purpose, we analyze data from the autumn ecosystem survey in the Barents Sea on the 33 most frequently sampled fish species and 10 environmental variables that can potentially limit their habitat. We develop QR nonlinear models for all combinations of species and environmental factors.

## 2 MATERIAL AND METHODS

### 2.1 DATA

#### 2.1.1 Fish biomass by species

Fishes were caught by a Campelen 1800 bottom trawl during the autumn IMR-PINRO joint ecosystem survey between 2004 and 2017 (Eriksen et al., 2018). The spatial extent is quite large (around 1.6 million km²), with 278 stations per year in average, depending on the sea ice extent in the North-eastern part of the sea. The sampling effort is regular in space with 35 nautical miles (35*1.852 km) between each trawling. The same stations are visited every year, in the limit of technical, time or climatic feasibility. A grid of 35 x 35 nm was fitted in an albers equal area projection, so that each grid cell contains only one station. The bulk of the fish species in the catch of the Campelen bottom trawl contained demersal species. However, bentho-pelagic and pelagic fish were also regularly caught in high abundances by the trawl and they are kept in the analyses. Estimated species biomasses were standardized by unit area, considering an opening of 25 m of the trawl. Only the trawls towed between 50 m and 500 m depth, in 15 to 60 minutes were kept. Towing speed was about 3 knots. In total, data comprised 3827 stations and 78 species over the 14 years. Taxa that were absent in more than 95% of the stations were removed, reducing the number of species to 33.

#### 2.1.2 Environmental predictors

Eleven variables reflecting the environmental conditions of fish habitat were gathered. Considerations on the nature and number of predictors to include according to sample size are described in appendix 1.

During the ecosystem survey, CTD were used to measure surface (10 m, T.surf) and bottom temperature (°C, T.bottom) and salinity (S.surf and S.bottom) at each station. Two stratification variables were calculated from temperature and salinity profiles following Planque et al. (2006). The surface mixed layer depth (SML, m) was calculated from a double layer model, and the potential energy anomaly (PotEnAno, kg.m-1.s-1, Simpson & Bowers, 1981) was estimated as the energy required to mix vertically the entire water column.

Bathymetry (m) and slope (degrees) were extracted from NOAA raster for the Barents Sea (Jakobsson et al., 2012). Sediment type was defined by extraction of seafloor description by NGU (Contains data under the Norwegian license for public data (NLOD) made available by Norway’s geological survey (NGU)). The 16 sediment classes described on the map were aggregated in 7 coarser classes following the EUNIS sediment hierarchical classification (Davies et al., 2004). Chlorophyll a (chla, mg/m^3^) average concentration between March and July of each year, as estimated by the NASA from ocean color (NASA OBPG, 2018). Number of days with ice cover (daysofice) were counted from daily sea ice extent maps from the NOAA (Cavalieri et al., 1996). For all those variables, values were extracted at the bottom trawl station position, i.e. there is only one of each per grid cell and per year.

Correlation analysis (described in appendix 2) showed a high correlation of potential energy anomaly with depth, so the former was removed from the analysis. To assess the potential of a species habitat suitability to be predicted in a changing environment, the ten remaining predictors were categorized into *fixed* (bathymetry, slope, sediment) and *dynamic* (all the others).

### 2.2 ANALYSIS

#### 2.2.1 Species response to environmental predictors

Prior to the regression analysis, species biomass data were log+d-transformed, where d is half the lowest biomass of the species. All quantitative environmental parameters were discretized in 20 categories of equal frequency to facilitate the model fitting process. In the case of days of ice cover, as there was a lot of 0, the first category comprised all the 0s, and the 19 others were spread equiprobably over the rest of the distribution.

One quantile generalized additive model (QGAM) was fitted for each pair of species-predictor (330 models) using the qgam function in the qgam package in R (Fasiolo et al., 2017) and setting the quantile level at 99%. The use of QGAMs allows for greater flexibility in the shape of the relationship between the predictor and the species response than linear quantile regression. It can capture bell-shaped responses, or responses that reach a plateau for high or low levels of the predictor. Other considerations about the theoretical roots, strengths and weaknesses of the methods are quickly described in appendix 3. To avoid regressions with shapes too complex to be ecologically meaningful, the number of degrees of freedom in the GAMs was limited to 3. For the qualitative variable (sediment), linear QR was applied to fit the 99^th^ quantile within each sediment category. The rq function from quantreg package in R (Koenker, 2018) was used.

Models were fitted using observations for years 2004 to 2013. They were then evaluated on observations for years 2014-2017. The evaluation was based on two metrics. The first metric is the proportion of observations in the evaluation dataset that were below the predicted 99th quantile. It is expected that 99% of the observations should fall below model predictions. If the observed proportion is higher, this means that the model is overestimating the maximum biomass (i.e. underestimating the limiting effect of the predictor). If the observed proportion is lower, too many observations in the evaluation dataset are higher than the expected maximum value, so the model is underestimating the maximum biomass (i.e. overestimating the limiting effect of the predictor). We categorized the variation from the 99^th^ quantile into a “slight” (98.5 to 99.5% of data below the predictions) and a “strong” (less than 98.5% or more 99.5%) over/underestimation of the maximum biomass. We considered that a model has a good predictive power if the predictions show a slight deviation from the 99^th^ quantile.

The second metric, termed ‘contrast’, is measured for each model on the predicted values, by the difference between minimum and maximum relative to the maximum. High (close to one) contrast occurs when the expected response of the species varies greatly across the environmental gradient. The predictor influences the species biomass, and has a limiting effect when biomasses are low. Low (close to zero) contrast occurs when there are little variations in the predicted species biomass along the environmental gradient. The predictor has a low effect on the species and is not limiting in the range of the Barents Sea. In the case of the sediment type, three of the seven classes (“Compacted sediments or sedimentary bedrock”, “Sand, gravel and pebbles”, and “Thin or discontinuous sediment on bedrock”) were associated to less than 1% of all the samples (appendix 4). Those sediment types are ignored for the calculation of the contrast to ensure that the metric is built on sediment categories that carry enough information.

#### 2.2.2 Spatial prediction of suitable habitats

Each year, it is possible to construct maps of habitat suitability for each species. Each station is associated with a set of predictor values. For a given species, each model predicts a 99^th^ quantile of biomass in response to that set of predictor values. The most limiting factor is the one leading to the lowest 99% quantile. From here on, we use the term “most limiting” factor as defined by this criterion, whether the predictors can have a direct (like bottom temperature and depth) or indirect limiting effect (like chlorophylle a, which is not in direct link with the species habitat, but is an indicator of primary production that can indirectly affect bottom species). The maximum (99^th^ quantile) biomass predicted based on the local environmental conditions is a local measure of habitat suitability. We applied this process to every location sampled each year.

This process results in two maps per year and per species: a habitat suitability map and a limiting factor map. The habitat suitability map displays the spatial distribution of the expected maximum biomass. The limiting factor map simply shows the most limiting factor at each location. However, when the biomasses are high, no factor can be considered limiting. In the limiting factor maps, wherever the maximum biomass predicted, at a given location, from the most limiting factor is superior to 25% of the species-predictor model maximum, we considered the factor to have a “weak limiting effect” on the species at the station. We use three categories to describe the limiting factors: fixed, dynamic, and weakly limiting (which can be both a dynamic or fixed predictor). From those maps, we looked at the proportion of locations where a given species biomass is limited by a given predictor. It is computed by i) counting for each predictor the number of stations where it is the most limiting for a given species a given year, ii) dividing that count by the number of stations sampled that year and iii) calculating the mean of that proportion over the years. It provides a measure of the limiting power of each predictor at the scale of the whole Barents Sea and across species.

#### 2.2.3 Predictability of future suitable habitats

To be able to predict a species suitable habitats in the Barents Sea using QGAMs, it is necessary that i) at least one selected predictor, dynamic or fixed, has an impact on the taxon response (i.e. the species-predictor model has a high contrast), ii) the value of the predictor(s) for which the species biomass is limited occurs in the study area and at the temporal scale of the study, iii) the modelled response is robust to new conditions (i.e. predicted maximum quantile on the evaluation dataset should be close to the 99^th^), iv) possible differences in specific respond of different groups of individuals (by age, size, physiological state and other) within a taxa to the environmental predictor(s) are avoided.

To evaluate the potential of a species suitable habitat to shift in a changing environment, we also look at the maximum contrast in fixed and dynamic predictor models. Species with a highest contrast in response to dynamic predictors are more susceptible to shift their habitat to follow changing environmental conditions.

## 3 RESULTS

### 3.1 RESULTS STRUCTURE

Norway pout (*Trisopterus esmarkii*) is used to illustrate the detailed results of the quantile regression on a single species in response to three predictors: two that are associated with a high and a low contrast in the species response and one qualitative predictor. This species was chosen because it showed high contrast in its responses to the selected variables and high consistency of the predicted quantiles between the training and the testing datasets.

The results for all other species analyzed in this study are provided in appendix 5. Tables summarizing the species responses to the different predictors are in appendix 6. Some figures use abbreviated species names. The correspondence between abbreviated and full names is provided in appendix 6.

A synthesis of the models of habitat suitability across all species is presented. In both parts, habitat suitability maps are shown only for 2013, which was the year with the widest spatial coverage. Maps for all the species are presented in appendix 7.

#### 3.2 LIMITS TO THE DISTRIBUTION OF THE NORWAY POUT (*TRISOPTERUS ESMARKII*)

##### 3.2.1 Species responses to environmental predictors

###### Convergence and predicted quantiles

*T. esmarkii* modelled response to depth, slope and sediment converged successfully. When fitted to the training dataset, 99.1% of the observations were below the modelled response to depth, and 99.0% were below the model for both slope and sediment (Figure 1). When the same models were applied to the testing dataset, 99.3% of the observations were below the depth model, and 99.4% below the slope model. Both models thus slightly overestimate the maximum biomass allowed by those predictors when applied to new environmental conditions. For sediment, the model strongly overestimates the maximum biomass of the predictor, with 99.9% of the data below the model.

###### Model contrast

The contrast in the response to depth was very high, 0.997. Such high value indicates that the minimum of the predicted maximum biomass was close to zero, i.e. that the sampling includes environmental conditions that are very limiting for the species. The response to slope shows the lowest contrast (0.81), indicating a relatively lower impact of slope on the species response. The contrast of the modelled response to sediment was intermediate with 0.90.

**Figure 1:**
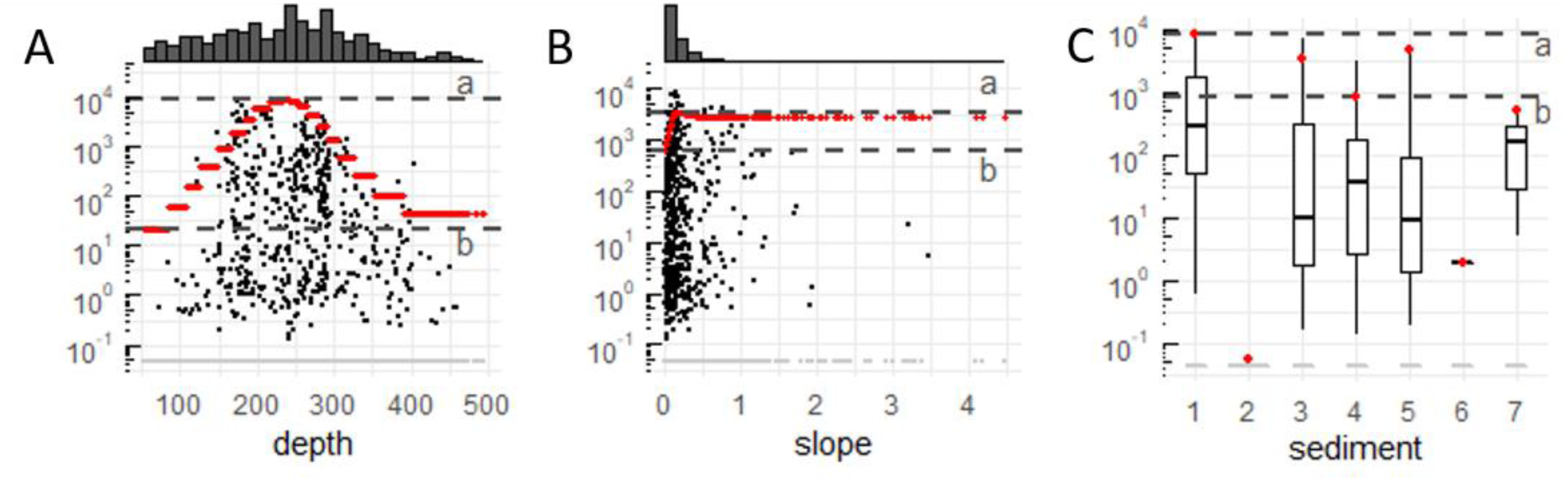
examples of Trisopterus esmarkii modelled log10 responses to three environmental predictors: depth, slope, and sediment type. A and B: black dotted scatterplot of the log of non-null biomasses of the species in response to the predictor. Red dots indicate modelled log of maximum biomass predictions. On top of the scatterplot, the marginal density shows the distribution of samples conditional to the predictor values. C: Boxplot of response to the sediment. The model prediction is the 99^th^ quantile for each sediment class: 1= Coarse sediment, 2= Compacted sediment or sedimentary bedrock, 3= Mixed sediment, 4= Mud, clay and sandy mud, 5= Sand and muddy sand, 6= Sand, gravel and pebbles, 7= Thin or discontinuous sediment on bedrock. All panels: horizonal dashed lines indicate references for the calculation of the contrast: a: max predicted maximum biomass, b: min predicted maximum biomass. Contrast = (a-b)/a.

###### Model shape

The responses of *T. esmarkii* to depth and slope are bell-shaped (Figure 1A and 1B respectively), although the response to slope is flatter. The “stair” pattern in the model predictions comes from the discretization of the predictors prior to fitting. The shape of the qualitative predictor, the sediment, is not relevant. The range of observations along the depth gradient is wide, covering 76% of the bathymetric range of the Barents Sea (from 73 m to 410 m depth, while conditions across the locations sampled scope from 52 to 494m). The maximum predicted biomasses occur at ∼240 m, while the minimal (i.e. the limiting values) occur at ∼70 m. For slope, most of the non-null biomasses were observed in areas with low degree of slope. For the sediment, the minimum prediction was close to 0 on compacted sediment or sedimentary bedrock, and maximum for coarse sediment.

###### Other predictors

All the modelled responses of *T. esmarkii* to the other predictors converged. Between 99.0% and 99.2% of the training dataset observations were below the model. The surface and bottom salinity and surface mixed layer depth models slightly overestimate the maximum biomass when applied to the testing dataset, with 99.2 to 99.4% of the data below the predictions. The other models more strongly overestimate the maximum biomass with 99.8 to 99.9% of the testing data below the models. Contrast is high for all the variables: from 0.73, 0.88, and 0.97 for surface mixed layer depth, *surface* and *bottom salinity* respectively to >0.99 for all other predictors.

The response of the species to surface temperature and salinity, bottom temperature, surface mixed layer depth, days of ice cover and chlorophyll *a* concentrations show complete or partial bell shapes (see appendix 5). The range of response to the different predictors is large, scoping from 55 to 96% of the conditions over the Barents Sea. Modelled response to bottom salinity shows a V shape. High biomasses of *T. esmarkii* are associated with warm surface (∼10°C) and bottom (>2.5°C) temperatures, high salinities (>34.5), rather shallow and weak stratification (SML ∼40-50m).

##### 3.2.2 Habitat suitability mapping

When applying the models for *T. esmarkii* for a given year, predictions are rather low (i.e. some factors are very limiting) in most of the Barents Sea, except in the south-west (e.g. in 2013, Figure 2A). Bottom temperature limited biomass in the majority (60.9%) of the stations in 2013 (58.3% in average across all years) and was most limiting in all the central area of the Barents Sea. Other environmental parameters are much less limiting. Depth (12.8% in 2013, 12.7% in average) is most limiting on the shallow areas of Murmansk bank and north of Bear Island, or in the depth of the Bear Island trough. Surface temperature was the third most frequent limiting factor in 2013 (9.0%); but second in average (14.0%). The ice coverage is the last predictor limiting more than 10% of the samples on average (11.6% on average, 5.4% in 2013). Both surface temperature and ice cover are most limiting in the north, between Svalbard and Franz Joseph Land. All other parameters are most limiting for less than 5% of all the samples in 2013 and on average over the years.

**Figure 2:**
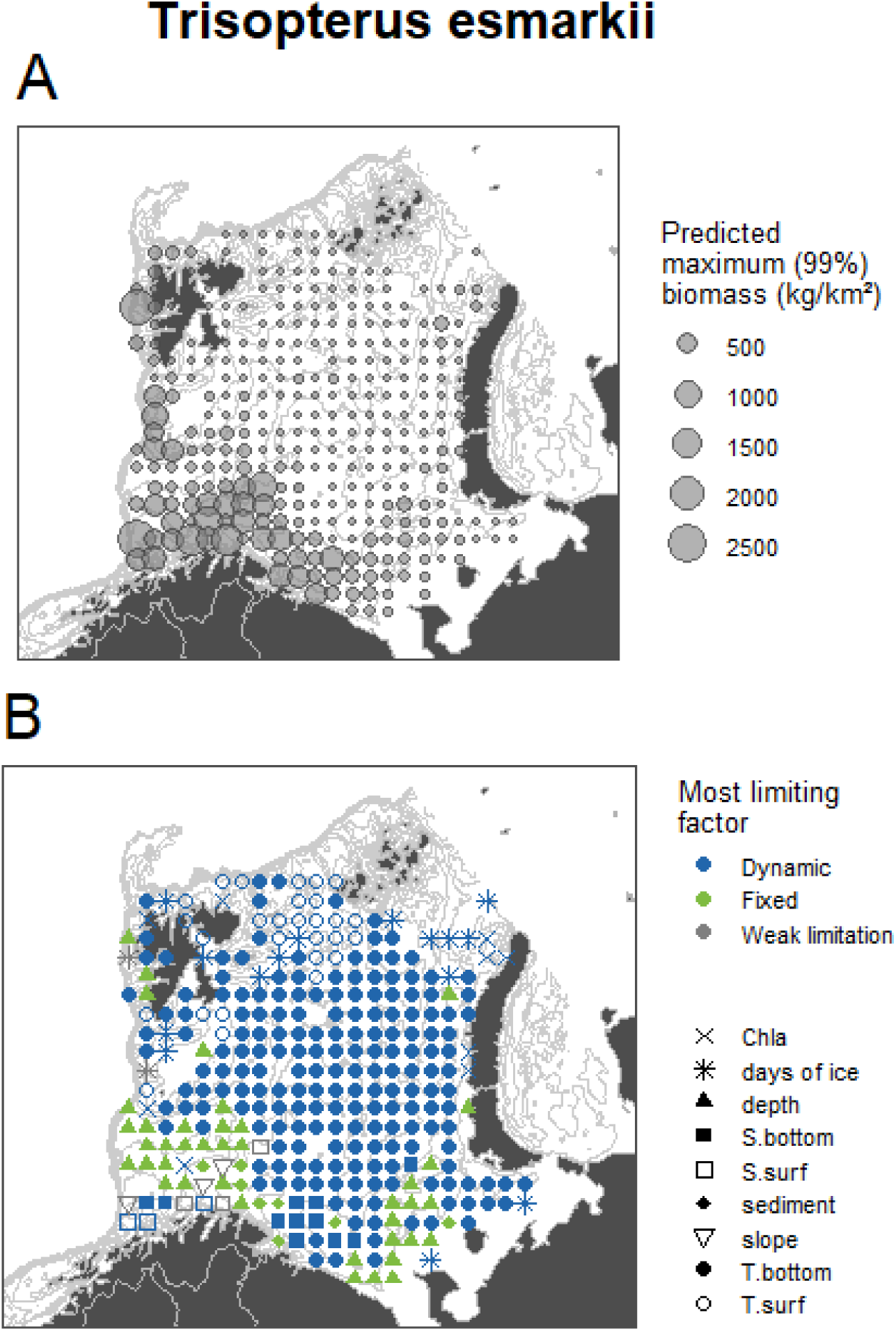
Spatial predictions in 2013 of Trisopterus esmarkii A) suitable habitat (maximum biomass) and B) most limiting predictor. Color indicates the predictor’s category: fixed (sediment, depth, slope), dynamic (all the others) or not weakly limiting. (predicted biomass > 25% of the model maximum)

##### 3.2.3 Predictability of the suitable habitats

The maximum contrast in the modeled response of *T. esmarkii* distribution was to depth (contrast: 0.991) among the static predictors and to bottom temperature (contrast > 0.999) among the dynamic predictors. However, depth is not that often a limiting factor in the Barents Sea. It is thus probable that this species suitable habitat will shift in response to changes in temperature, in the limit of the bathymetric constrains. The predictive power of the *T. esmarkii* – T. bottom model is poor as it tends to overestimate the maximum biomass, while that of the T. esmarkii – depth model is good. This means that the predicted habitat suitability might be overestimated if based only on bottom temperature.

#### 3.3 LIMITS TO THE DISTRIBUTION OF 33 FISH SPECIES

##### 3.3.1 Species responses to environmental predictors

###### Convergence and predicted quantiles

All models successfully converged. The training and testing sets performed quite differently on predicting the 99^th^ quantile (Figure 3). When fitted on the training set, most of the models (94%) were between 98.5% and 99.5% of the data. Only 6% strongly overestimated the maximum biomass (i.e. were above more than 99.5% of the data). None of them strongly underestimated the maximum biomass.

**Figure 3:**
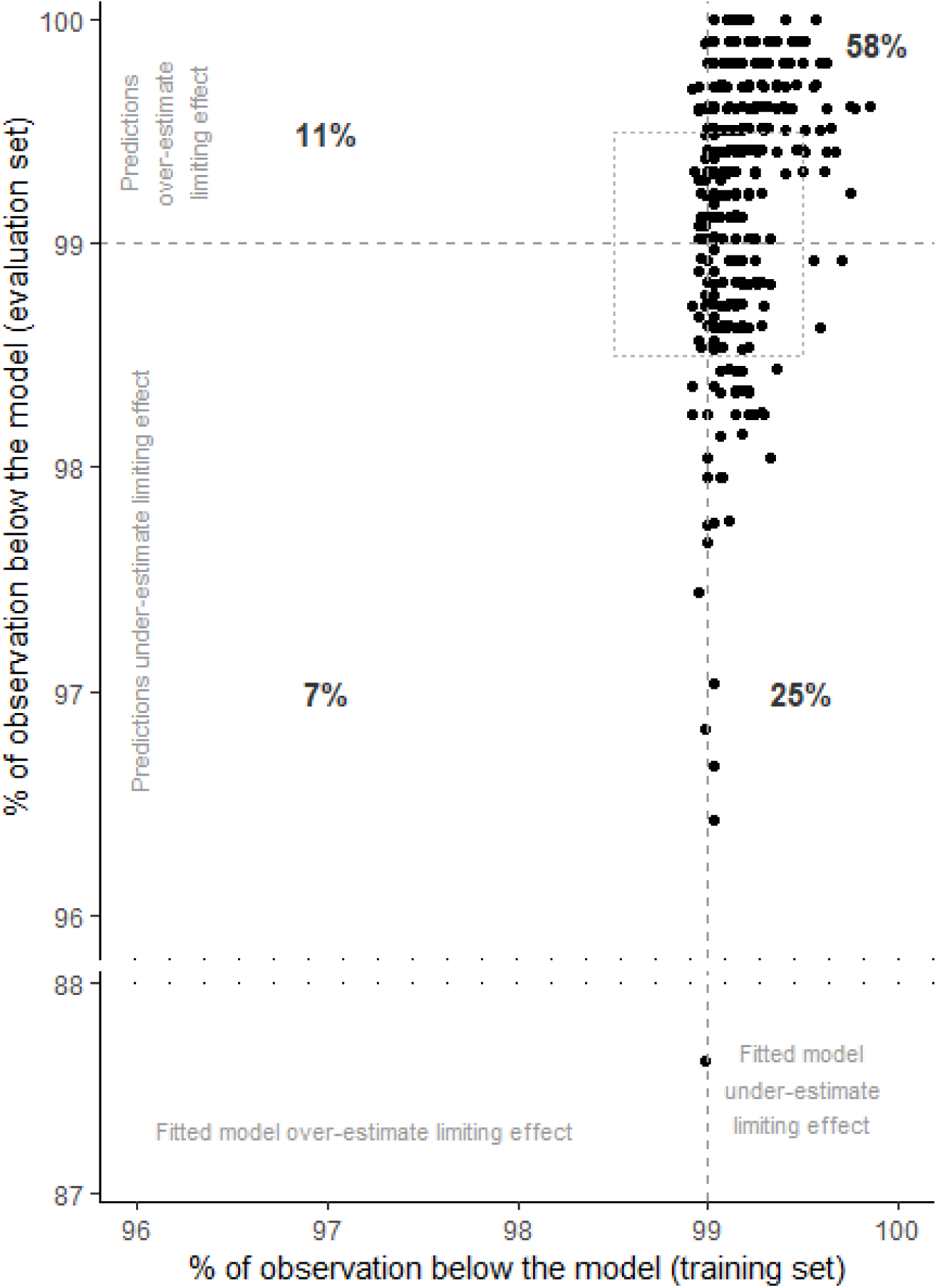
scatterplot of the predicted percentile of the observation by the models in the training and testing set. The dotted rectangle indicates models that slightly under or overestimate the maximum biomass. Outside of that rectangle are the models that strongly under or overestimate the maximum biomass.

The models performed less well at predicting the 99^th^ quantile when applied to the testing data set, as only 50% of the models were between 98.5 to 99.5% of the data; 38% strongly overestimated the maximum biomass and 11% strongly underestimated it. One model is an outlier, performing very poorly in the testing set: *Arctozenus risso* response to sediment. This may be because this species reached higher biomasses in 2016, during the testing set, than any other year before.

###### Model contrast

Most of the model showed relatively high level of contrast: 45% had high contrast (>0.90), and 37% had an intermediate contrast (0.50 to 0.90). Slope has the lowest mean contrast (0.44) across the 33 species, followed by surface mixed layer depth (0.58). Surface salinity and chlorophyll a are associated to similar contrast in the species response (∼0.75). Following predictors display high contrasts: sediment (0.83), ice cover (0.85), bottom salinity (0.86) and bottom temperature (0.87). Depth and surface temperature models are the most contrasted with an average of 0.90.

Among temporally fixed predictors, the most contrasted modeled responses were to depth (24 of the 33 species), sediment (8 species) and slope (1 species). Bottom and surface temperature caused the highest contrast among dynamic predictors for 12 and 13 species respectively, ice cover for 5 and bottom salinity for 3.

###### Model shapes

Most model shapes can be interpreted as a complete or a partial bell, with large differences in amplitude, from very contrasted to very flat models. Occasionally, species response models to surface or bottom salinity or ice coverage would take a v shape. Distribution of the species responses along the different predictor gradients can be found in appendix 5.

##### 3.3.2 Habitat suitability mapping

The mean proportion of samples limited by a single predictor over the years ranged from 0.3 to 58.8% (Figure 4, left panel). Some predictors limit on average 50% or more of a single species samples: bottom temperature (50.4% of *Argentina silus* samples, 58.3% of *Trisopterus esmarkii* samples), depth (53.9% of *Arctozenus risso* samples) and surface temperature (54.3% of *Triglops nybelini* samples).

**Figure 4:**
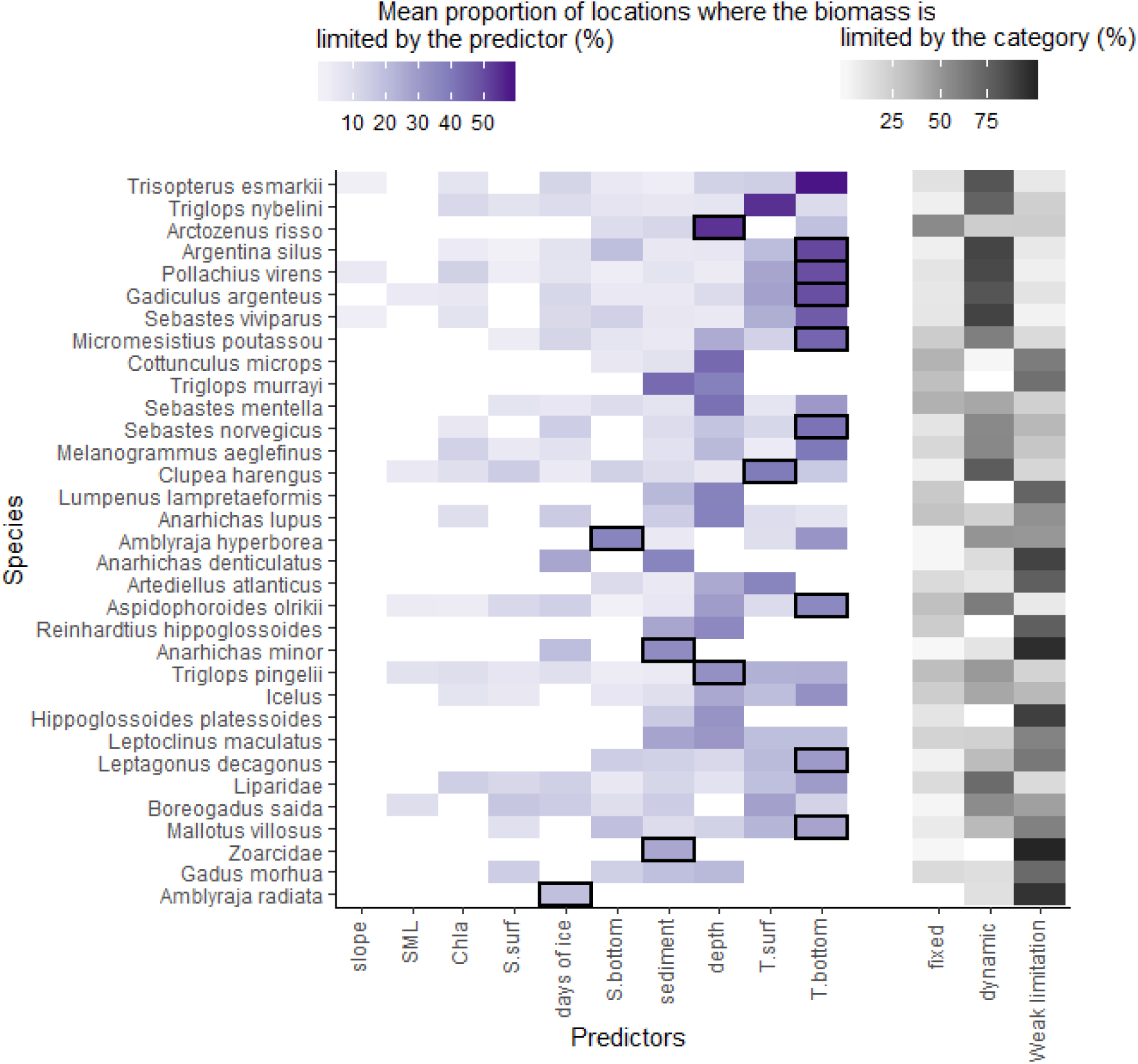
Frequency of the limiting effect across space and years. Right panel: for each species, mean proportion of samples limited by each predictor category: fixed, dynamic, or weakly limiting. Fixed parameters are slope, depth and sediment. Dynamic predictors are all the others. Predictors are weakly limiting a sample if the corresponding predicted biomass is >25% of the model maximum. Left panel: for each species, mean proportion of samples limited by each factor. Only the samples strongly limited are shown (predicted biomass <25% of the model maximum). Black rectangles identify for each species the most frequent most limiting predictor, but only if its predictive power is acceptable (predicted quantile of the testing dataset ranging from 98.5 to 99.5).

Bottom temperature is the most frequent most limiting predictor (22% of all samples). Depth is most limiting in 20% of the samples, and surface temperature and sediment in 14%. Slope (2%), surface mixed layer depth (2%), surface salinity (4%), chlorophyll a (5%), bottom salinity (7%) and ice cover (8%) are the least limiting among the species. However, these predictors are not always strongly limiting the species biomasses. For 15 of the 33 species, environmental conditions weakly limit the biomass in most of the sampled locations (Figure 4, right panel). Dynamic predictors are most often limiting for 17 species and fixed ones only for 1 (*Arctozenus risso*).

##### 3.3.3 Predictability of suitable habitats

Profiles of species responses to the selected predictors vary a lot in the Barents Sea (Figure 4). Some species are strongly limited by a low number of predictors, mainly dynamic ones (Figure 4, top species), while others are rather evenly limited by several predictors (Figure 4, bottom species). Most frequent most limiting predictors that have good predictive power (predicted quantile of the testing dataset between 98.5 and 99.5) are sediment, depth and bottom and surface temperature and bottom salinity. Nearly half (15 out of 33) of the considered species are most frequently most limited by a predictor for which the model has good predictive power.

It is the case of species situated toward the top of figure 4 for which we can thus evaluate current suitable habitats. They are limited by a low number of parameters. Those are e.g. *Arctozenus risso*, *Argentina silus*, *Pollachius virens*, *Gadiculus argenteus* and *Micromesistius poutassou*.

*Trisopterus esmarkii*, *Triglops nybelini* and *Sebastes viviparus* are also strongly limited by few predictors, but their predictive power is less good so the model might over or underestimate the habitat suitability.

Species for which it is hard to decipher suitable from unsuitable habitats are situated toward the bottom of figure 4 and most of their sample are only weakly limited: e.g. *Amblyraja radiata*, *Gadus morhua*, *Zoarcidea, Hippoglossoides platessoides, Anarhichas minor* or *Anarichas denticulatus*. Although some are limited by few predictors, and despite the good predictive power of the corresponding models, those species tend to be mostly weakly limited by the environmental variables, i.e. display high predicted biomasses on most of the Barents Sea. For 21 of the 33 species, the maximum contrast to dynamic predictors is higher than that of the fixed ones (Figure 5). This maximum predictor is bottom salinity for 1 species, ice for 3, bottom temperature for 8 and surface temperature for 9. All those species are thus more susceptible to shift their habitat to follow a change in the environment, particularly those with the highest maximum contrast. The 12 other species have higher contrast in fixed predictors. The maximum predictor is depth for all those 12 species. Those are more constrained by depth and their habitat might not be influenced by a change in dynamic environmental conditions.

**Figure 5:**
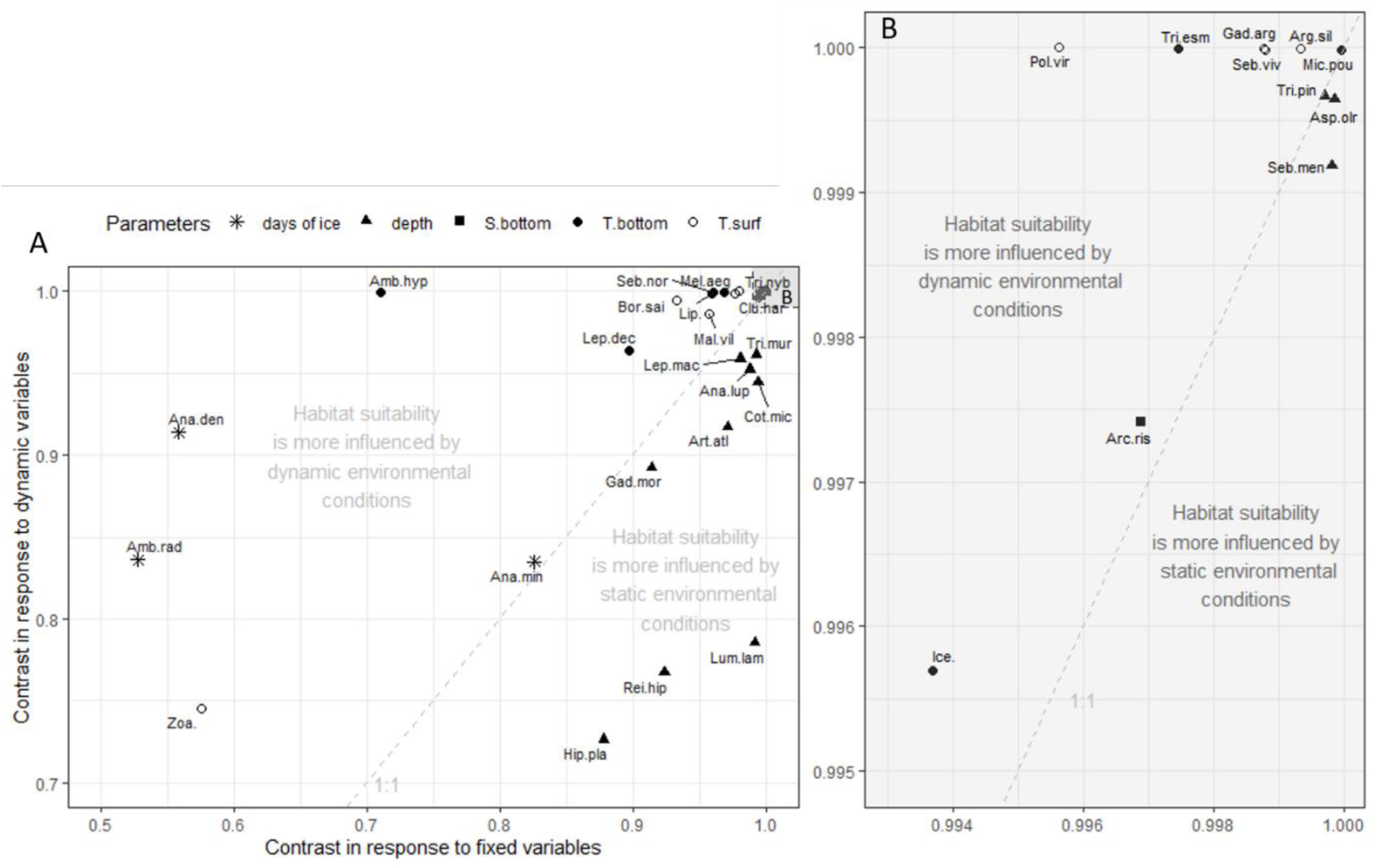
Scatterplot of species maximum contrasts in response to fixed versus dynamic variables. The shape of the point indicates which parameter is the one with the maximum contrast. The 1:1 line is grey and dashed. B is a zoom in the grey area of A.

## 4 DISCUSSION

In the present work we explored the limiting effect of 10 environmental predictors on the individual responses of 33 fish species of the Barents Sea and assessed our capacity to predict their suitable habitats. From the results, we can estimate the species ability to track potential changes in their suitable habitats in response to climate change.

### 4.1 LIMITING EFFECT OF ENVIRONMENTAL PREDICTORS ON INDIVIDUAL

#### SPECIES SUITABLE HABITATS

The shapes of QGAM models provide the information about the limiting effect of predictors on species. In this study, QGAMs were fitted with a maximum degree of freedom of 3, so that the resulting models display simple shapes that can be interpreted in the context of the niche theory. Most frequently, models display bell shapes that can sometimes be skewed and/or incomplete (i.e. only one side of the bell is visible). V-shapes occur occasionally (in response to salinity or ice cover) and are more difficult to interpret. Causes of those v shapes could include the existence of two population within the Barents Sea with different habitat preferences, or strong non-linear links to other variables with strong spatial structure (proximity to coast, river outflow, depth, etc.).

The flatness of the model shape is an indicator of the limiting power of the predictor and is reflected in the contrast metric. Some predictors are more contrasted (i.e. limiting) than others. Depth and surface and bottom temperature have the highest average contrast over the considered species. This is consistent with the literature, as many authors have highlighted the importance of depth and temperature in the habitat requirements of demersal fish over the world (see Johnson et al., 2013 for a review). In the habitat suitability maps, they are also the most frequent most limiting predictors across the study area. The reason for these three parameters to limit the distribution of shallow-water species (< 200m depth) in depth and latitude is well studied (Brown & Thatje, 2015; Pörtner, 2010) and is linked to the thermal, oxic and hydrostatic conditions necessary for those species to maintain aerobic metabolism. Depth has been reported to be one of the most, if not the most, important predictor of demersal fish distribution, regardless of the method used or the geographical location of the study (Chatfield et al., 2010; Leathwick et al., 2006; Moore et al., 2010, 2011; Ross et al., 2015; Rutterford et al., 2015; Smoliński & Radtke, 2017). In this study, depth was often limiting either on the shelf for species living in deep areas (e.g. spotted barracudina *Arctozenus risso* or deepwater redfish *Sebastes mentella*, mainly found in the Bear Island Trough), or on the deepest and shallowest areas (for e.g. snakeblenny, *Lumpenus lampretaeformis,* or wolfish *Anarhichas lupus*).

Limiting values of bottom temperature for the distribution of the demersal fish occur often in the Barents Sea. For species that are distributed in the south west part of the Barents Sea (e.g. Norway pout *Trisopterus esmarkii,* greater argentine *Argentinus silus,* saithe *Pollachius virens,* silvery pout *Gadiculus argenteus*) this predictor was the most limiting in more than half of their samples. Their spatial distribution appear to be limited by the low bottom temperatures currently occurring in the rest of the Barents Sea. Byrkjedal & Høines (2007) obtained similar results in a study focusing on the south-western part of the Barents Sea, and explained the strong influence of the temperature by the conjunction of the cold, subzero, Artic and warm Atlantic water at the polar front, creating strong latitudinal gradients of temperature.

Surface temperatures cause high contrast in the species response are frequently the most limiting either i) in the north, approximately northeast of the polar front, for some species distributed mostly in the south of the Barents Sea (e.g. Atlantic herring *Clupea harengus* and Saithe *P. virens*), or ii) in the south for species considered as arctic (e.g. polar cod *Boreagadus saida* and bigeye sculpin *Triglops nybellini*). Some of those species have been shown to follow yearly variations in sea ice extent in other sub-arctic areas (Wyllie-Echeverria and Wooster, 1998). In our samples, surface temperature and ice cover are often limiting in the same area, in the North, so the limitation of the species responses by low surface temperature might also be a proxy of the limitation by cold, ice covered water masses north of the polar front.

The most limiting factors of species suitable habitats revealed by the QGAMs are consistent with the literature and reflect the strong environmental gradients existing across the Barents Sea.

### 4.2 ASSESSING OUR CAPACITY TO IDENTIFY SPECIES SUITABLE HABITATS

All the species are not impacted in the same way by the different predictors, and suitable habitats are thus not equally identifiable across species.

Some of the species have a taxonomic resolution too coarse to ensure a uniform response to the predictors across all individuals. A recent study (Smith et al., 2019) showed that grouping related taxa that are likely to share environmental tolerances, or splitting species in smaller population units that have adapted independently can improve niche estimates. In the case of cod (*Gadus morhua*), or eel pouts (zoarcids) the widespread spatial distributions and environmental tolerance partially reflect the variety of habitats used by different age groups (cod) or species (eel pouts). Modelling habitat suitability at a finer biological scale (e.g. by age or species) might be required to improve habitat suitability models for these groups (M. McPherson & Jetz, 2007; Morán-Ordóñez et al., 2017; Porfirio et al., 2014; Thuiller et al., 2005). In addition, suitable habitats are also hard to identify for species that are abundant and widespread like Long rough dab (*Hippogloissoides platessoides*), Greenland halibut (*Reinhardtius hippoglossoides*), Thorny skate (*Amblyraja radiata*) and two species of wolfish (*Anarhichas minor* and *denticulatus*). Long rough dab inhabits most of the Barents Sea and operates an east to west spawning migration against the larval drift, which allows it to maintain its position in the region (Walsh, 1996). This shows its wide tolerance for the conditions in the Barents Sea. The habitat mapping in the current study show that Long rough dab and Greenland halibut are never strongly limited by environmental conditions, except by extreme depths in shallow (for Greenland halibut) or deep areas (for Long rough dab). Thorny skate, on the other hand, thrives in all ranges of depth and temperatures of the Barents Sea (Dolgov et al., 2005). All those species are very abundant across the whole Barents Sea and thus mostly weakly limited by selected environmental factors. There is therefore substantial information on where these species are, but little on where there aren’t. It is thus difficult to identify their unsuitable habitats and how environmental conditions may limit their distributions. Species for which it is possible to identify suitable habitats are e.g. the spotted barracudina (*Arctozenus risso)*, the greater argentine (*Argentina silus)*, saithe (*Pollachius virens)*, silvery pout (*Gadiculus argenteus),* Norway pout (*Trisopterus esmarkii*), Norway redfish (*Sebastes viviparus*), or Blue whiting (*Micromesistius poutassou*). Those are exclusively south-western species inhabiting rather deep areas with warmer Atlantic bottom waters at the entrance of the Barents Sea. For example, blue whiting and adult saithe resides in the Norwegian Sea and expands into the Barents Sea when the Norwegian stock is large (for blue whiting: Heino et al., 2008) or as a seasonal migration during the second and third quarter (for saithe:Olsen et al., 2010). For all those boreal species, suitable habitats are mainly limited by only one predictor (most of the time the bottom temperature). *Triglops nybelini* is the only arctic species for which there is a clear limitation by a single factor, the surface temperature, which highly linked to ice cover in the north-east.

Some species habitats can be determined even though each predictor limits only a small portion of samples; i.e. there is no clear limitation by a single factor. For those species, the proportion of samples that are weakly limited by the environment is not as important as for widespread species, so we have some information on where the species is absent, or in low abundances. It is the case for the habitats of polar cod (*Boreogadus saida*), capelin (*Mallotus villosus*), eel pouts (liparids), Atlantic poacher (*Leptagonus decagonus*), daubed shanny (*Leptoclinus maculatus*) or scaled sculpin (*Icelus* spp.). Those are mainly arctic species, abundant but not widespread in the Barents Sea, spatially limited to colder waters north of the polar front (Fossheim et al., 2006; Hop & Gjøsæter, 2013). We can determine suitable habitats, but we need for that to consider several predictors.

The biogeography and the environmental affinity and tolerance of a species in the Barents Sea seems to be major indicators of our capacity to identify its habitat. Together with the results of the current study, this help us build hypotheses on the potential shifts in suitable habitats for individual species of the Barents Sea.

### 4.3 PREDICTING FUTURE HABITAT SUITABILITY IN RESPONSE TO CLIMATE

#### CHANGE

Quantitative predictions of suitable habitats for fish in the Barents Sea can be obtained by applying the QGAM models on projected maps of predictors showing the possible future environmental conditions. The predictive power of a model determines how well it will perform when transferred to a new area or another time, which is particularly important in the context of climate change (Dormann, 2007; Porfirio et al., 2014). Although it is not possible to quantitatively assess model’s performance in future climate, the recent rapid warming in the Barents Sea provides suitable conditions to test the performance of the habitat models in two periods with contrasting ocean climate. Half of the models performed well when applied to the testing dataset. The poorer performances of the other models may reflect that the training dataset did not include enough of the variability in the species response to the predictor. For those models, prediction can still be done, but the resulting habitat suitability might be over/underestimated.

Without projected environmental maps, it is still possible to use the results from the QGAM fit to hypothesize qualitatively the evolution of Barents Sea fish suitable habitats in response to environmental changes. Recent climate predictions show increasing water temperatures in the Barents Sea (Stenevik & Sundby, 2007) and decrease in sea ice possibly leading to ice free winters by 2061-2088 (Onarheim & Årthun, 2017). Species that would be more sensitive to these projected changes, i.e. that would be forced to move to track suitable habitats, are those that display a highest contrast in response to dynamic - rather than fixed - variables. This is the case for two thirds of the species. The limiting factors are bottom and surface temperature (that is projected to increase with climate change), bottom salinity (correlated to depth) and ice cover (which is projected to decrease). However, species tracking their environment might be limited in their progress by unsuitable fixed environmental conditions. A good example is *Anarhichas lupus*, which responds with the highest contrast to ice but is more often limited by depth across the Barents Sea. Predicting its future suitable habitat necessitate to consider both fixed and dynamic parameters. This supports a recent study projecting that depth will strongly limit the availability of suitable habitats (Rutterford et al., 2015).

Predicting potential shifts in suitable habitat for a species thus requires integrating all the information gathered in the current study on niche preferences and ranges, most contrasted models, spatially most limiting factors, response to dynamic and fixed factors and predictability of suitable habitats. Here we make tentative qualitative predictions on the future of demersal fish in the Barents Sea based on the two most limiting predictors of the region: bottom temperature and depth (Figure 6).

**Figure 6:**
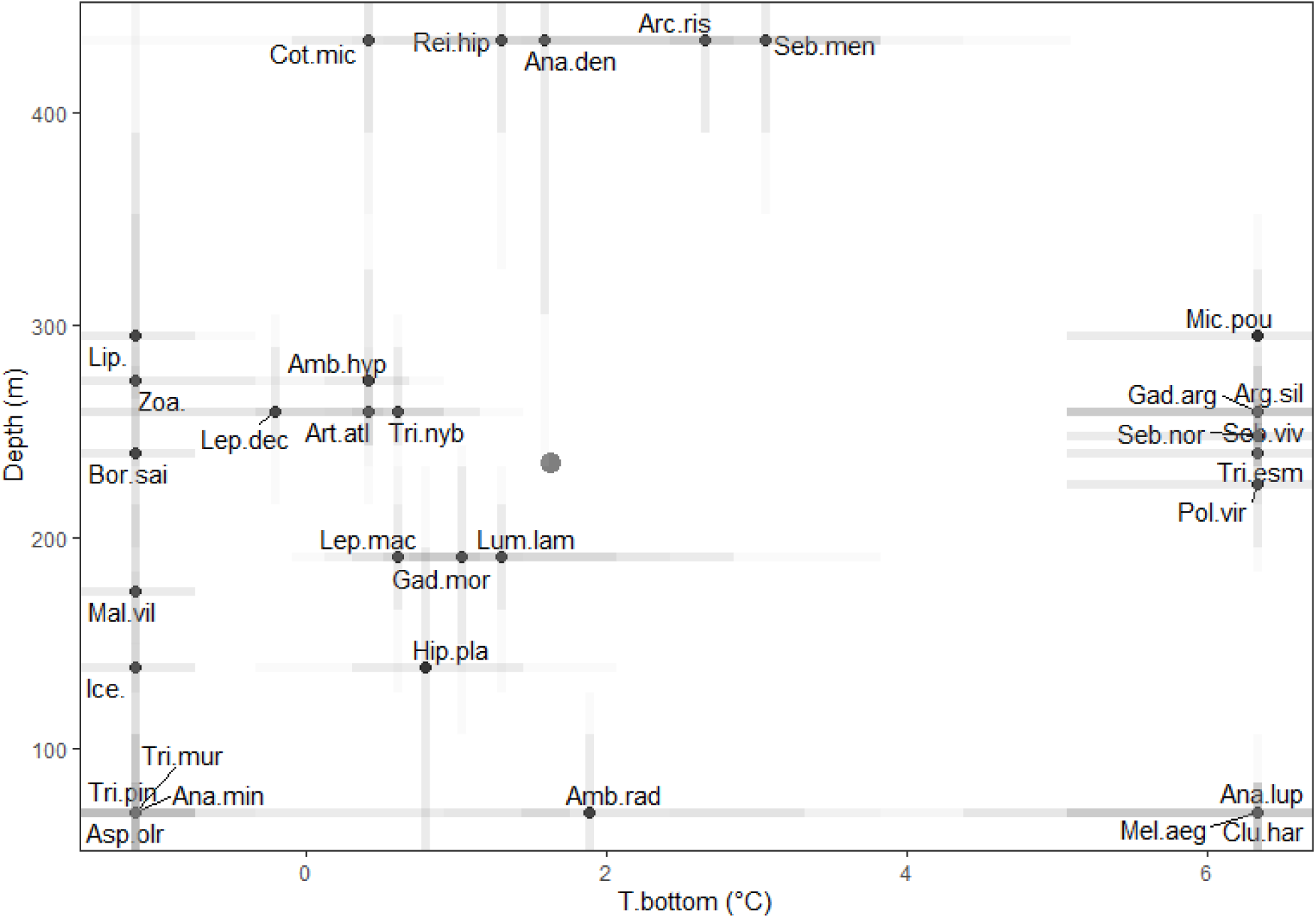
Bottom temperature - depth habitat preferences of the Barents Sea demersal fish. Dots indicate the predictor value at which the species predicted biomass is maximum. Grey lines indicate the width of the species optimum i.e.for which predictors values the predicted biomasses reach 90 (dark grey) and 80% (light grey) of their maximum. Big grey dot indicates the average values of depth and bottom temperature in the Barents Sea.

The warming of the Barents Sea is likely to increase the extent of suitable habitat for species with preferences for warmer waters (right side of the figure 6). They are susceptible to migrate or expand northward as new habitats become available, if the depth is suitable. The species concerned respond strongly to and are spatially more limited by either bottom or surface temperature. They are species for which it is easy to estimate qualitatively their future habitat because few predictors control their niche et the scale of the Barents Sea. Species that prefer intermediate depths (*Micromesistius poutassou, Argentina silus, Gadiculus argenteus, Sebastes norvegicus* and *viviparus*, *Trisopterus esmarkii* and *Pollachius virens*) and two shallower species (*Clupea harengus* and *Melanogrammus aeglefinus)* are likely to migrate north as most of the Barents Sea is in the range of their suitable depths. This is supported in the literature. Ecological niche models have predicted a gain in suitable habitat in the Barents Sea for saithe (*P.virens*) and haddock (*M. aeglefinus*) in the middle of the Barents Sea between 1960 and 2090 (Lenoir et al., 2011). (Hollowed et al., 2013) also hypothesized a northward shift of *C. harengus*. Some of those species have already been noticed to displace northward: (Perry et al., 2005) noticed that *M. poutassou* and *T. esmarkii* distribution boundaries have shifted northward in relation to the warming between 1977 and 2001 in the North Sea, and in the Barents Sea, both species and *A. silus* are part of the boreal or intermediate communities that also have shifted between 2004-2012 (Fossheim et al., 2015). Unlike this group of species, *A. lupus* is already widespread on the shallow banks of the Barents Sea and is spatially limited by depth and sediment in its northern boundary. It is thus unlikely that the warming will open new suitable habitats for that species. However, this species is also limited by other parameters so future suitable habitats are hard to predict.

Temperature increase in the Barents Sea will cause the loss of the coldest habitats of the region. Species that prefer cold habitats (left side of figure 6) are the most threatened as they will then experience temperatures warmer than their current optima. To come back to temperatures closer to their optimum, they would need to migrate further north into the deep Arctic ocean, or retract around Svalbard where they would ultimately be trapped if they don’t tolerate high depths. The concerned species are mainly arctic ones with large depth tolerance, so both scenarios are possible. Ribbed sculpin (*Triglops pingelii)* and Arctic alligatorfish (*Aspidophoroides olrikii)* are exceptions as they respond more strongly to depth and might not be able to retract to deeper and colder areas. However, all those species are part of the group for which suitable habitats are harder to predict qualitatively because of the many predictors involved in the biomass limitation. To understand potential shifts in their future habitat, the knowledge gained on their habitat requirements needs to be integrated and applied to projected environmental conditions.

Species currently preferring intermediate temperatures (0 to 2°C) can be divided into shallow and intermediate depth loving species and deep associated ones. Shallower species (optimum >300m) are generally widespread. Some are common over the whole Barents Sea (like *Gadus morhua*, *Hippoglossoides platessoides* or *Ambyraja radiata*) and little can be said about their future habitat. Others are widespread in the north and south-east of the Barents Sea (*Leptoclinus maculatus, Lumpenus lampretaeformis, Leptagonus. decagonus, Artediellus atlanticus, Triglops nybellini* and *Amblyraja hyperborea*). They all tolerate a wide range of depth conditions so they would able to track their preferred environmental conditions by moving northward or diving deeper. However most of them are also limited by other predictors, so their future suitable habitats is hard to estimate qualitatively.

Deeper species (*Cottunculus microps, Reinhardtius hippoglossoides, Anarhichas denticulatus, Arctozenus risso* and *Sebasted mentella*) are mainly found around the Bear island Trough. Their response to climate change depends more on their tolerance to shallower depths. *R. hippoglossoides* and *A. denticulatus* are widespread, with wide tolerance to depth and might be able to expand northward. In Hollowed et al. (2013), *R. hypoglossoides* is indeed suspected to move in or expand in the high Arctic. *S. mentella* is more constrained by shallower depths but has expanded into the Barents Sea during the period of the study (as hypothesized by Hollowed et al., 2013). *A. risso* however, is not very tolerant to shallower depth and respond strongly to salinity (which is itself very correlated to depth). If its habitat conditions were to change, the species could not move northward on the shallower Barents Sea shelf.

Similar tradeoffs will constantly occur for all species as changes in dynamic variables will interact with limitations caused by fixed ones. Light conditions might be a particularly strong tradeoff at those latitudes (Poloczanska et al., 2016).

## 5 CONCLUSIONS

The use of QGAM allowed to explore the potential environmental niche of 33 fish species in the Barents Sea. The models show a wide variety of responses to environmental stressors. The application of the Liebig’s law on the mapped conditions of the region highlighted the importance of depth and temperatures as limiting factors for most of the species. But the set of selected predictors influence each taxon differently, which leads to some species suitable habitats being more difficult to predict than others. While species responding more strongly to dynamic variables should be the most responsive to changes in their habitats, this study highlighted the importance of considering their interaction with fixed predictors when predicting future suitable habitats.

In the face of the complexity of the response at the individual species scale, it seems clear that explaining and predicting the responses of whole communities to changes in their habitat is challenging. Yet, ecosystem studies need for those individual responses to be integrated at larger scales. An advantage of the QGAM methods is that the models can easily be used as habitat preferences prior that input end-to-endo models. This would allow to predict suitable habitats maps on top of which other processes would refine the species distribution. Such empirical knowledge at the basis of the modelling process would greatly benefit our models and can inform resource management and conservation.

## Supporting information

Appendix 5 QGAM models

Appendix 6 niche descriptions

Appendix 7 Habitat suitability maps

## Acknowledgments

This work is part of the BSECO project funded by Arktis2030 (BSECO Barentshavet i endring Contract nr. QZA-15/0137). We are grateful to Dr. Randi Ingvaldsen for her help with collecting the ice cover data. We are also grateful to all the people involved in the joint ecosystem surveys at the Institute of Marine Research (IMR), Norway, and Polar Branch of the Federal State Budget Scientific Institution (PINRO), Russia.

## 7 TABLES

No tables in this paper

## 9 APPENDICES

Supplementary material can be found as PDF on the server

