## Appendices text and captions for "Suitable habitats of fish species in the Barents Sea"

#### 1 Appendix 1: Environmental variables to include based on the literature and our sample size

Eleven predictors were considered. Bottom temperature and salinity, sediment, slope and bathymetry were selected as they reflect the direct environment of demersal fish. Chlorophyll a is a proxy of the local productivity. Ice cover, surface temperature and salinity and stratification are indications of the water and nutrients circulation in the water column and the resulting potential bentho-pelagic coupling.

For demersal fish, most of the studies focusing on the habitat requirements over the world highlighted the importance of depth, salinity, temperature, hydrography, substrate and other abiotic variables like light or dissolved oxygen concentration (Johnson et al., 2013). Most of those variables have been used here, except for hydrography, light and oxygen concentration. Currents were considered but not included because of the lack of measurements at the station level, while using model outputs would have introduced a bias and possible multicollinearity in the model. Light would have been an interesting variable as it constraints numerous processes like primary production, visual predation (Langbehn and Varpe, 2017) or zooplankton depth in the Barents sea (Aarflot et al., 2019). However, a proxy of primary production was already included in the selected variables with the chlorophyll a.

Sample size has been shown to influence greatly the predictions, depending on the type of model (Hernandez et al., 2006; Stockwell and Peterson, 2002; Thibaud et al., 2014; Wisz et al., 2008). Harrell et al., (1996) suggested that the ratio of the number of presences to the number of predictors should be of 10 for 1. Here, we kept only species with at least 5% of occurrences among the 3827 samples, so all our species should be present in about 194 samples, for 10 predictors (for which we would need 100 samples). We could thus consider that we have enough data to produce suitable predictions.

### 2 Appendix 2 : Environnemental variables correlation analysis

As common SDM methods, quantile regression is also sensitive to collinearity of the predictors and their selection should be based on sound knowledge of the mechanisms involved (Austin, 2007; Dormann, 2007; Merow et al., 2014)

We conducted a simple Pearson's correlation analysis with rcorr and corrgram functions from the Rcmdrmisc and corrgram R libraries respectively (Fox, 2018; Wright, 2018). Results show a high correlation between depth and potential anomaly deficit

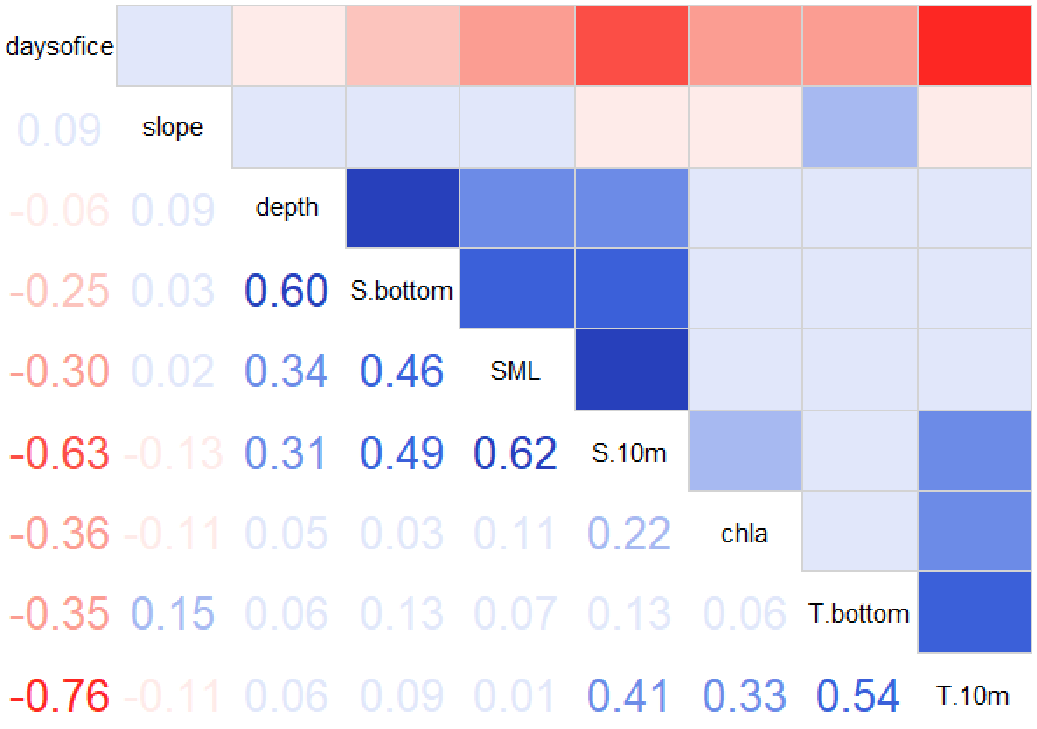

### 3 APPENDIX 3: ABOUT QUANTILE REGRESSION

#### 3.1 LINK TO THEORY

Quantile regression on GAMs is a flexible tool with strong theoretical background that allows to model any quantile of species-environment relationships (Cade et al., 1999; Cozzoli et al., 2013; Vaz et al., 2008).

(Hutchinson, 1957) defines the fundamental niche as an *n*-dimensional hypervolume that determines the values along *n* gradients in which a species can survive. Important subdivisions of the fundamental niche are the potential niche, which is the part that is available in the environment (Ackerly, 2003), and the realized niche, which is the restriction of the fundamental niche by the existing biotic and abiotic pressures (Hutchinson, 1957). These various pressures occur in permanence in nature, and when building SDMs, the observed distribution of a species is hypothesized to represent the geographical realized niche (Guisan and Zimmermann, 2000).

QGAM on a high quantile models the maximum response of a species to an environmental gradient. When comparing predicted responses to a set of environmental conditions of a location, it assesses if the habitat is *a priori* suitable for a species, even if the species is not always observed at these locations. It thus describes potential rather than the realized niche (Cade et al., 2005; Jiménez-Valverde et al., 2008).

#### 3.2 STRENGTHS AND WEAKNESSES OF QUANTILE REGRESSION

QR methods have several advantages with regards to SDM. First, they do not need complex model selection procedure, as each environmental effect is modeled one by one. They are not as sensitive as conventional SDM to the existence of unmeasured factors (M. P. Austin & Niel, 2011; Mod et al., 2016), as long as the most limiting factors are included (Cade et al., 2005). However, the method does not consider factor interactions, including possible co-limitation or multiple limitation occurring at the same time (Rubio et al., 2003). As any other SDM, it is also sensitive to collinearity in the predictors.

Second, QR is generally more robust than ordinary least-squared regressions, in which the absolute value of an outlier can strongly influence the average response of the dependent variable. When modelling a quantile, the absolute value of an outlier has less impact on the shape of the model (Scharf et al., 1998). If many absences occurring along the environmental gradient are susceptible to pull down a model of the mean, these absences will have a moderated impact on a quantile model (Cade and Noon, 2003; Schröder et al., 2005).

### 4 APPENDIX 4: SEDIMENT

Table A.4 : Table of number of sediment for each sediment category

| Sediment id | Sediment category | nb samples (% of total number of samples) |
|---|---|---|
| 1 | Coarse sediment | 122 (3.2%) |
| 2 | Compacted sediments or sedimentary bedrock | 1 (0.026%) |
| 3 | Mixed sediment | 829 (22%) |
| 4 | Mud, clay and sandy mud | 2183 (57%) |
| 5 | Sand and muddy sand | 388 (10%) |
| 6 | Sand, gravel and pebbles | 22 (0.57%) |
| 7 | Thin/discont. sedim. cover on bedrock | 5 (0.13%) |
| 8 | No sediment data available | 277 (7.2%) |

### 5 APPENDIX 5: INDIVIDUAL SPECIES FITTED QGAM MODELS

All fitted QGAM models on the training dataset are available in the pdf "Annex5_qgam_models.pdf". Each page corresponds to one of the 33 species.

Caption: Modelled log10 responses to the 10 selected environmental predictors. A and B: black dotted scatterplot of the log of positive biomasses of the species in response to the predictor. Red dots indicate model maximum biomass predictions. On top of the scatterplot, the marginal density shows the distribution of samples conditional to the predictor values. C: Boxplot of response to the sediment. The model prediction is the $99^{th}$ quantile for each sediment class: 1= Coarse sediment, 2= Compacted sediment or sedimentary bedrock, 3= Mixed sediment, 4= Mud, clay and sandy mud, 5= Sand and muddy sand, 6= Sand, gravel and pebbles, 7= Thin or discontinuous sediment on bedrock

### 6 APPENDIX 6: INDIVIDUAL SPECIES NICHE DESCRIPTION

Description of all the species niche along the 10 selected environmental gradients are available in pdf "Annex6_species_niche_desc.pdf". There is one page per parameter

Figure explanation: Species habitat preference and tolerance. The mode is the maximum of the modelled response to the predictor. The range is calculated as the ratio of the difference between the range (max – min) of predictor values where the species has been found to the range of all the Barents Sea. Species tolerant regarding the predictor are situated in the top of the plot, while species with a narrower niche are in the bottom. The dashed vertical line indicates the mean value of the parameter in the Barents Sea.

Table explanation: Species niche and model description. Species: Latin names of the species. Code: abbreviated names of the species. Min and Max: minimal and maximum values where the species has been observed. Range: ratio of Max-Min of the species occurrences to Max-Min of the whole Barents Sea. BS mean: mean value of the predictor in the Barents Sea. Sp mean: mean value of the predictor linked to the species occurences. Contrast: contrast in the QGAM modelled response of the species to the predictor. Rows of the tables are organized by increasing contrast. Data under the model: % of data from the testing data set that are below the modelled response (should be close to 99.0%)

### 7 APPENDIX 7: INDIVIDUAL SPECIES HABITAT SUITABILITY MAPS

Habitat suitability and most limiting factors maps are available in pdf "Annex7_hs_maps.pdf"

Figure explanation: *Spatial predictions in 2013 of the species A) suitable habitat (maximum biomass) and B) most limiting predictor. Color indicates the predictor's category: fixed (sediment, depth, slope), dynamic (all the others). Grey symbols indicate that the predictor is not very limiting (predicted biomass > 25% of the model maximum)*
