## Appendix 5 QGAM models for "Suitable habitats of fish species in the Barents Sea"

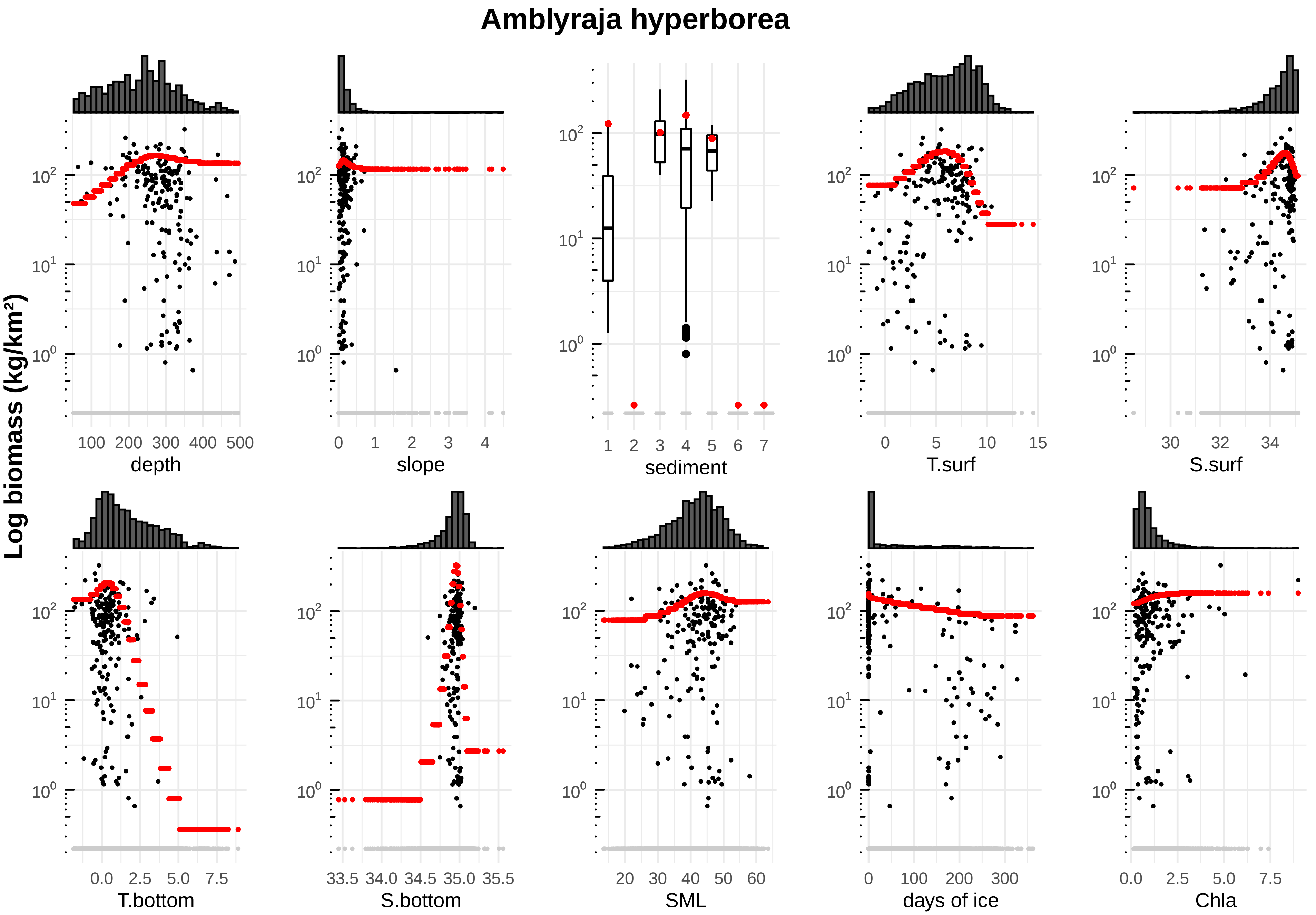

Amblyraja hyperborea
Log biomass (kg/km²)
depth
100 200 300 400 500
slope
0 1 2 3 4
sediment
1 2 3 4 5 6 7
T.surf
0 5 10 15
S.surf
30 32 34
T.bottom
0.0 2.5 5.0 7.5
S.bottom
33.5 34.0 34.5 35.0 35.5
SML
20 30 40 50 60
days of ice
0 100 200 300
Chla
0.0 2.5 5.0 7.5

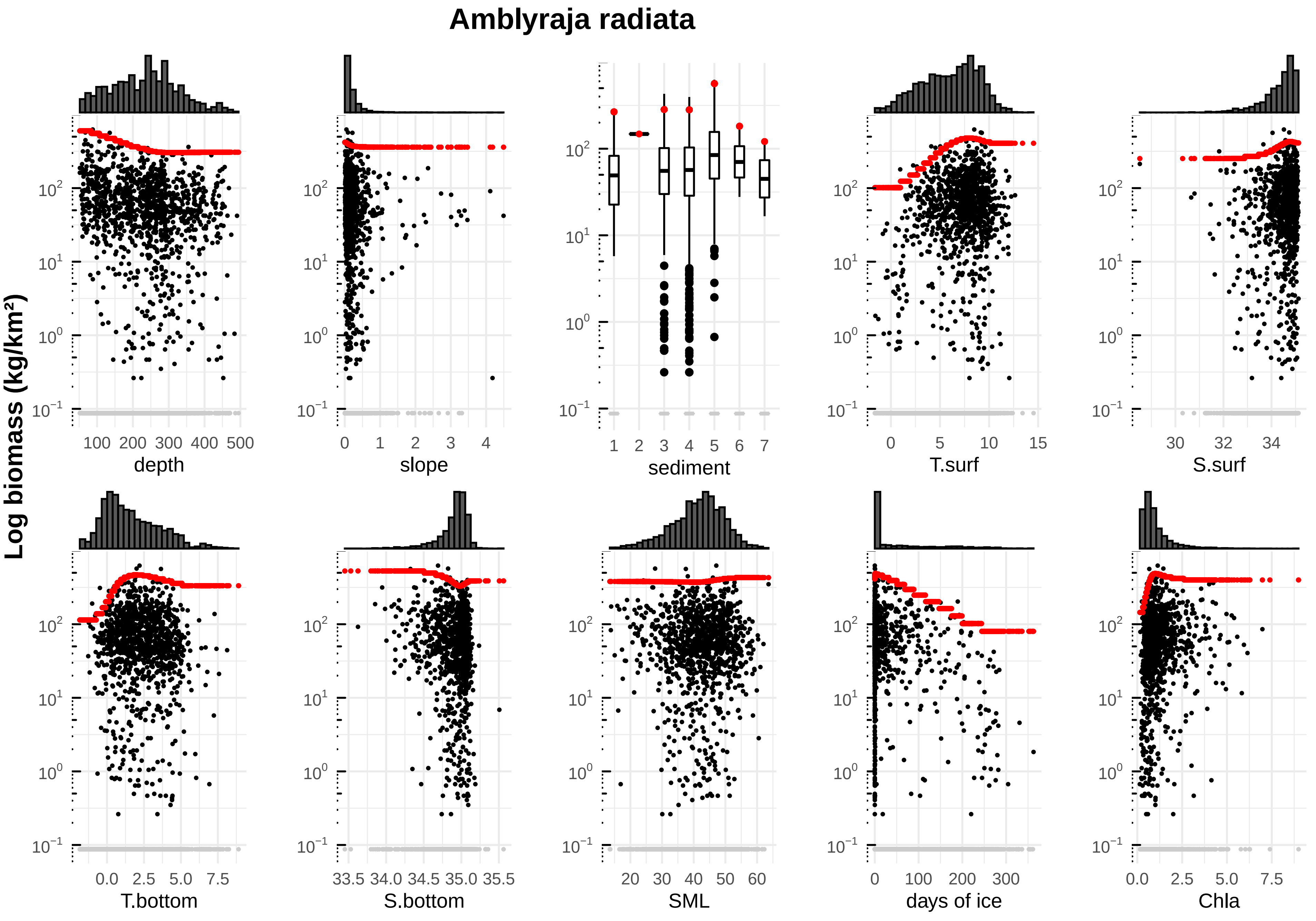
Amblyraja radiata
Log biomass (kg/km²)
depth
slope
sediment
T.surf
S.surf
T.bottom
S.bottom
SML
days of ice
Chla

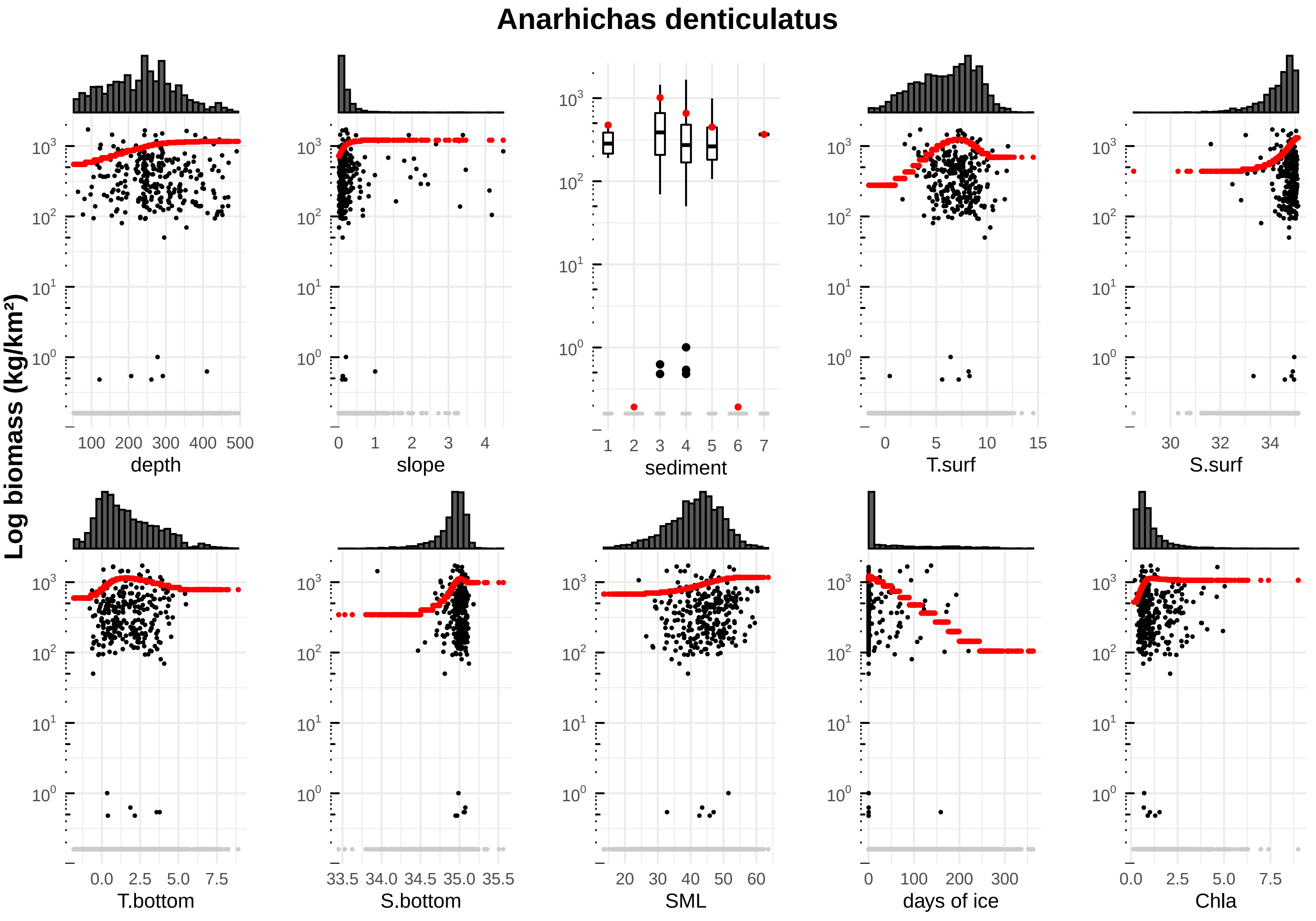

Anarhichas denticulatus
Log biomass (kg/km²)
depth
slope
sediment
T.surf
S.surf
T.bottom
S.bottom
SML
days of ice
Chla

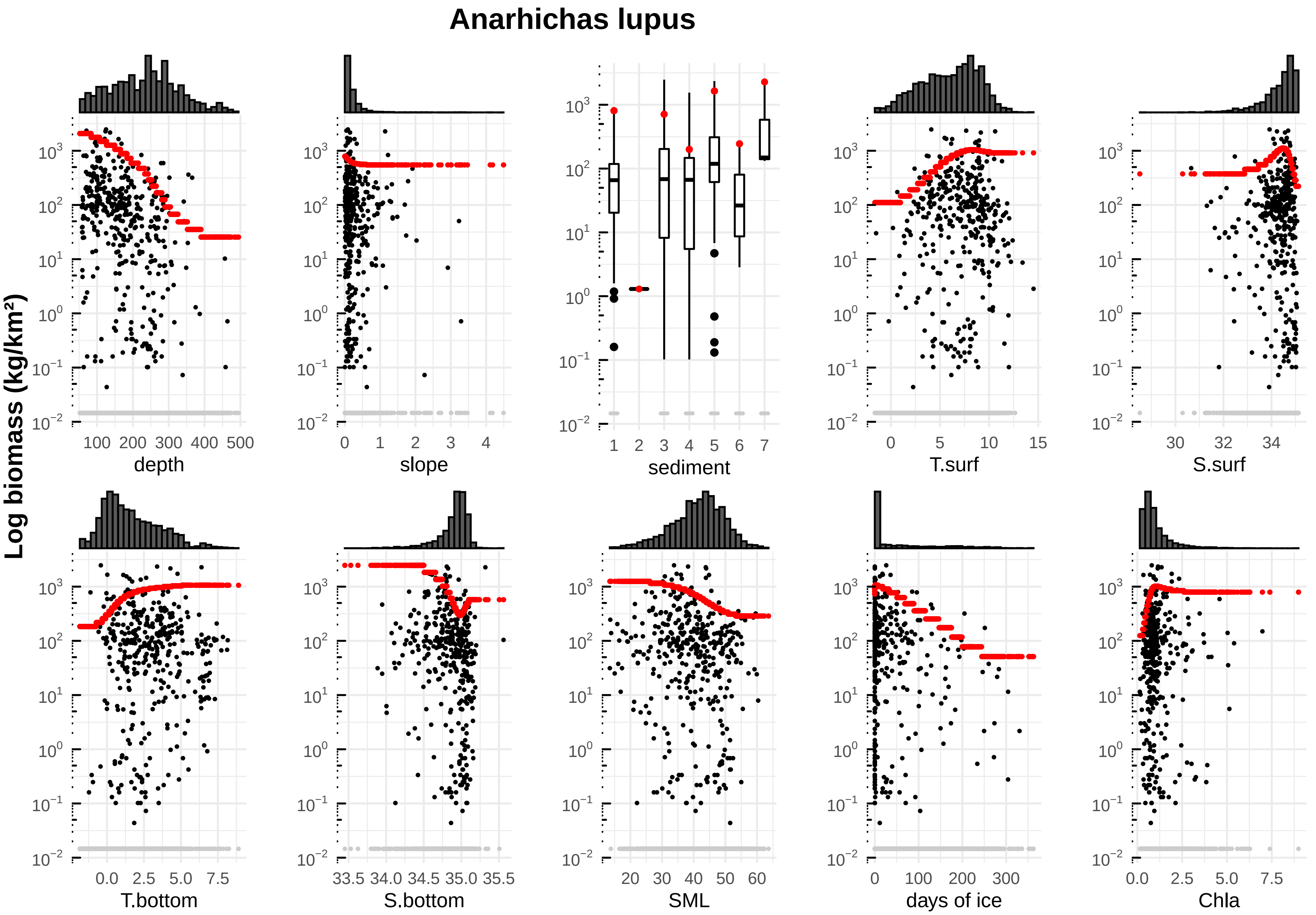

Anarhichas lupus
Log biomass (kg/km²)
depth
slope
sediment
T.surf
S.surf
T.bottom
S.bottom
SML
days of ice
Chla

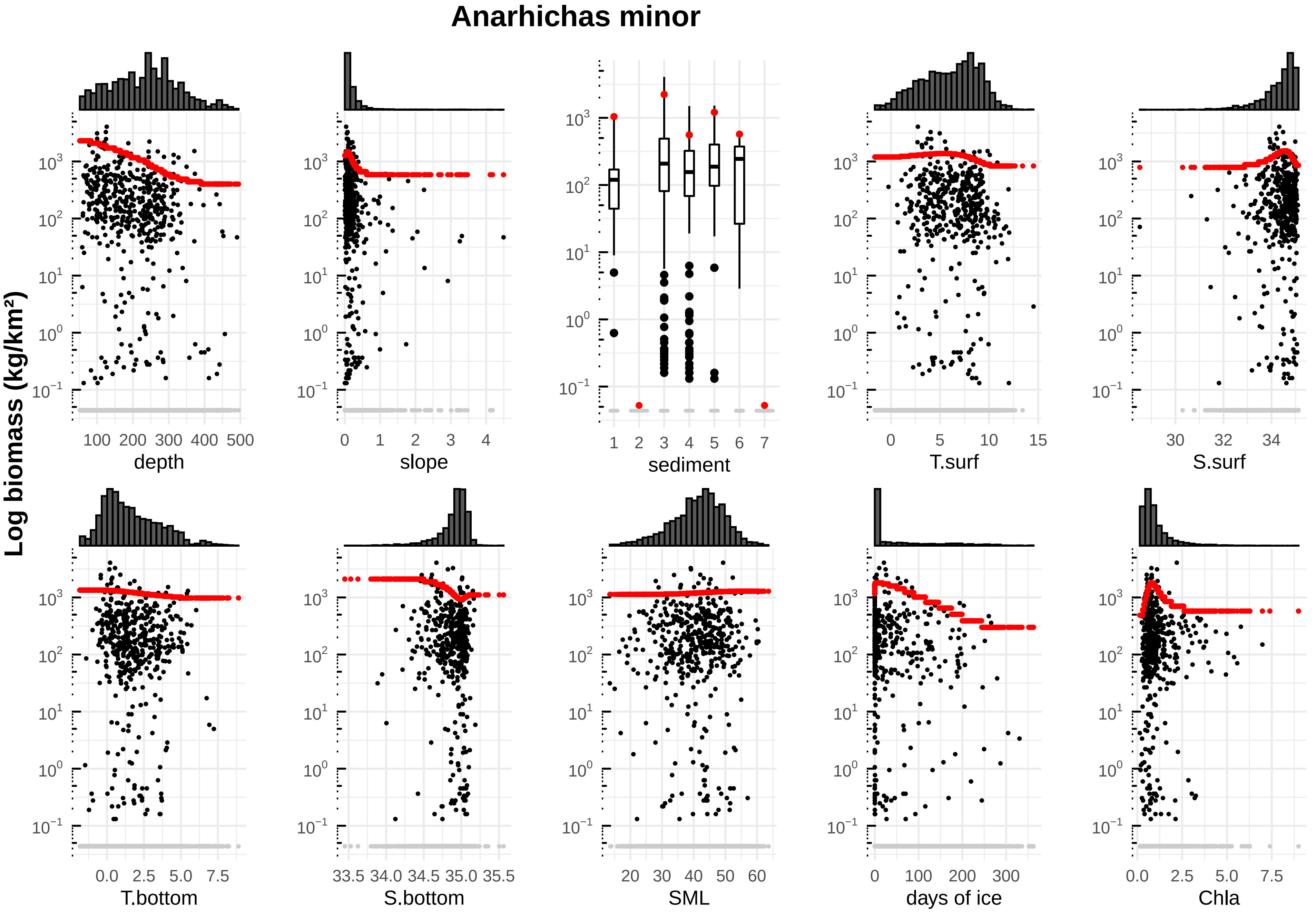

Anarhichas minor
Log biomass (kg/km²)
depth
slope
sediment
T.surf
S.surf
T.bottom
S.bottom
SML
days of ice
Chla

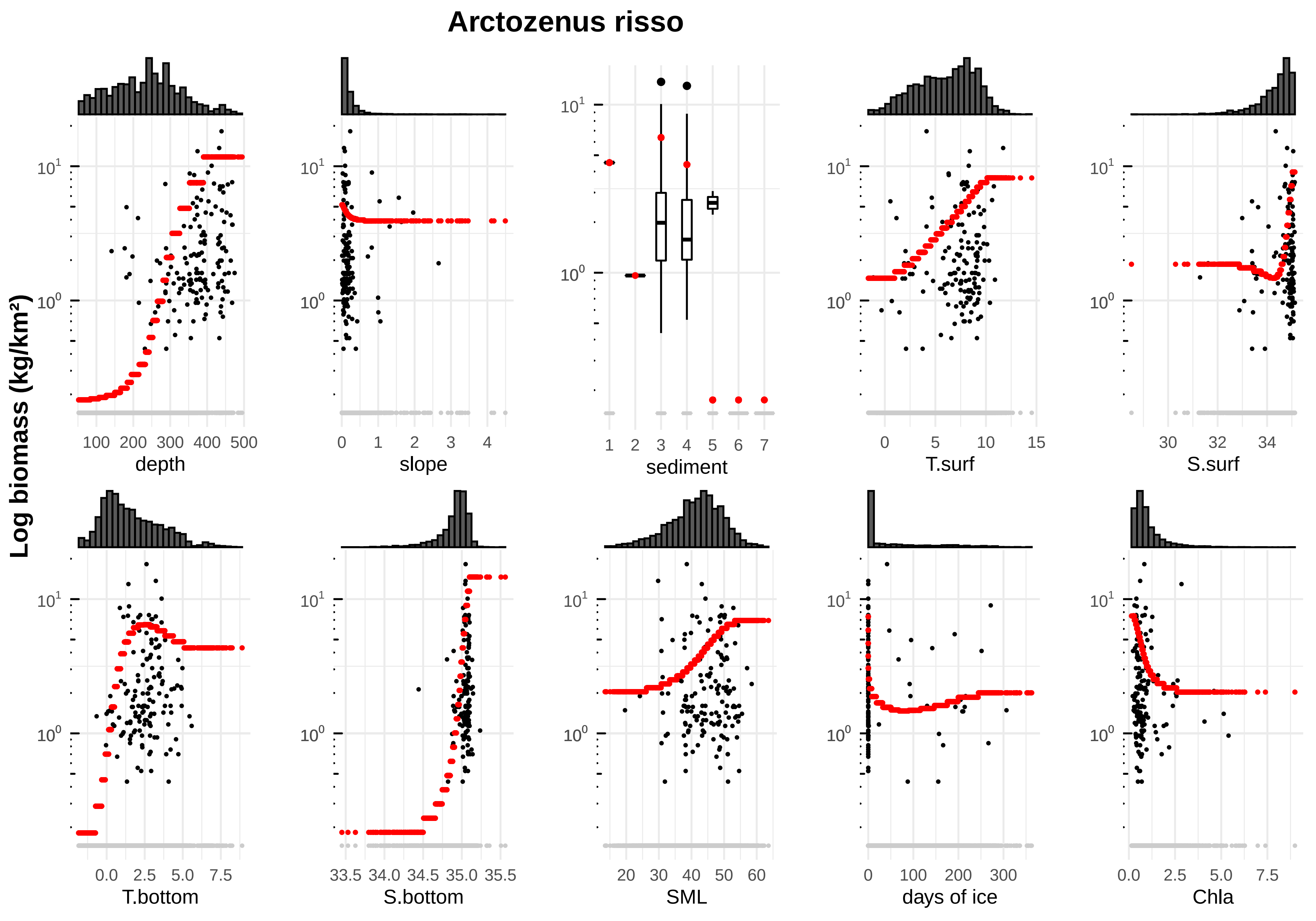
Arctozenus risso
Log biomass (kg/km²)
depth
slope
sediment
T.surf
S.surf
T.bottom
S.bottom
SML
days of ice
Chla

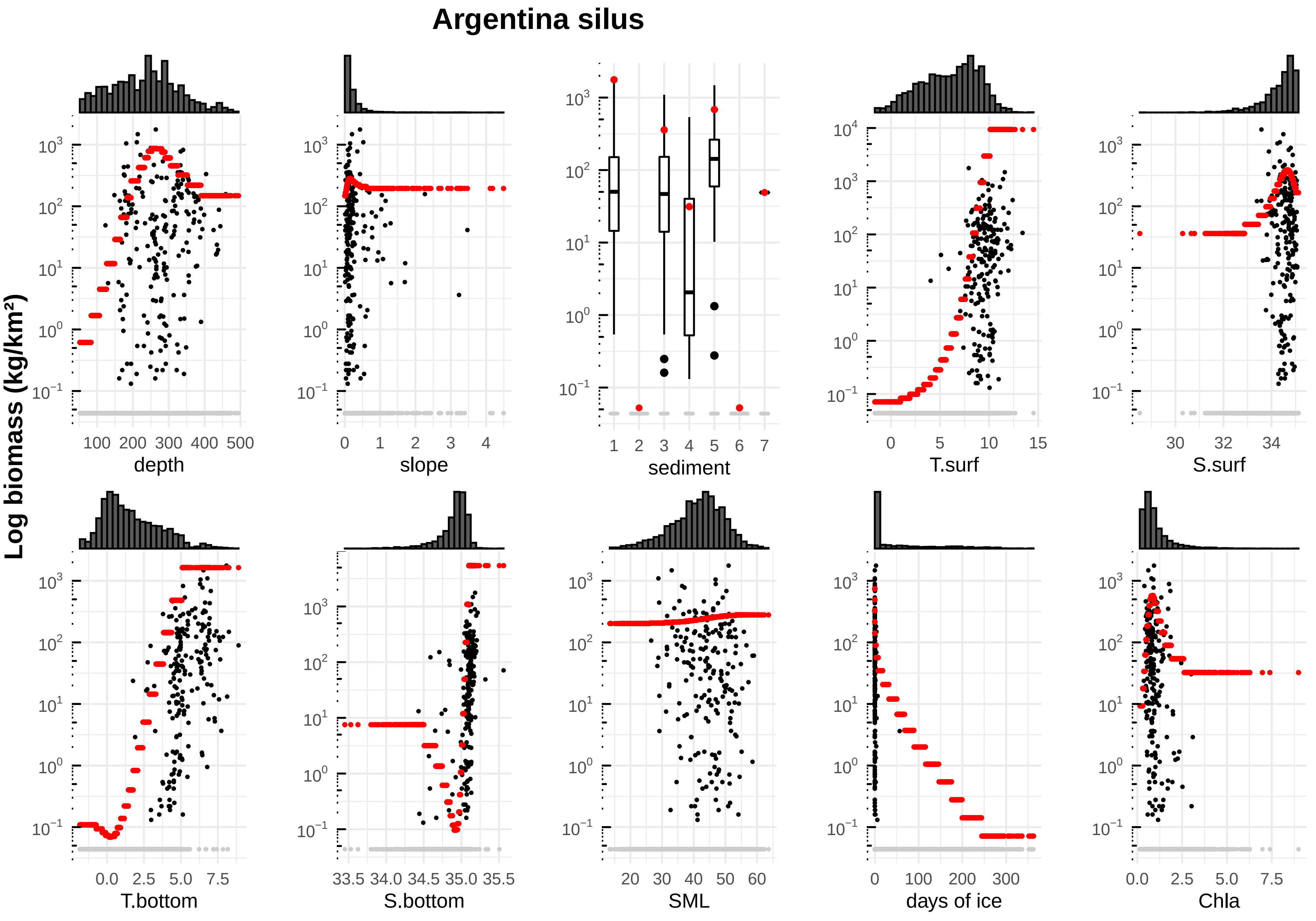

Argentina silus
Log biomass (kg/km²)
depth
slope
sediment
T.surf
S.surf
T.bottom
S.bottom
SML
days of ice
Chla

**Artediellus atlanticus**

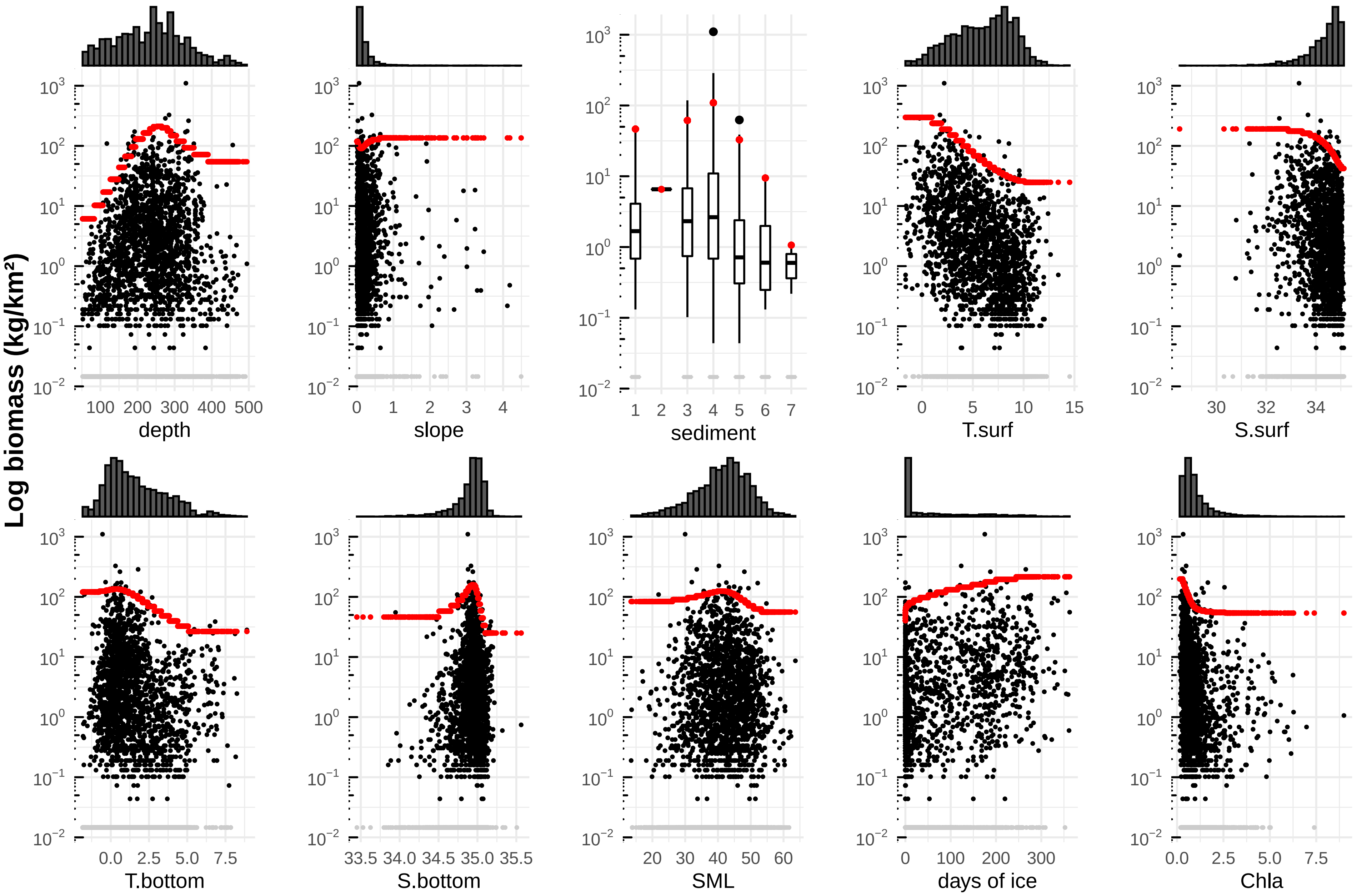

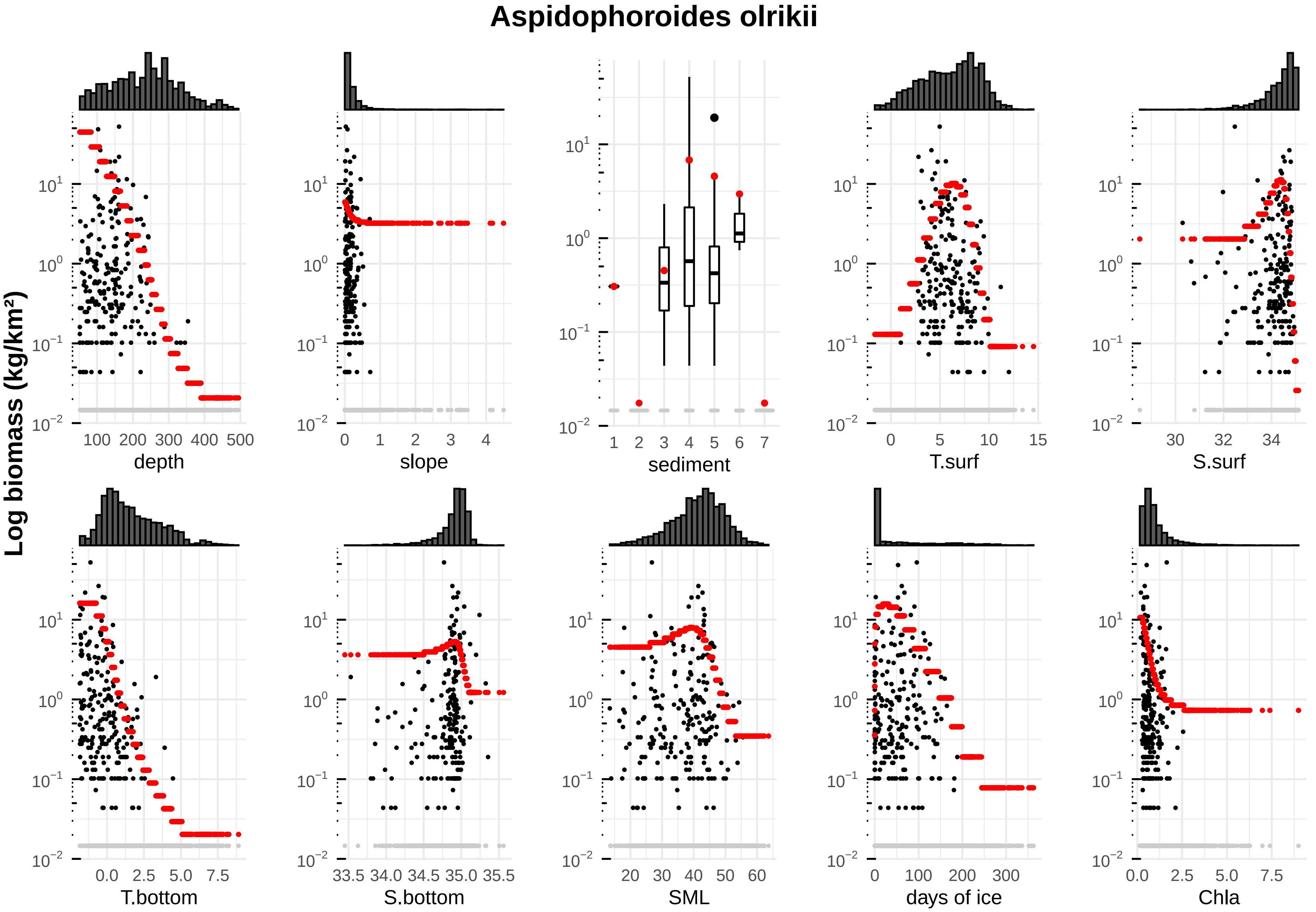

Aspidophoroides olrikii
Log biomass (kg/km²)
depth
slope
sediment
T.surf
S.surf
T.bottom
S.bottom
SML
days of ice
Chla

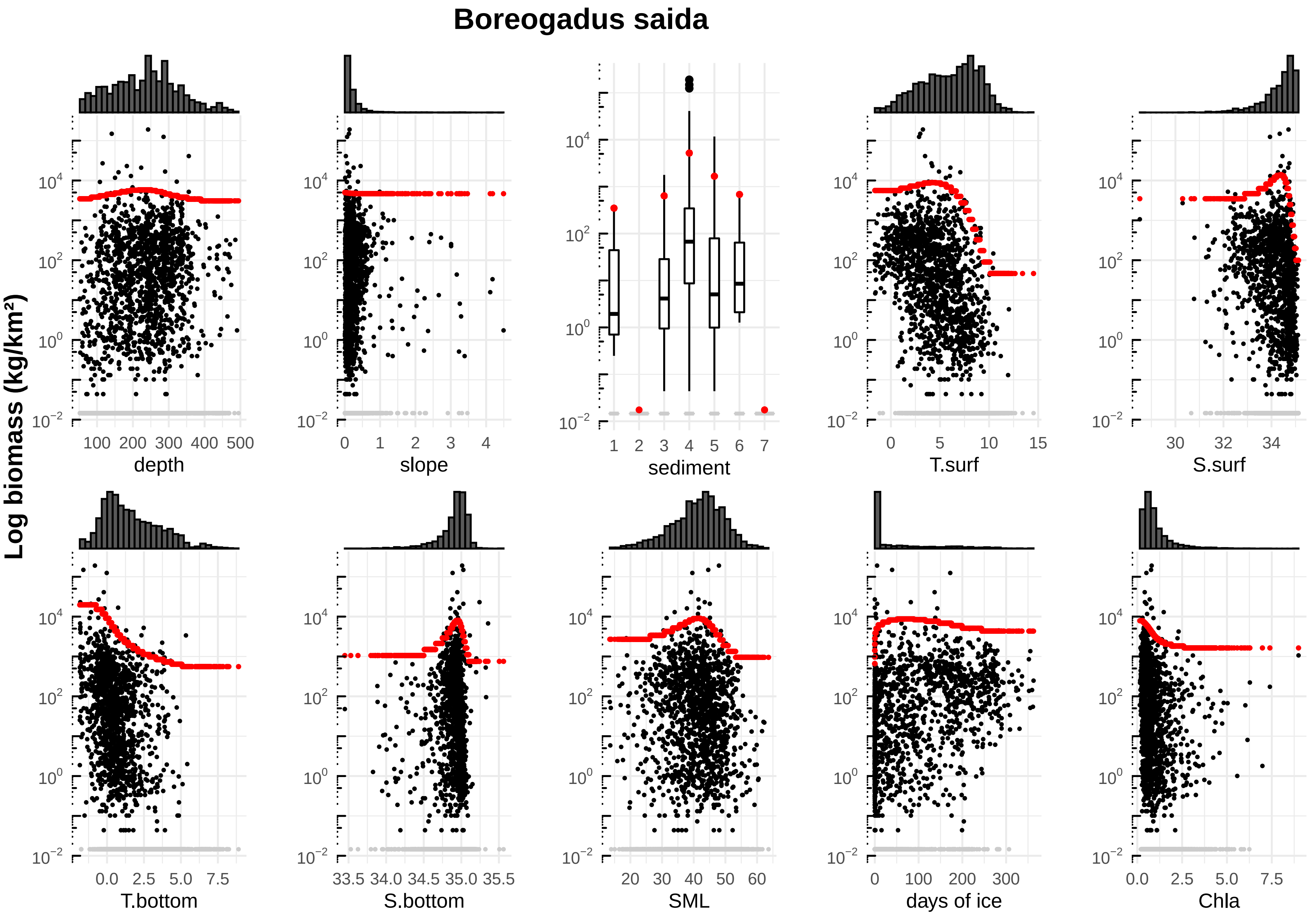

Boreogadus saida
Log biomass (kg/km²)
depth
slope
sediment
T.surf
S.surf
T.bottom
S.bottom
SML
days of ice
Chla

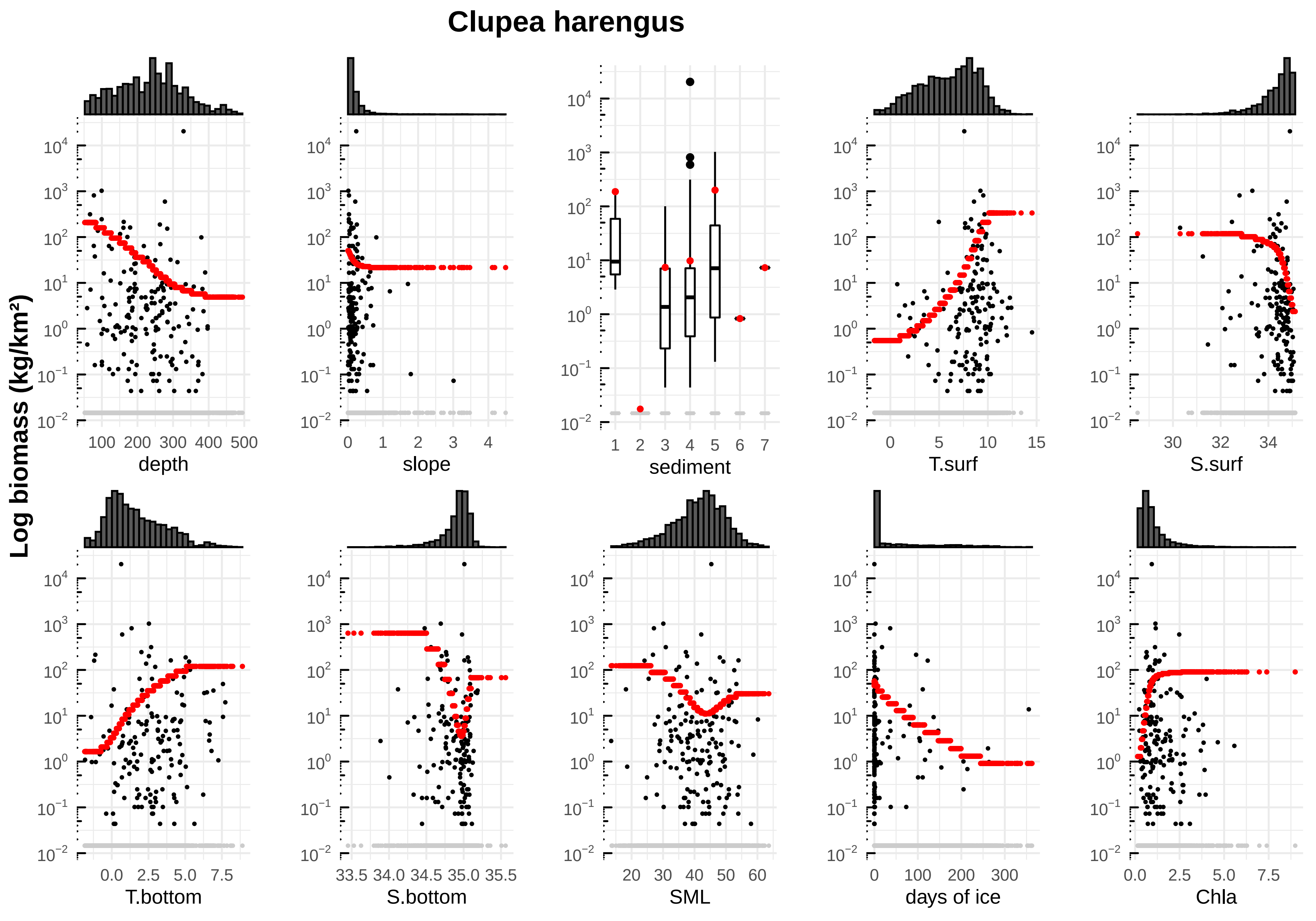

Clupea harengus
Log biomass (kg/km²)
depth
slope
sediment
T.surf
S.surf
T.bottom
S.bottom
SML
days of ice
Chla

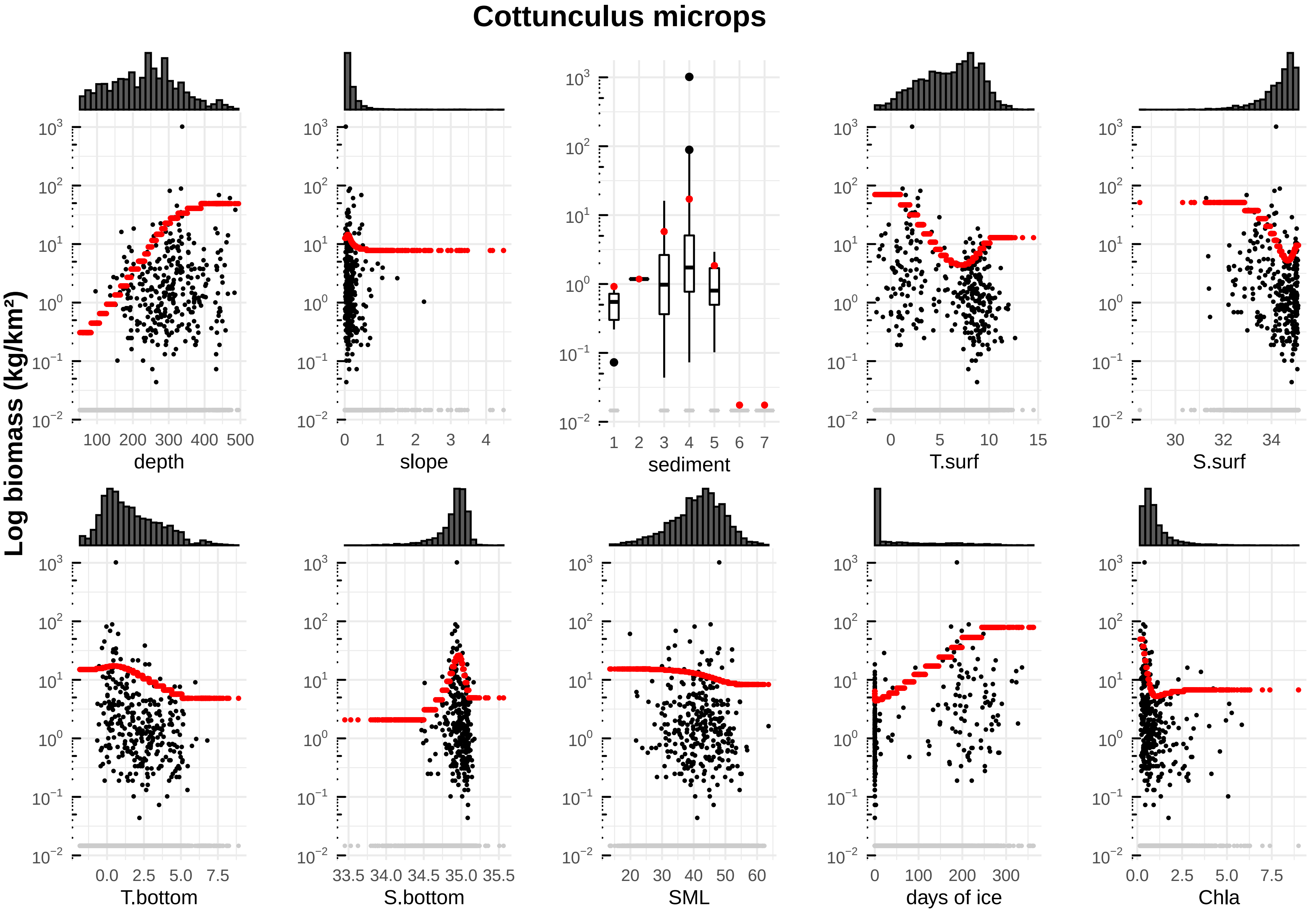

Cottunculus microps
Log biomass (kg/km²)
depth
slope
sediment
T.surf
S.surf
T.bottom
S.bottom
SML
days of ice
Chla

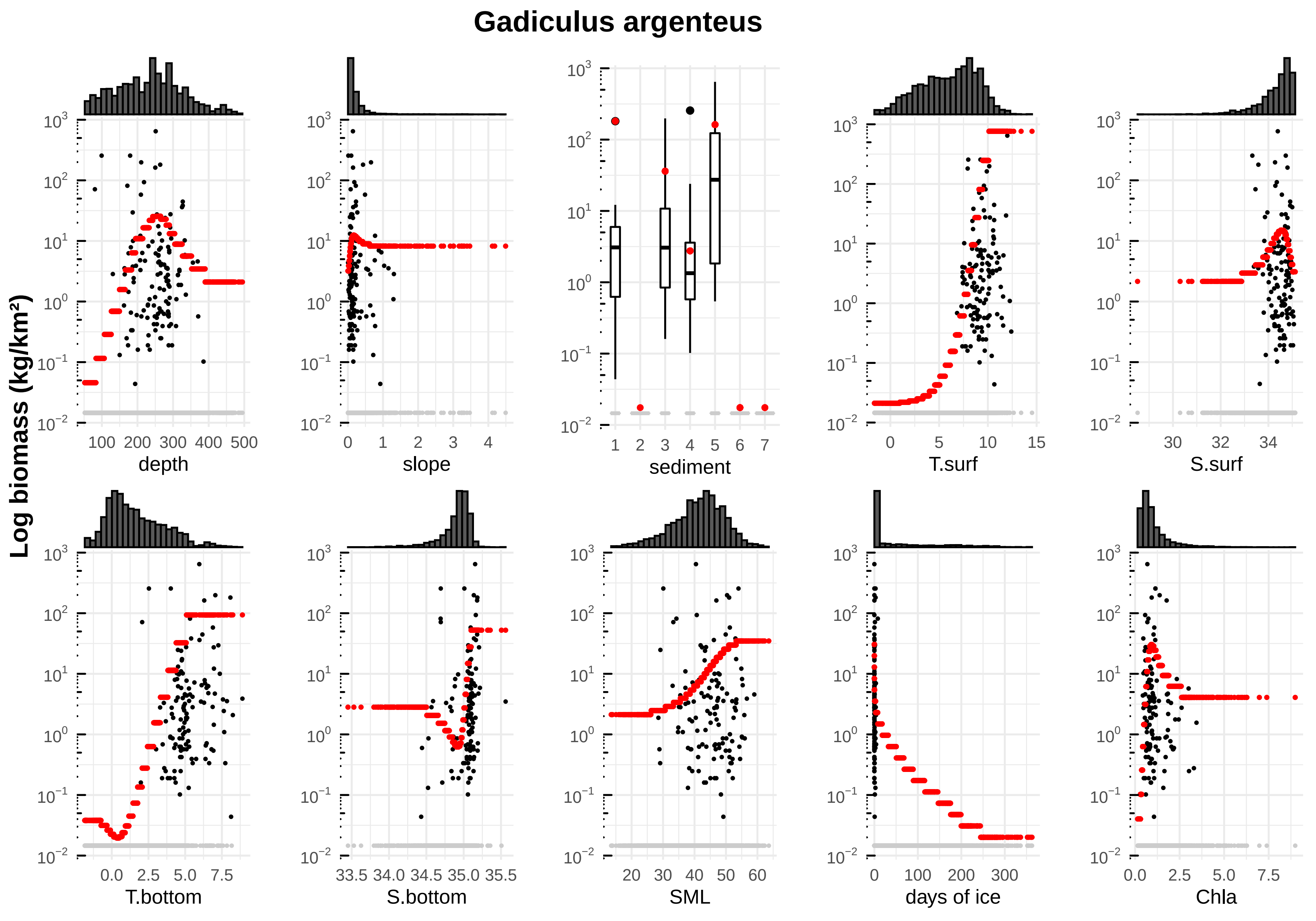

Gadiculus argenteus
Log biomass (kg/km²)
depth
slope
sediment
T.surf
S.surf
T.bottom
S.bottom
SML
days of ice
Chla

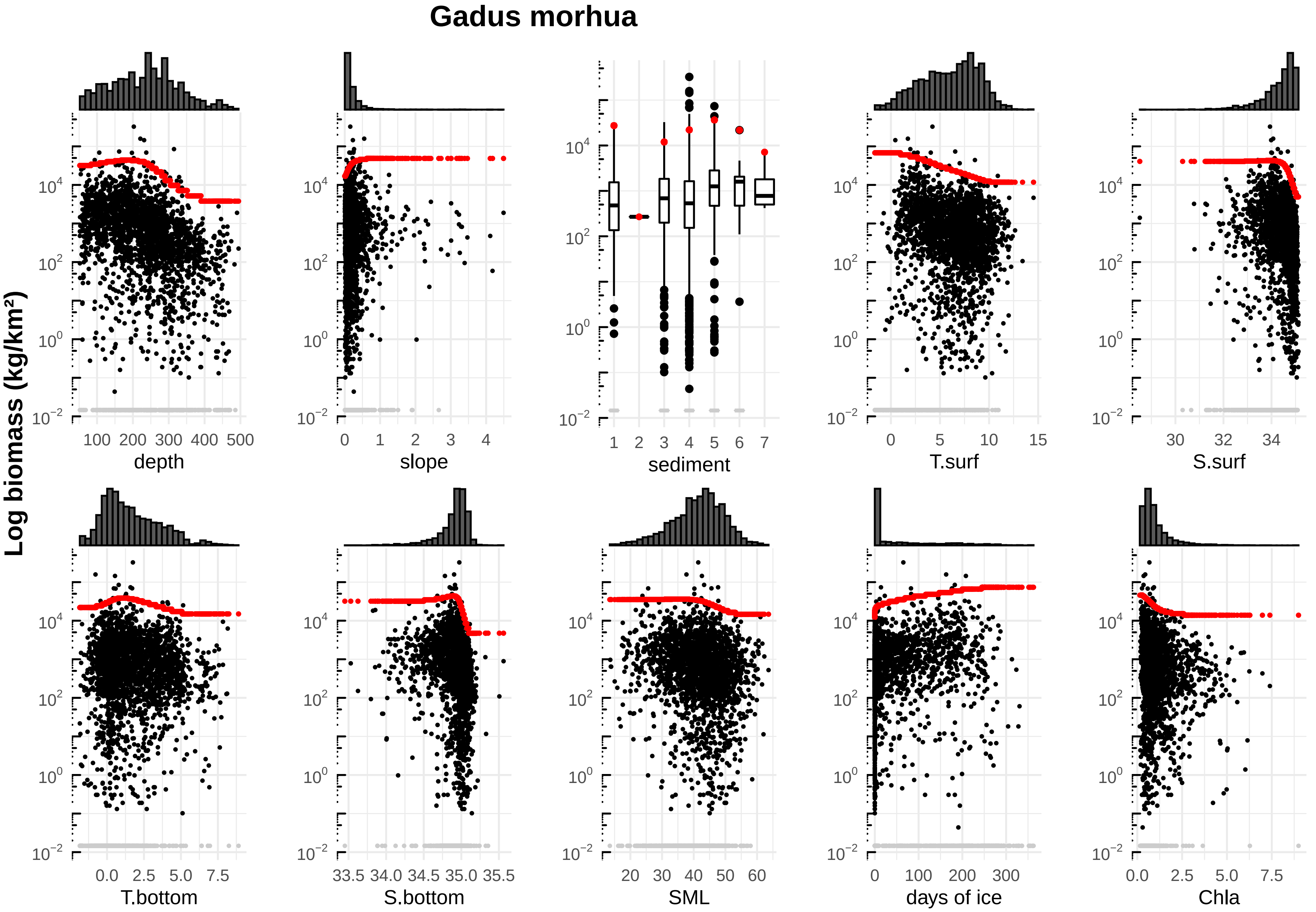

Gadus morhua
Log biomass (kg/km²)
depth
slope
sediment
T.surf
S.surf
T.bottom
S.bottom
SML
days of ice
Chla

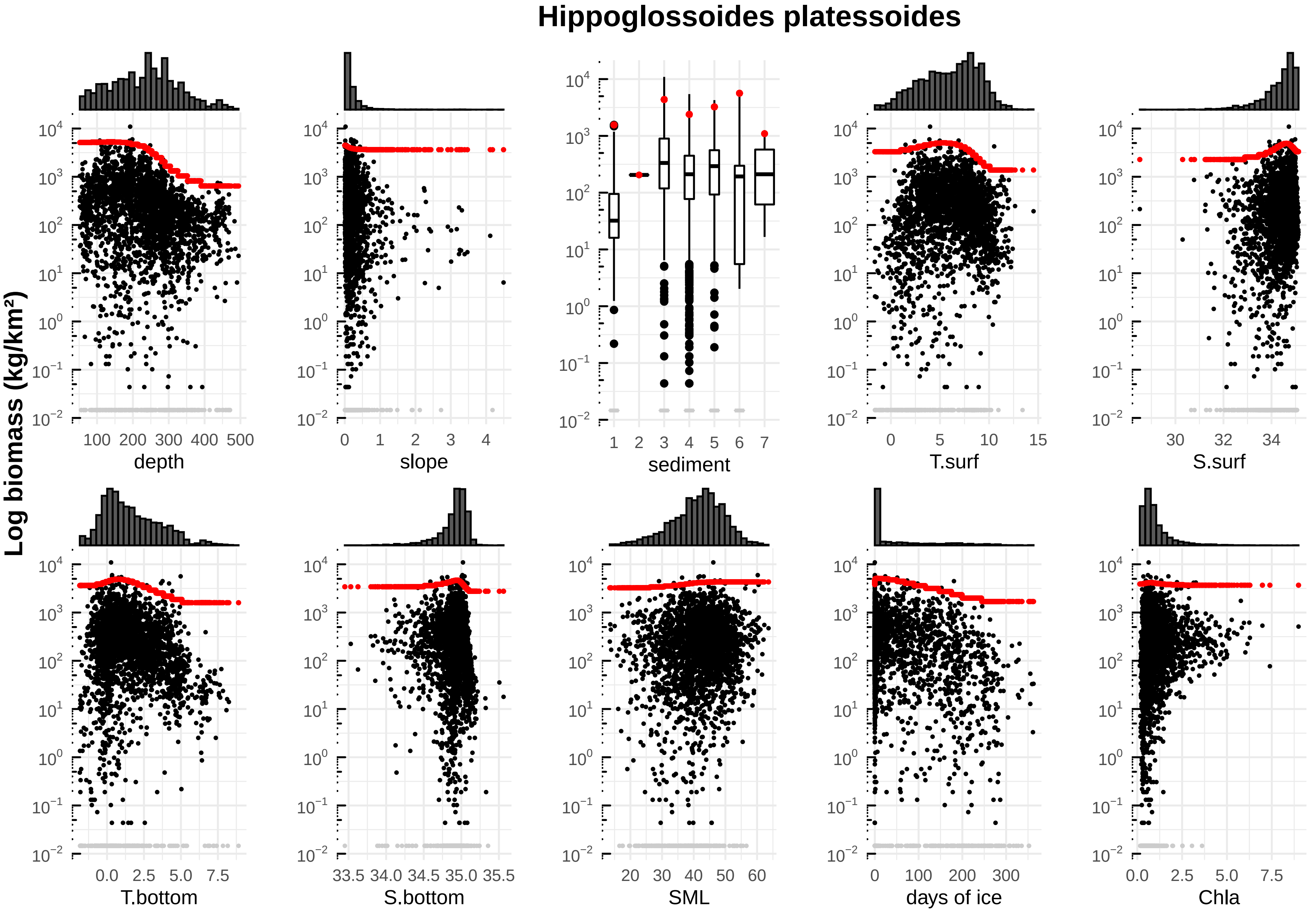

Hippoglossoides platessoides
Log biomass (kg/km²)
depth
slope
sediment
T.surf
S.surf
T.bottom
S.bottom
SML
days of ice
Chla

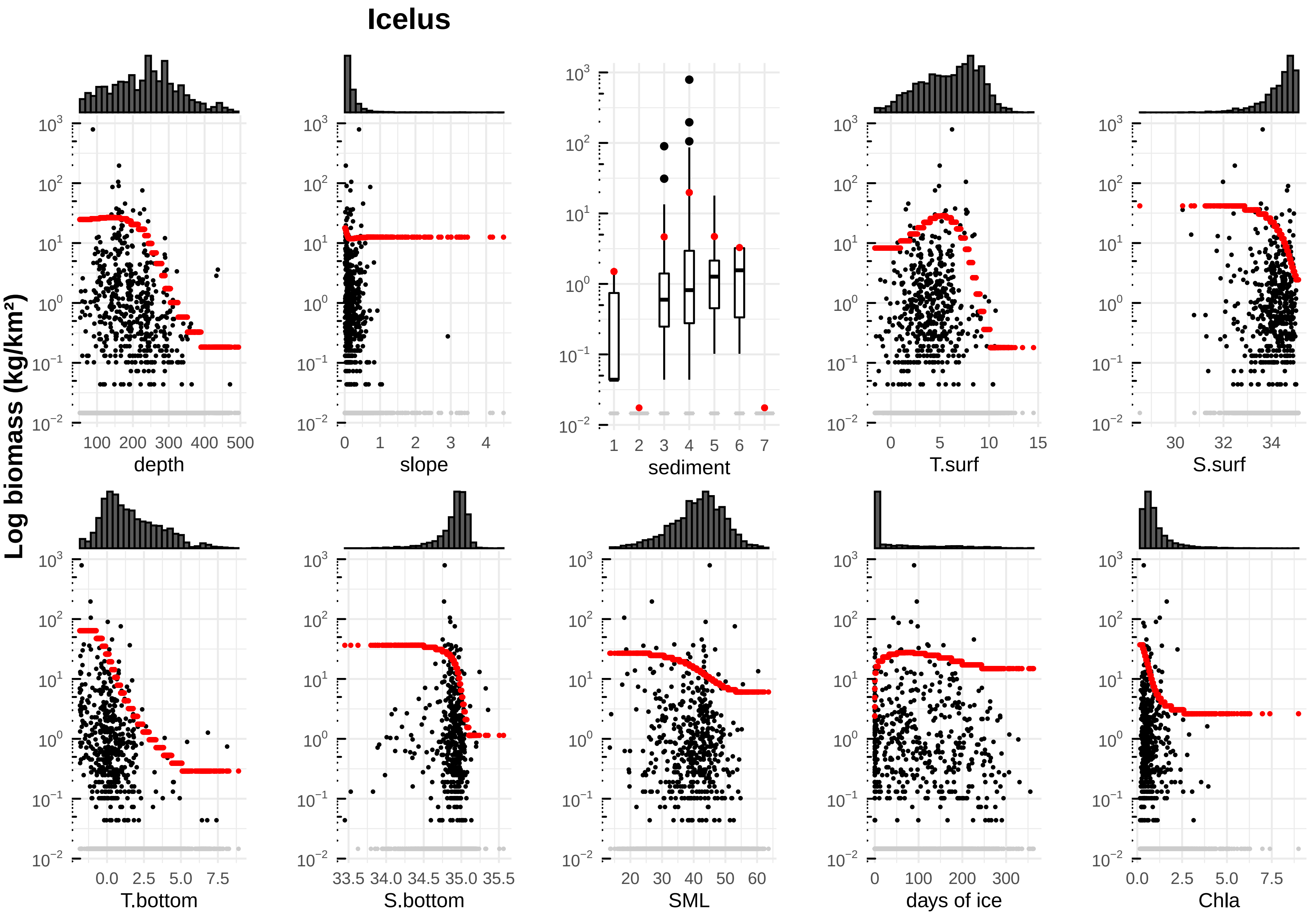

Icelus
Log biomass (kg/km²)
depth
slope
sediment
T.surf
S.surf
T.bottom
S.bottom
SML
days of ice
Chla

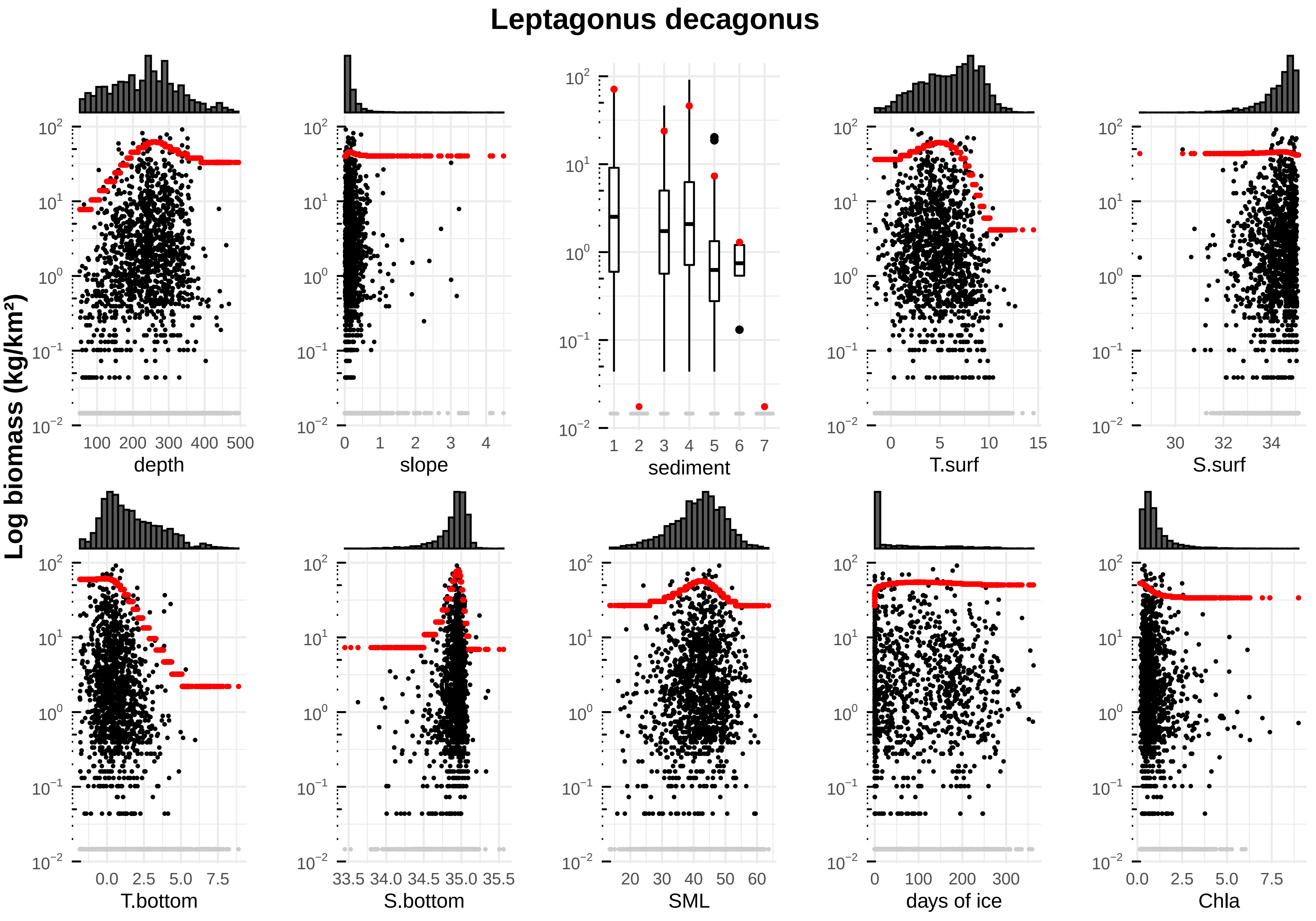

Leptagonus decagonus
Log biomass (kg/km²)
depth
slope
sediment
T.surf
S.surf
T.bottom
S.bottom
SML
days of ice
Chla

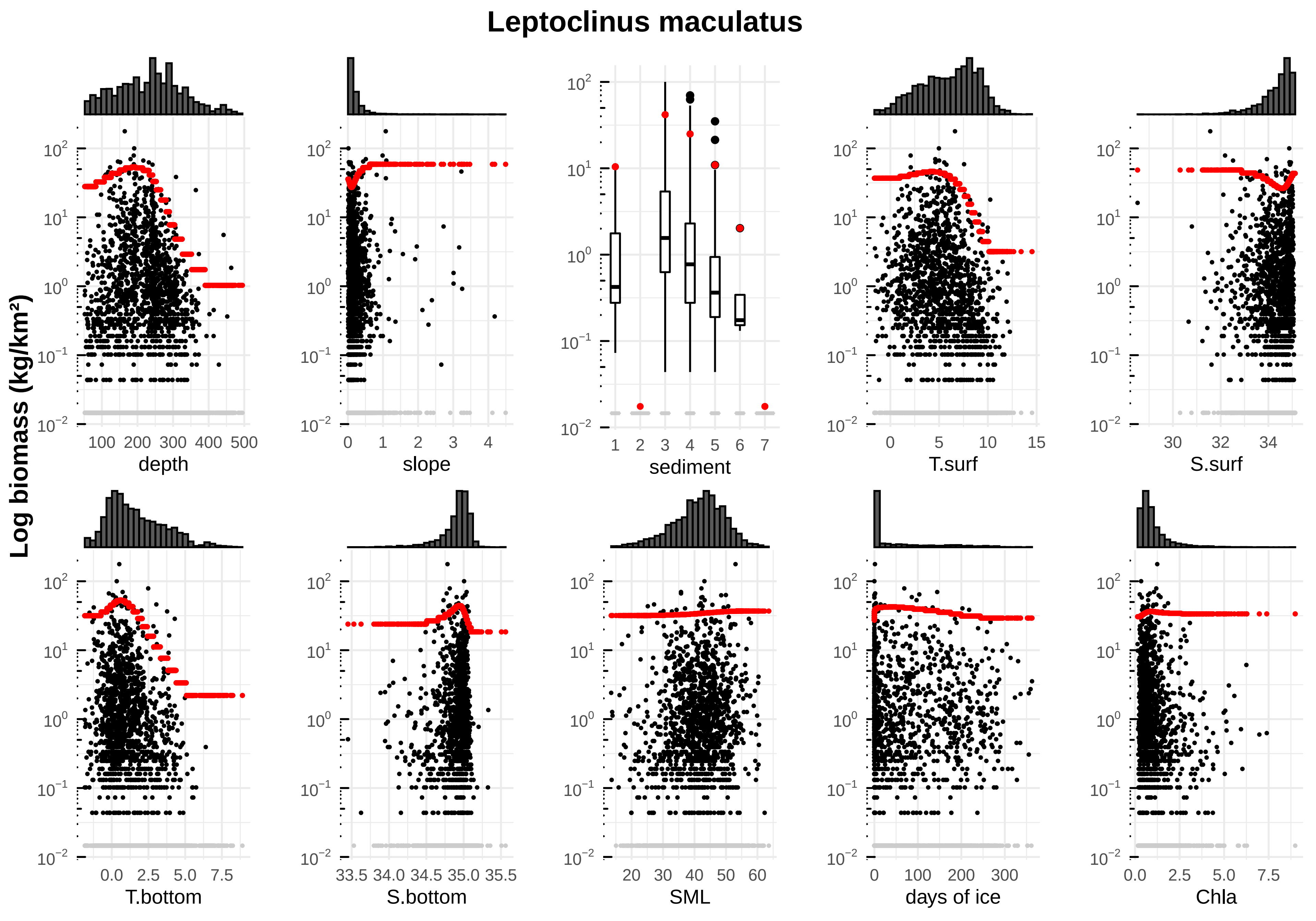

Leptoclinus maculatus
Log biomass (kg/km²)
depth
slope
sediment
T.surf
S.surf
T.bottom
S.bottom
SML
days of ice
Chla

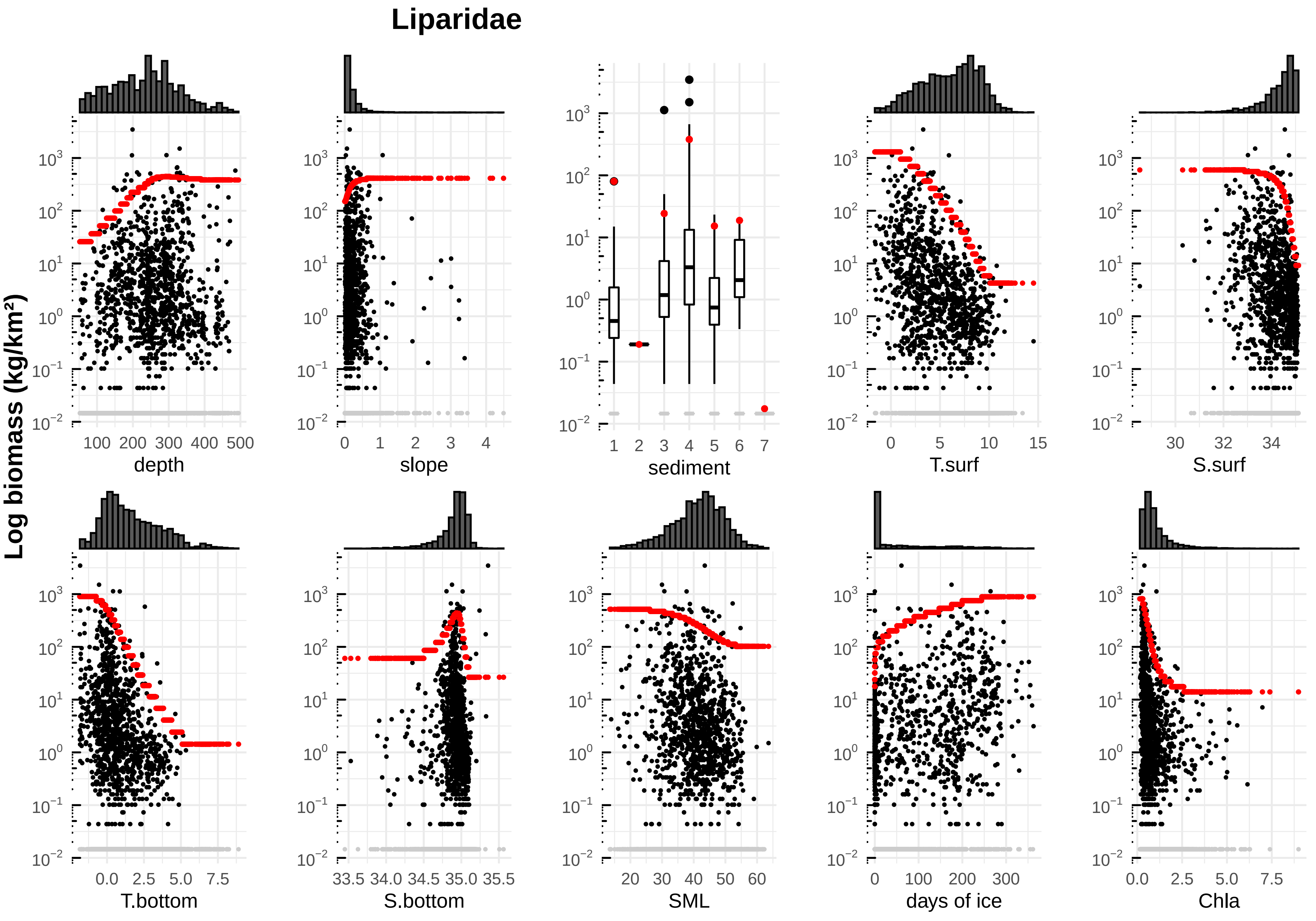

Liparidae
Log biomass (kg/km²)
depth
slope
sediment
T.surf
S.surf
T.bottom
S.bottom
SML
days of ice
Chla

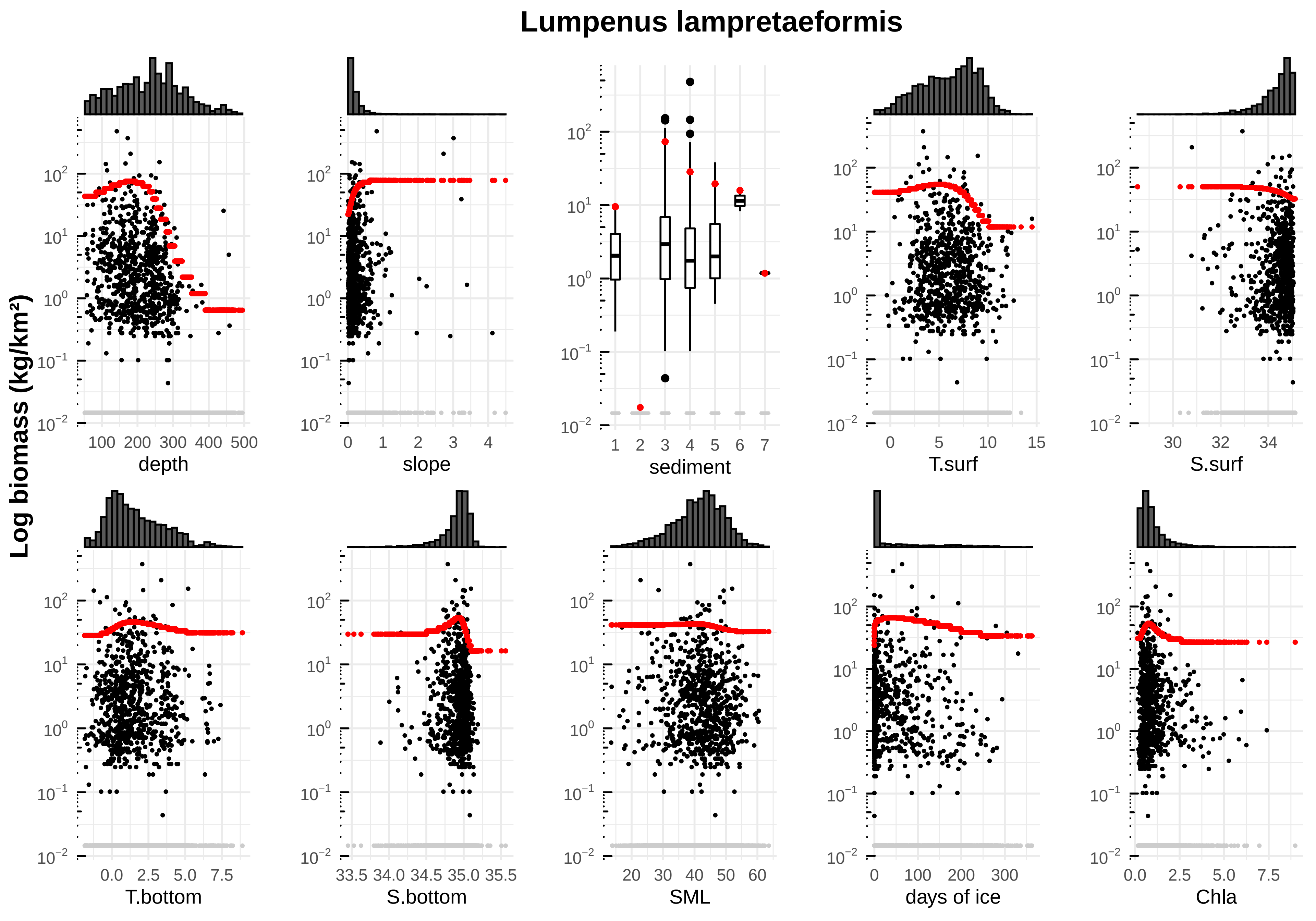

Lumpenus lampretaeformis
Log biomass (kg/km²)
depth
slope
sediment
T.surf
S.surf
T.bottom
S.bottom
SML
days of ice
Chla

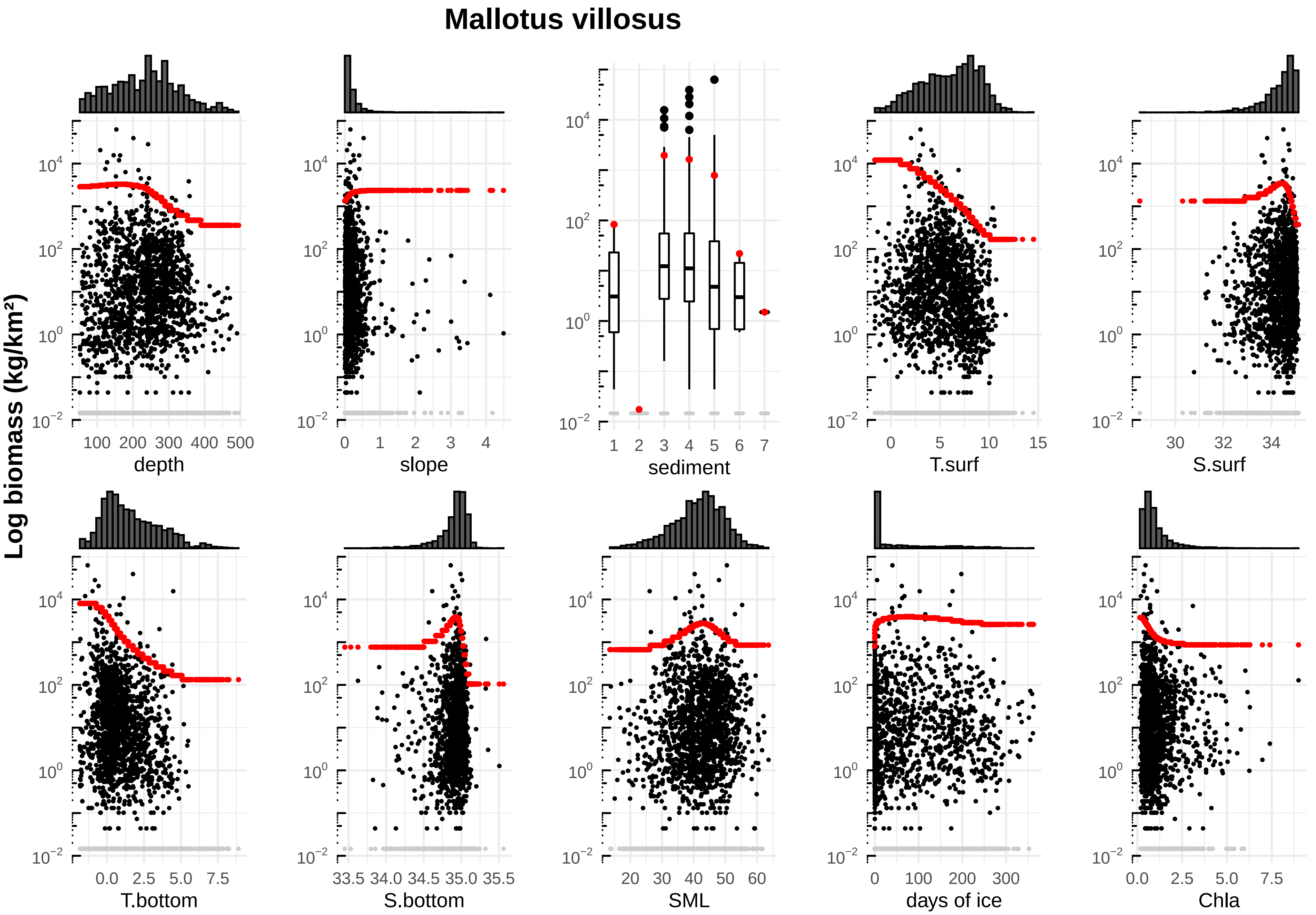

Mallotus villosus
Log biomass (kg/km²)
depth
slope
sediment
T.surf
S.surf
T.bottom
S.bottom
SML
days of ice
Chla

#### Melanogrammus aeglefinus

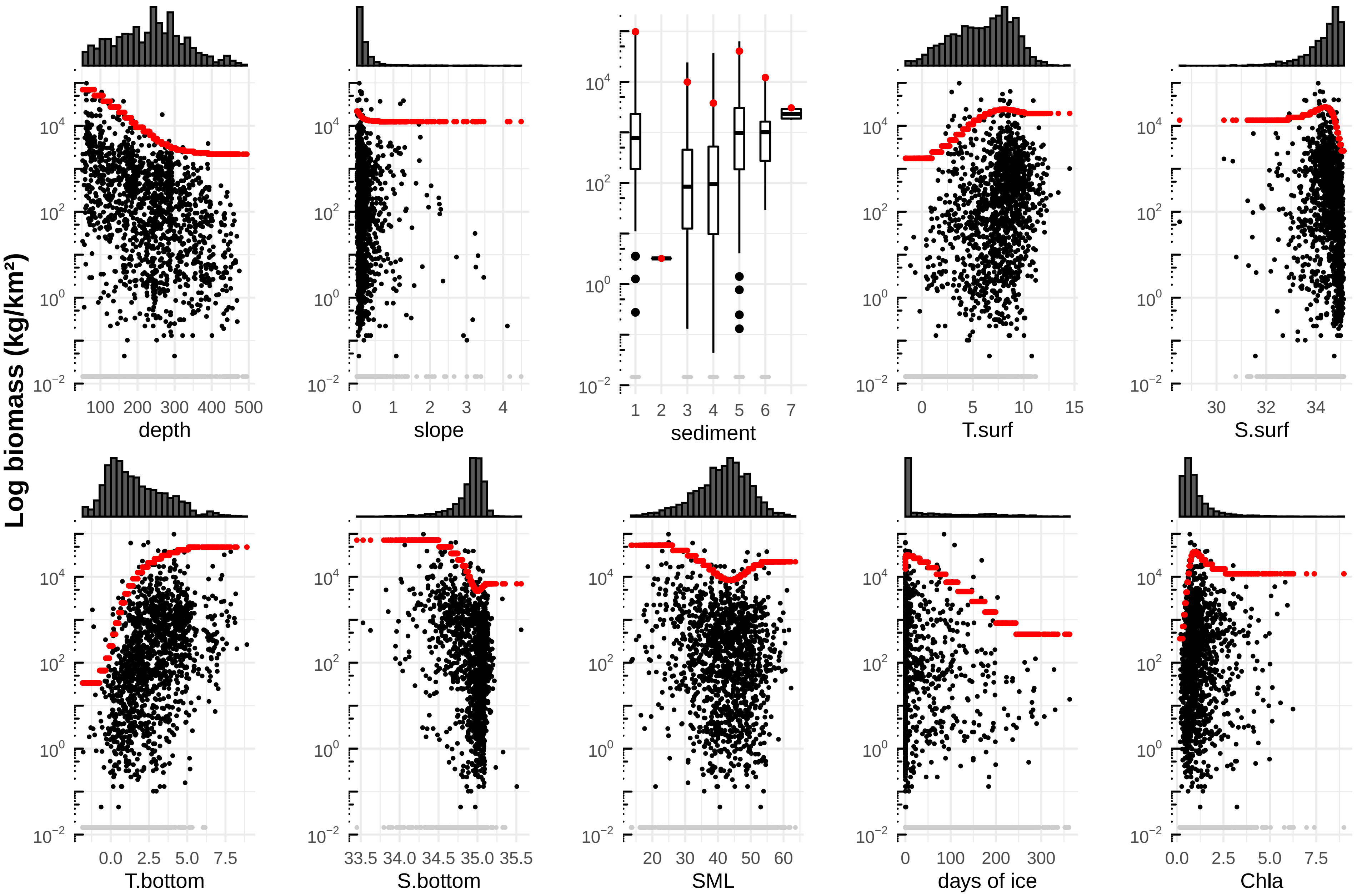

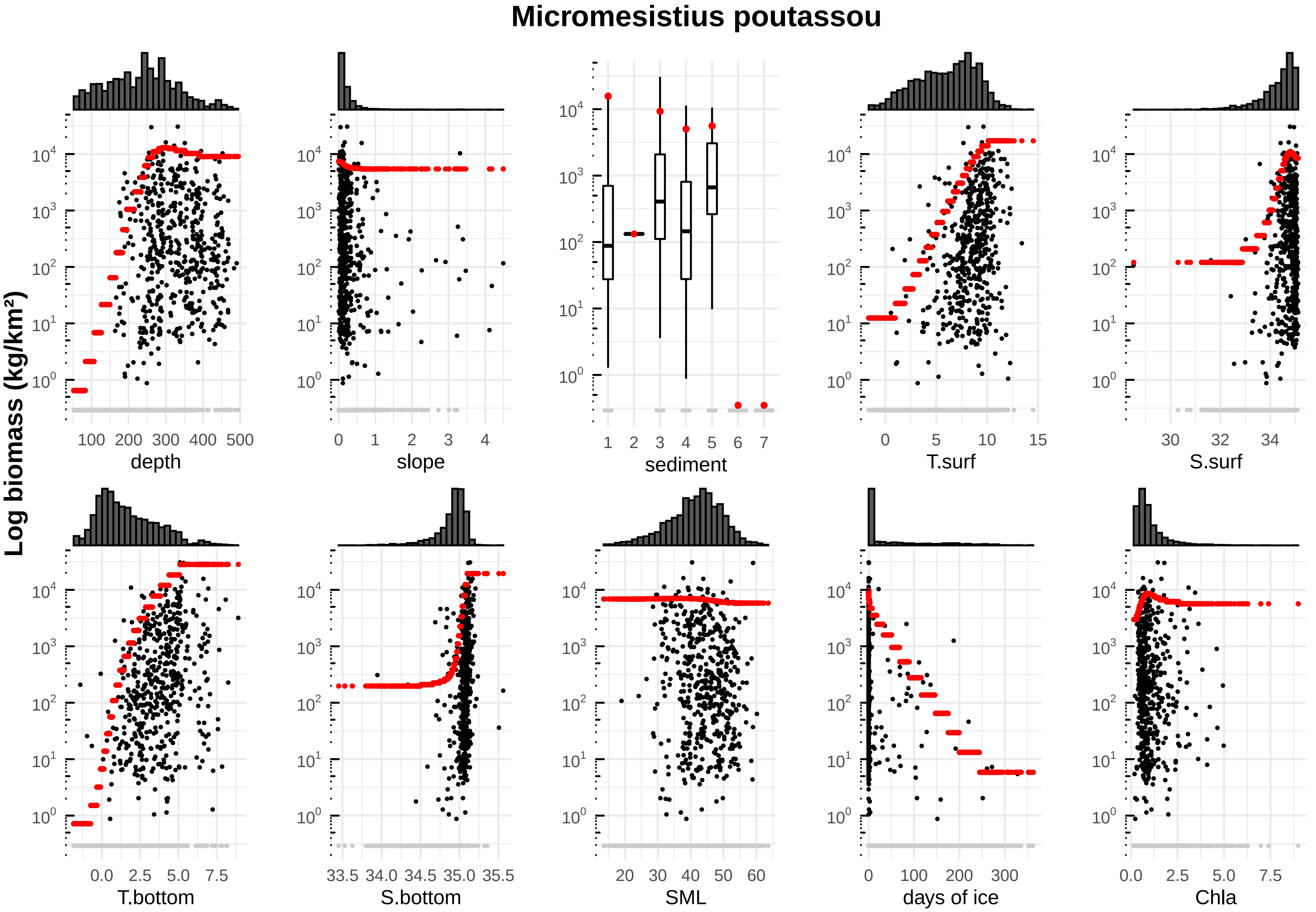

Micromesistius poutassou
Log biomass (kg/km²)
depth
slope
sediment
T.surf
S.surf
T.bottom
S.bottom
SML
days of ice
Chla

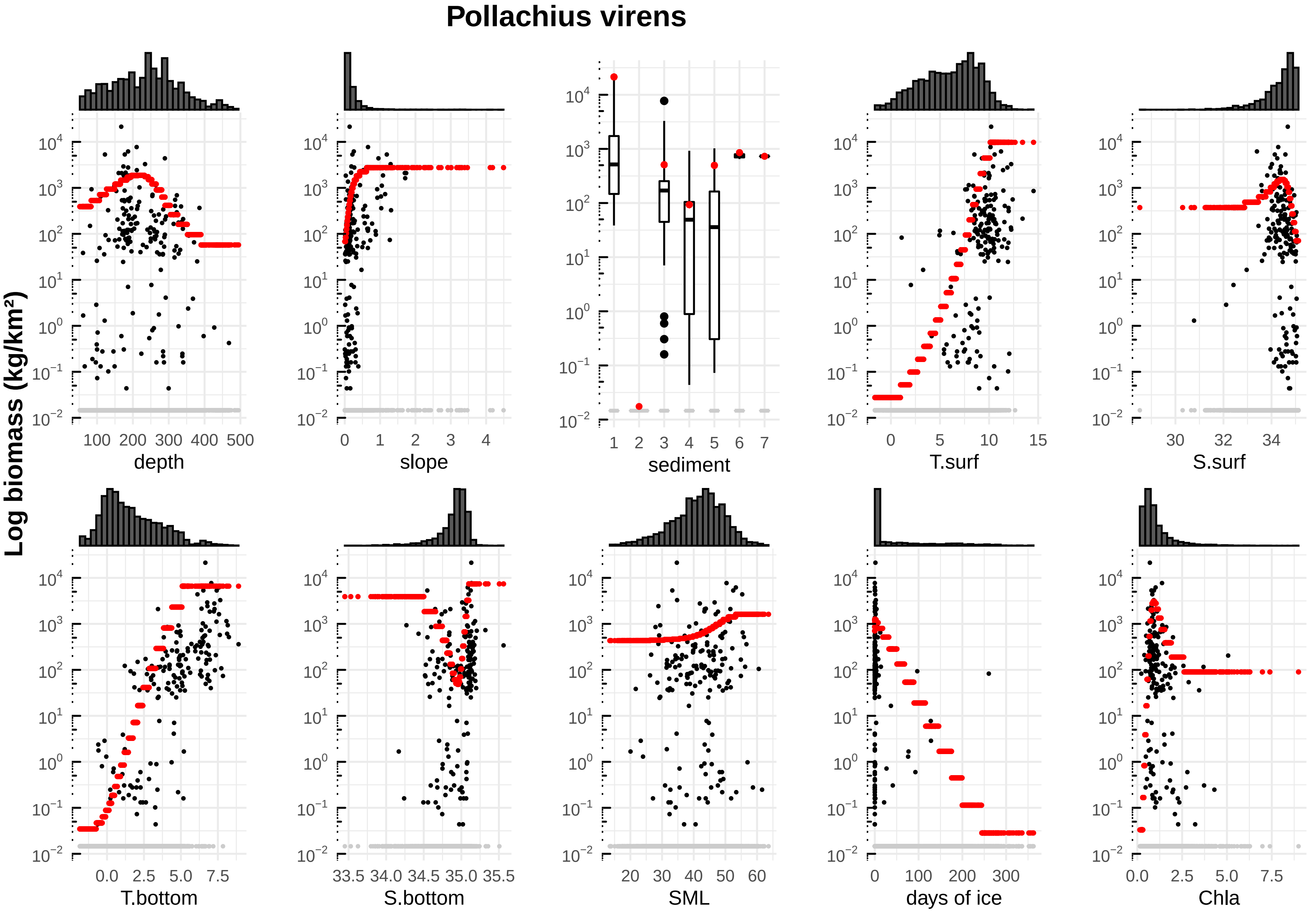

Pollachius virens
Log biomass (kg/km²)
depth
slope
sediment
T.surf
S.surf
T.bottom
S.bottom
SML
days of ice
Chla

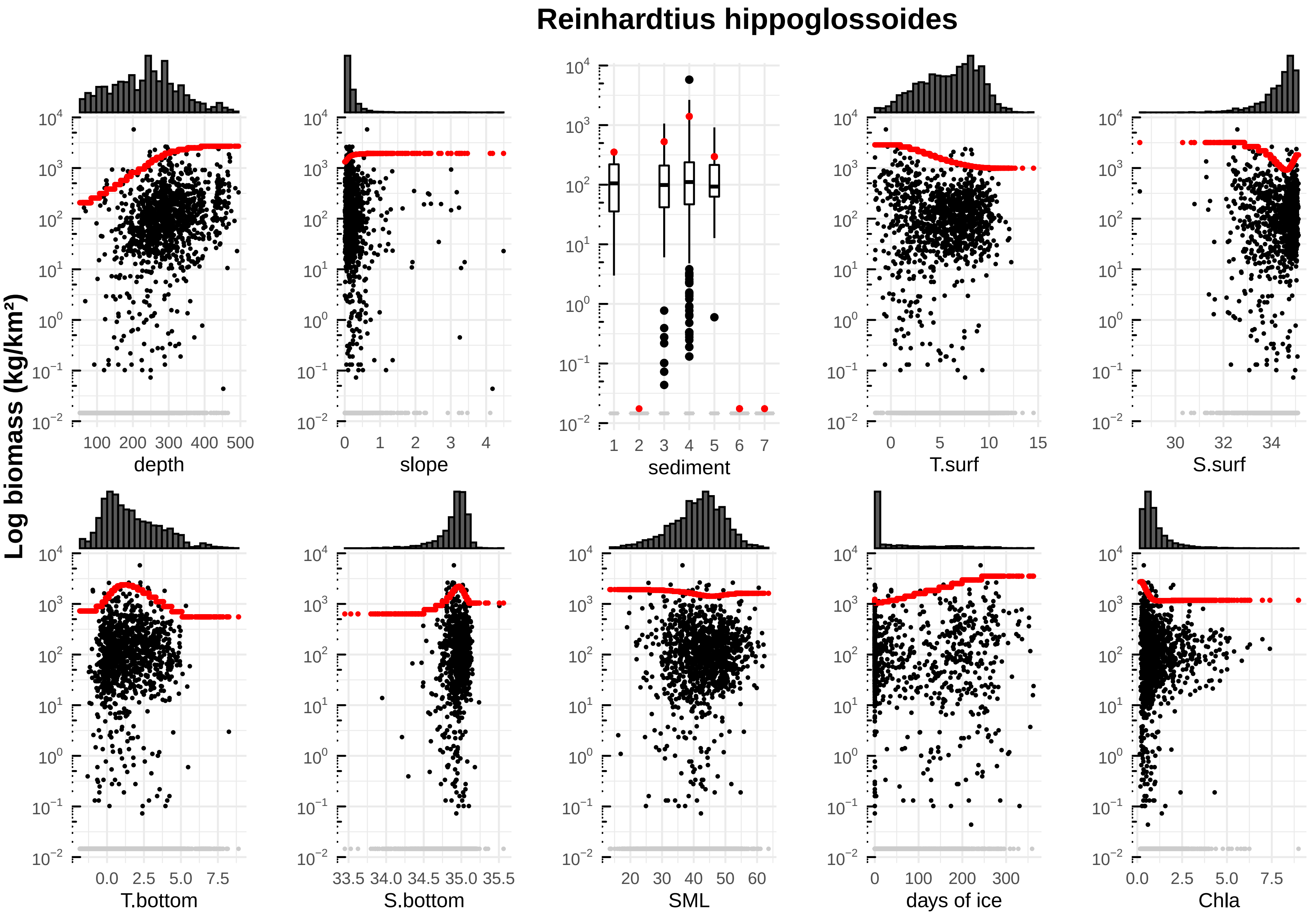

Reinhardtius hippoglossoides
Log biomass (kg/km²)
depth
slope
sediment
T.surf
S.surf
T.bottom
S.bottom
SML
days of ice
Chla
100 200 300 400 500
0 1 2 3 4
1 2 3 4 5 6 7
0 5 10 15
30 32 34
0.0 2.5 5.0 7.5
33.5 34.0 34.5 35.0 35.5
20 30 40 50 60
0 100 200 300
0.0 2.5 5.0 7.5

### Sebastes mentella

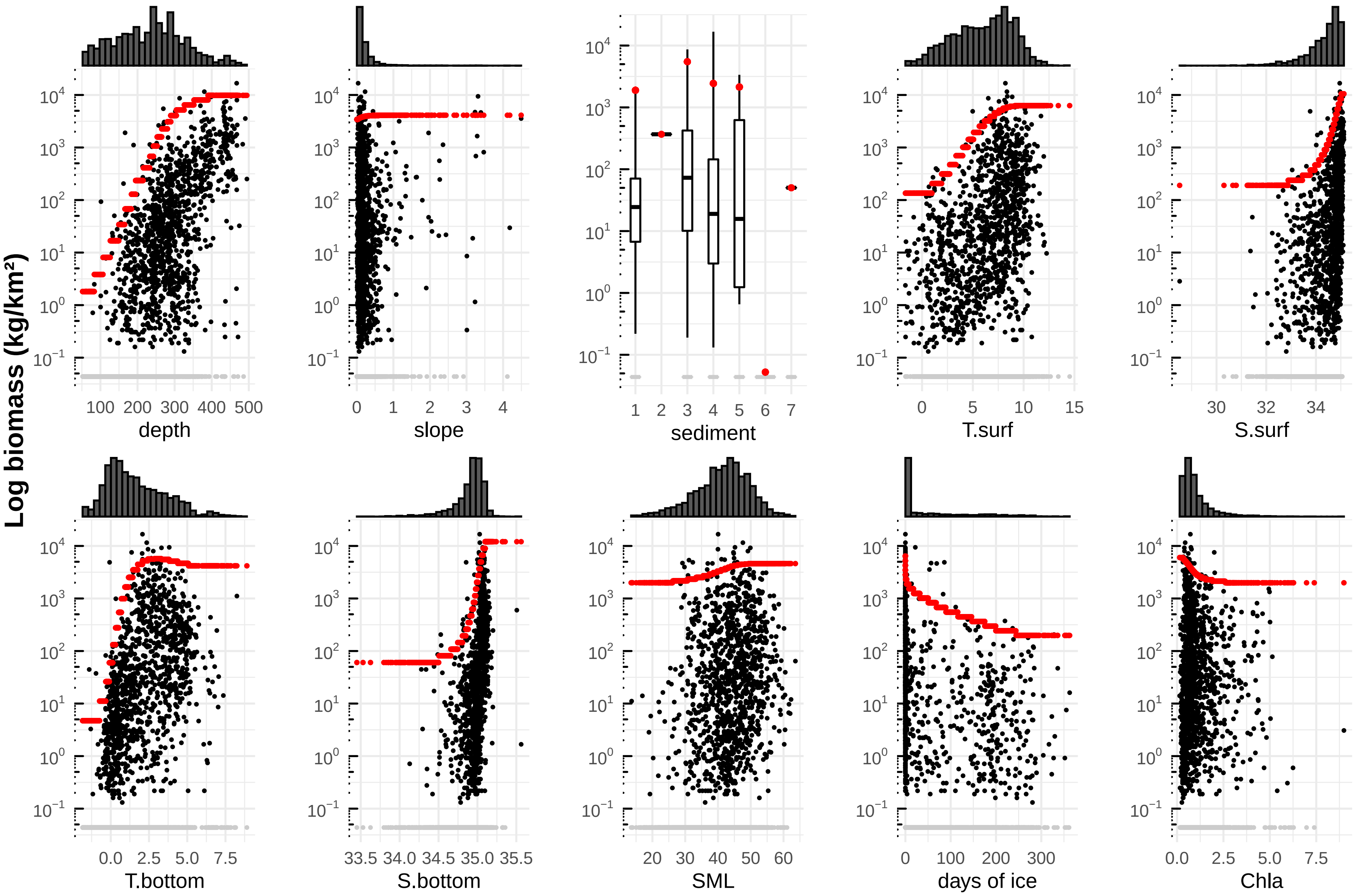

### Sebastes norvegicus

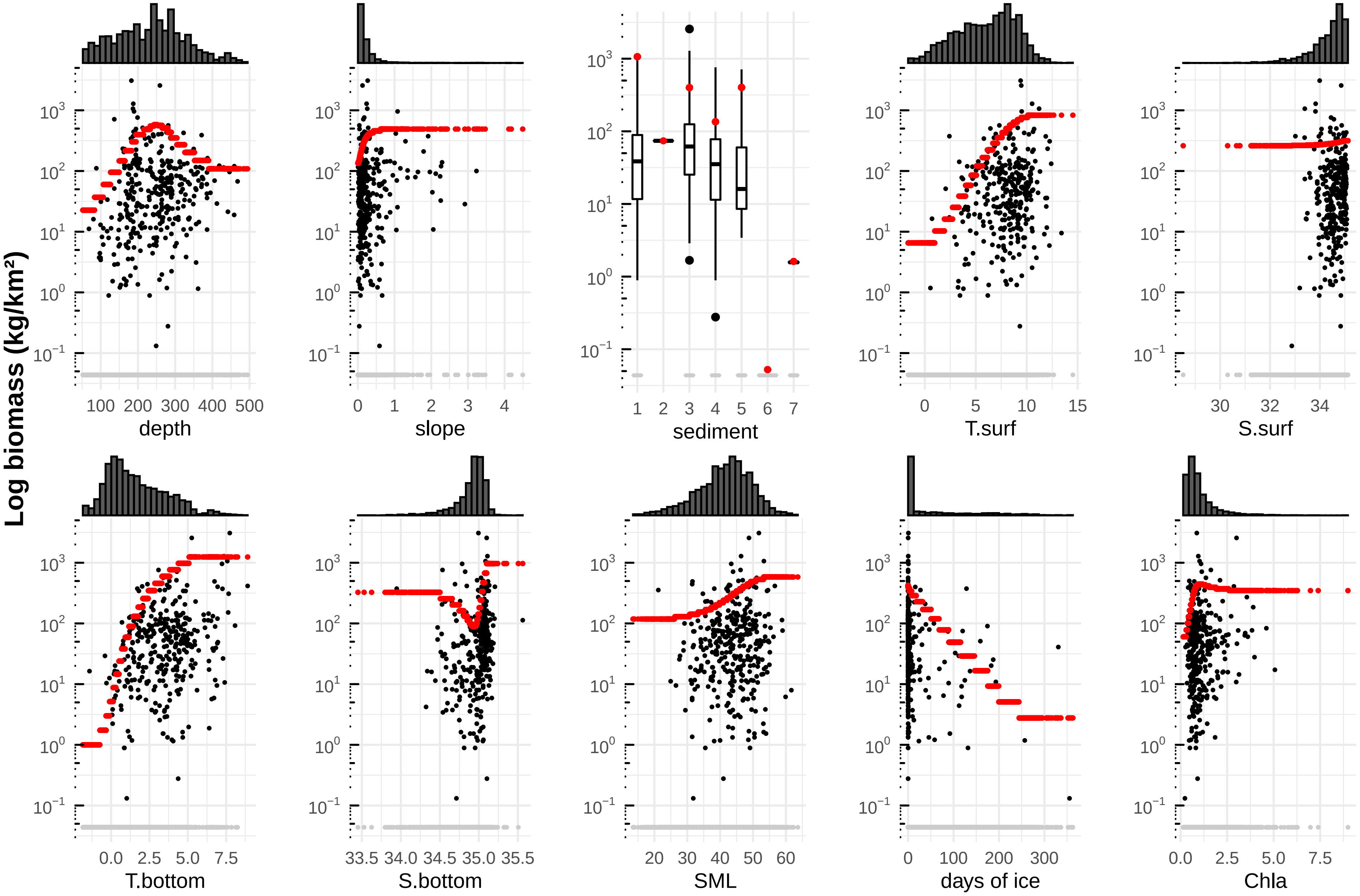

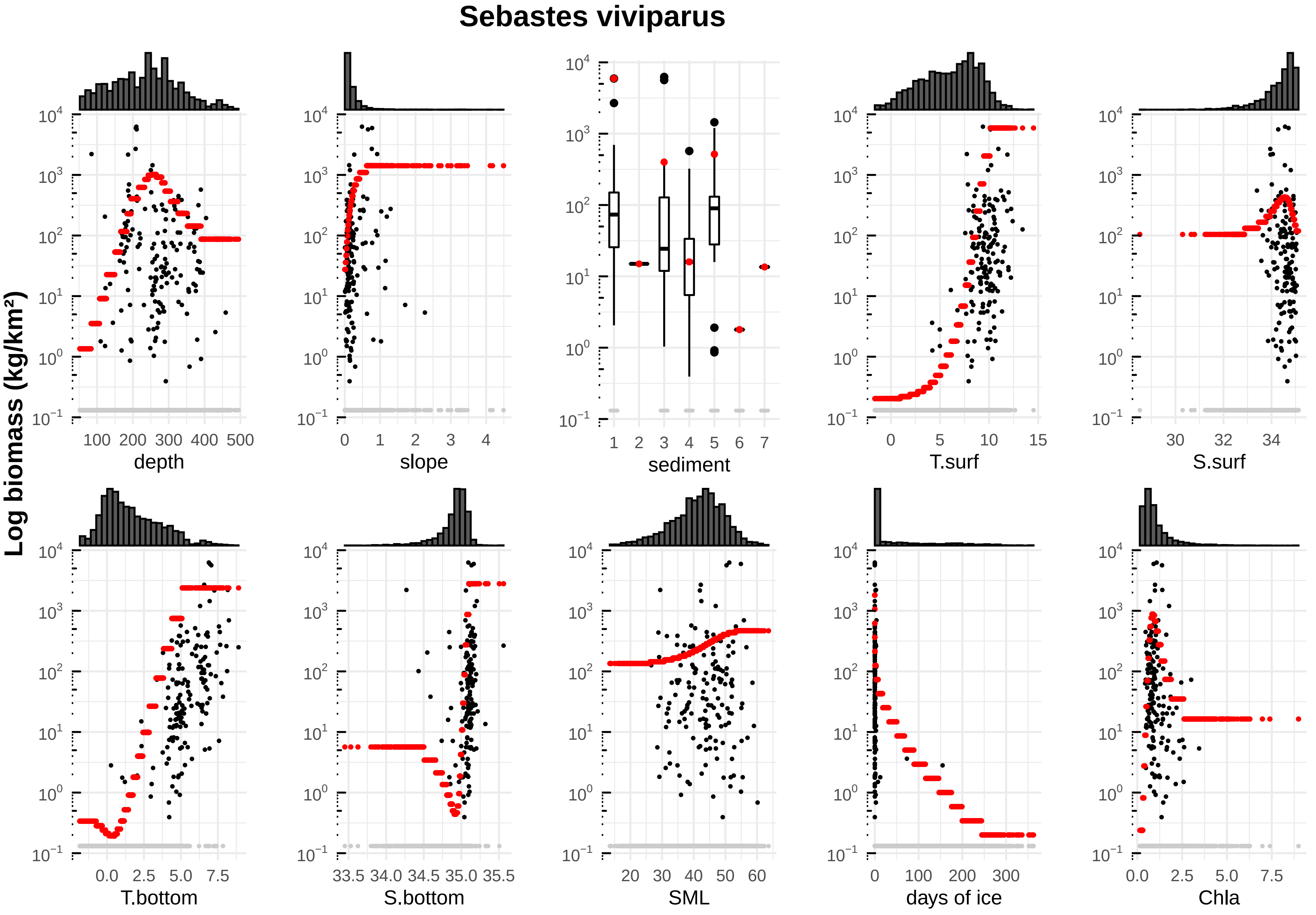
Sebastes viviparus
Log biomass (kg/km²)
depth
slope
sediment
T.surf
S.surf
T.bottom
S.bottom
SML
days of ice
Chla

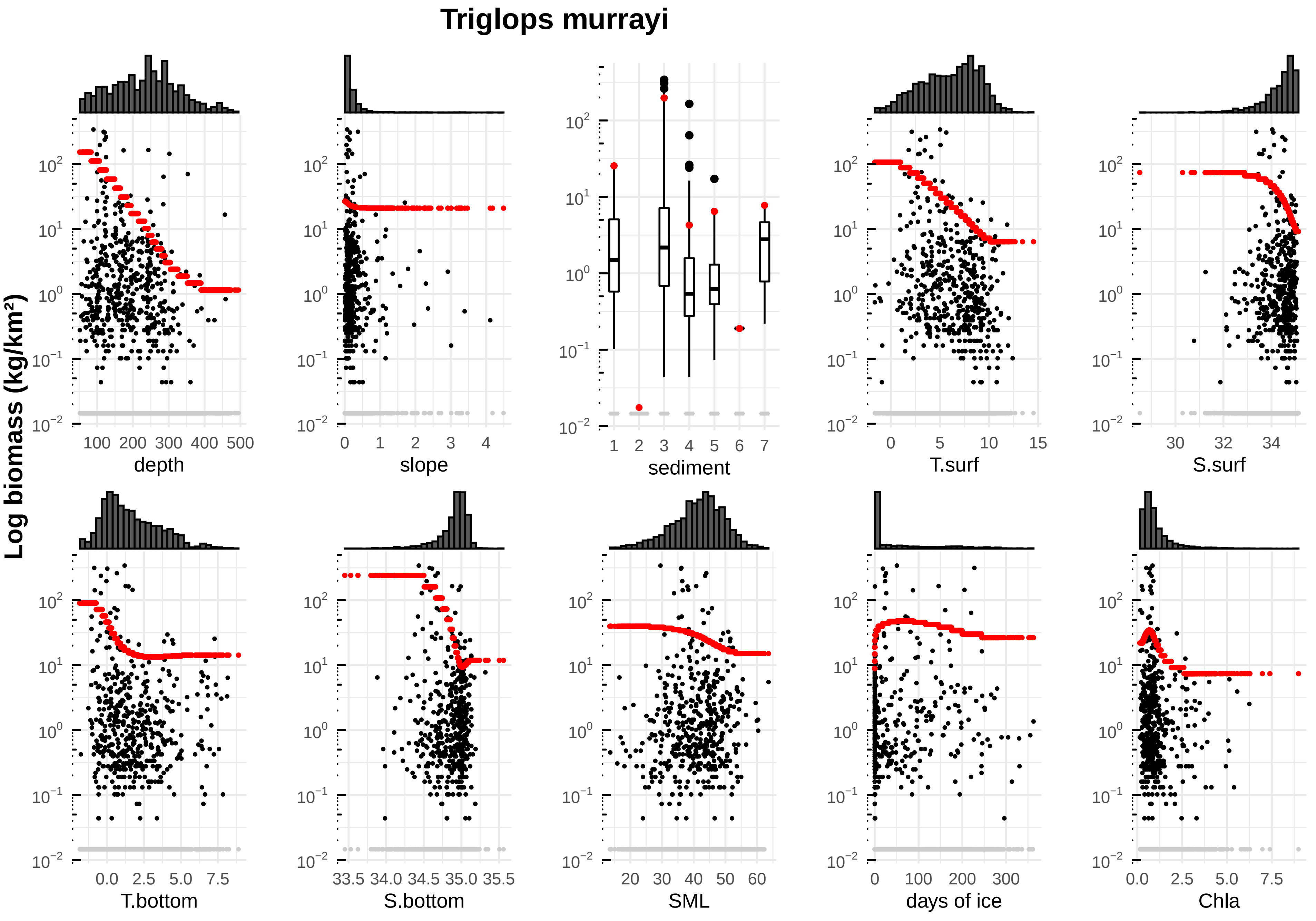
Triglops murrayi
Log biomass (kg/km²)
depth
slope
sediment
T.surf
S.surf
T.bottom
S.bottom
SML
days of ice
Chla

### Triglops nybelini

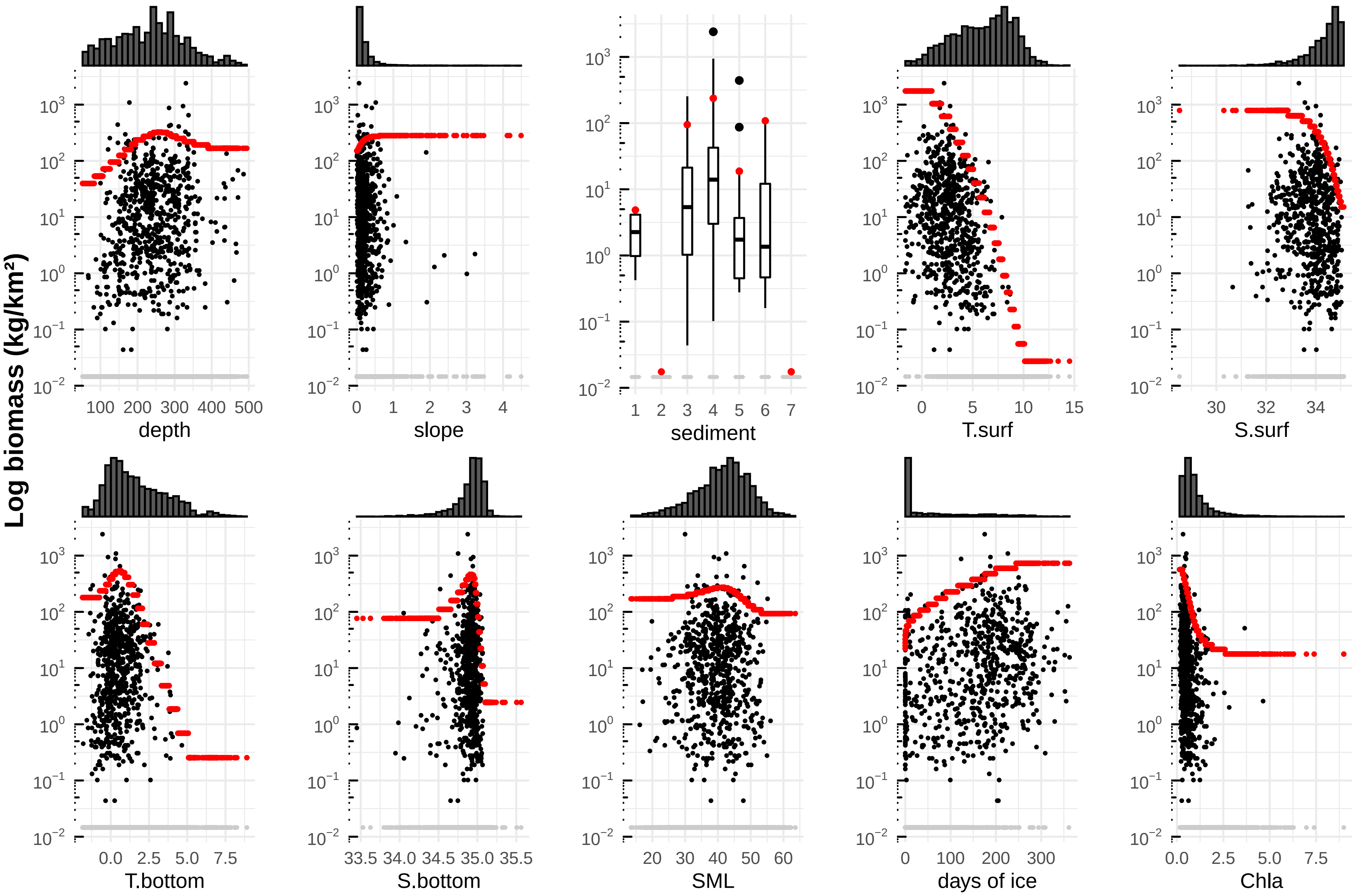

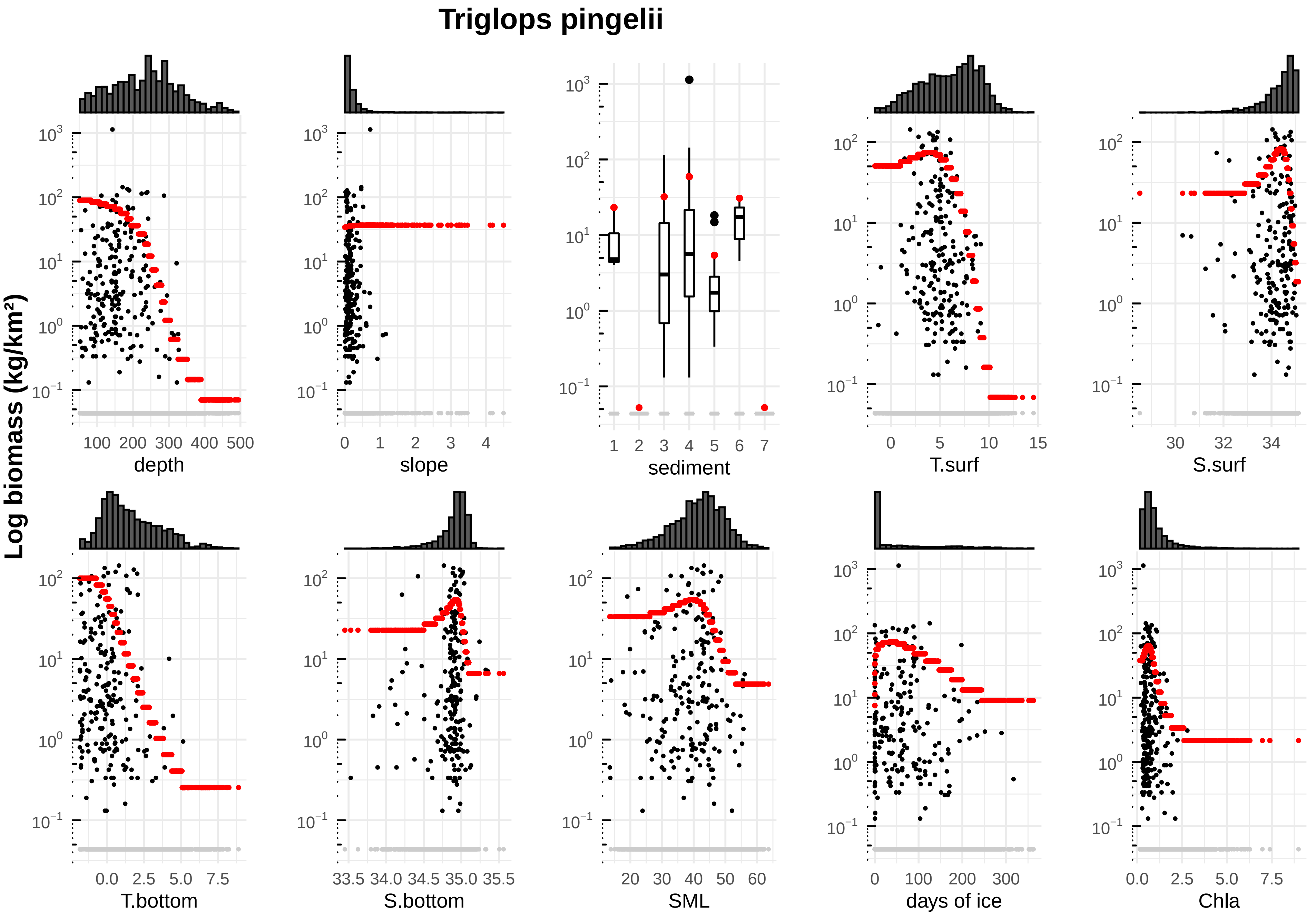

Triglops pingelii
Log biomass (kg/km²)
depth
slope
sediment
T.surf
S.surf
T.bottom
S.bottom
SML
days of ice
Chla

### Trisopterus esmarkii

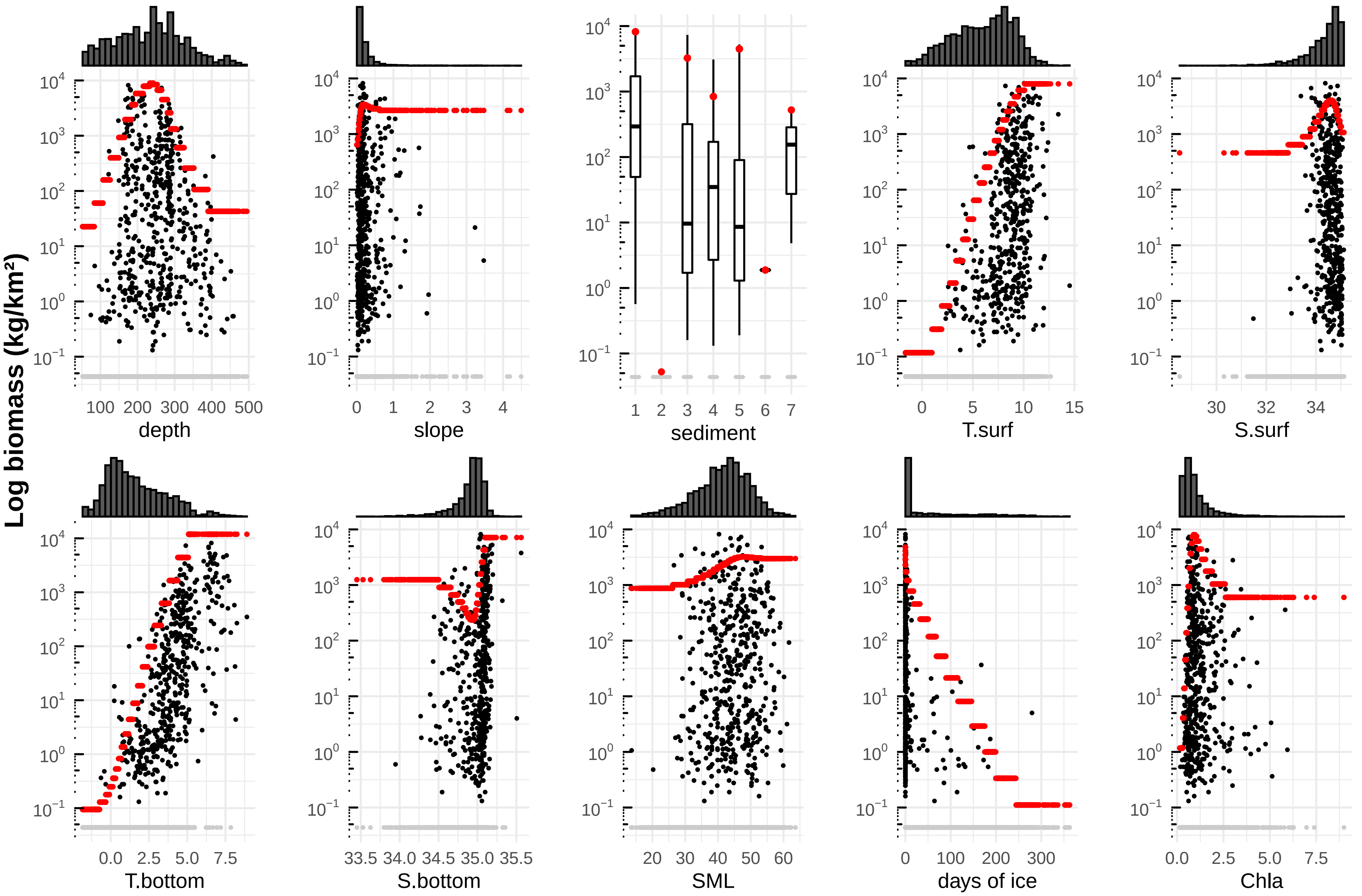

### Zoarcidae

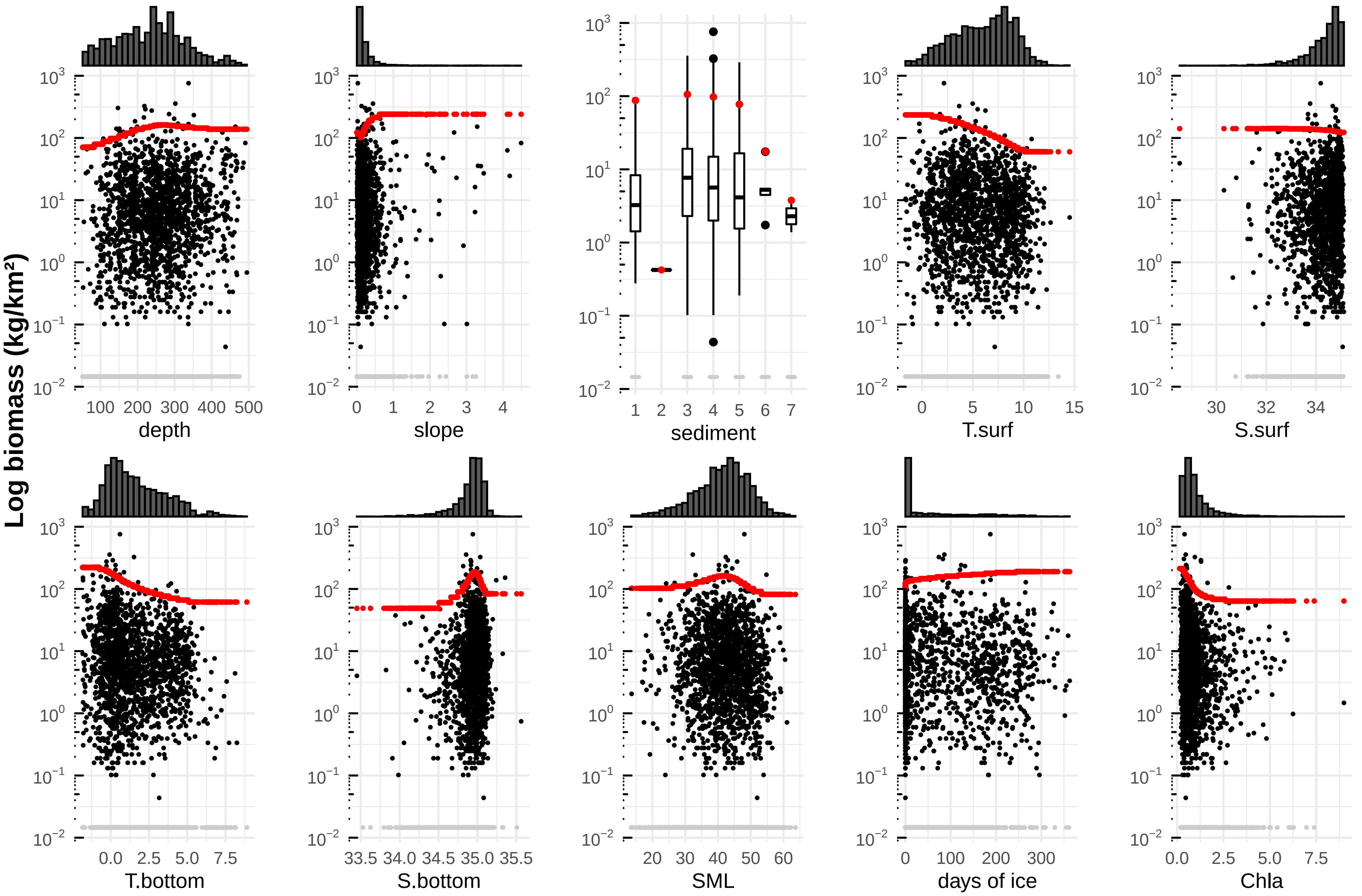

Log biomass (kg/km²)
depth
slope
sediment
T.surf
S.surf
T.bottom
S.bottom
SML
days of ice
Chla
