## Appendix 6 niche descriptions for "Suitable habitats of fish species in the Barents Sea"

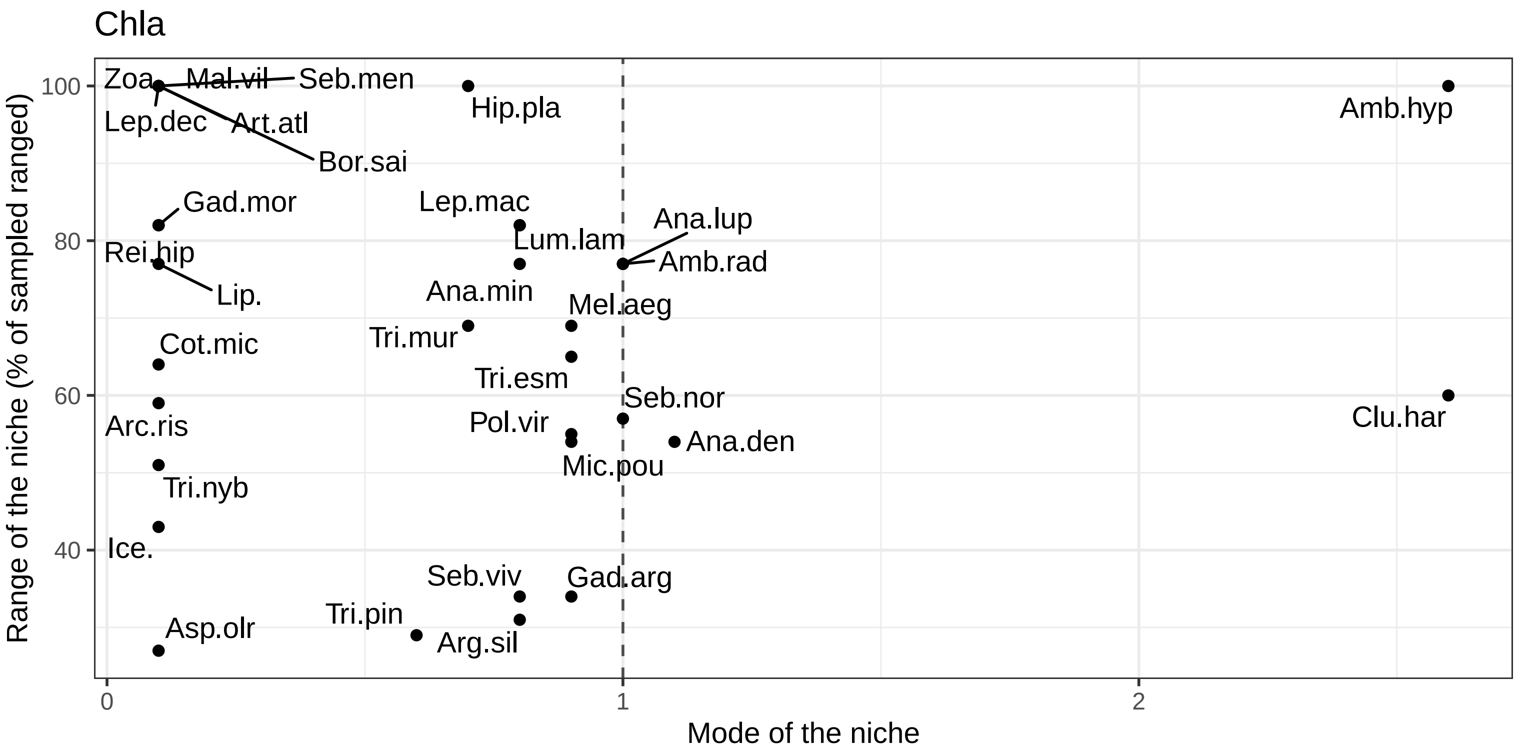


| Species | Code | Min | Max | Range | BS mean | Sp mean | Contrast | Data under the model |
|---|---|---|---|---|---|---|---|---|
| Hippoglossoides platessoides | Hip.pla | 0.1 | 9.0 | 100 | 1 | 0.7 | 0.1138 | 99.42 |
| Leptoclinus maculatus | Lep.mac | 0.1 | 7.4 | 82 | 1 | 0.8 | 0.1607 | 98.54 |
| Amblyraja hyperborea | Amb.hyp | 0.2 | 9.0 | 100 | 1 | 2.6 | 0.2353 | 99.51 |
| Leptagonus decagonus | Lep.dec | 0.1 | 9.0 | 100 | 1 | 0.1 | 0.3631 | 99.32 |
| Lumpenus lampretaeformis | Lum.lam | 0.1 | 7.4 | 82 | 1 | 0.8 | 0.5008 | 99.61 |
| Anarhichas denticulatus | Ana.den | 0.2 | 5.0 | 54 | 1 | 1.1 | 0.5385 | 98.64 |
| Reinhardtius hippoglossoides | Rei.hip | 0.1 | 7.4 | 82 | 1 | 0.1 | 0.5773 | 99.81 |
| Micromesistius poutassou | Mic.pou | 0.2 | 5.0 | 54 | 1 | 0.9 | 0.6538 | 99.61 |
| Sebastes mentella | Seb.men | 0.1 | 9.0 | 100 | 1 | 0.1 | 0.6700 | 98.25 |
| Gadus morhua | Gad.mor | 0.1 | 7.4 | 82 | 1 | 0.1 | 0.6973 | 99.81 |
| Zoarcidae | Zoa. | 0.1 | 9.0 | 100 | 1 | 0.1 | 0.6996 | 99.71 |
| Amblyraja radiata | Amb.rad | 0.1 | 7.0 | 77 | 1 | 1.0 | 0.7034 | 99.22 |
| Artediellus atlanticus | Art.atl | 0.1 | 9.0 | 100 | 1 | 0.1 | 0.7286 | 99.51 |
| Anarhichas minor | Ana.min | 0.1 | 7.0 | 77 | 1 | 0.8 | 0.7303 | 98.83 |
| Arctozenus risso | Arc.ris | 0.2 | 5.4 | 59 | 1 | 0.1 | 0.7435 | 99.22 |
| Mallotus villosus | Mal.vil | 0.1 | 9.0 | 100 | 1 | 0.1 | 0.7705 | 99.42 |
| Triglops murrayi | Tri.mur | 0.1 | 6.2 | 69 | 1 | 0.7 | 0.7870 | 98.74 |
| Boreogadus saida | Bor.sai | 0.1 | 9.0 | 100 | 1 | 0.1 | 0.7924 | 99.61 |
| Sebastes norvegicus | Seb.nor | 0.2 | 5.3 | 57 | 1 | 1.0 | 0.8650 | 99.03 |
| Anarhichas lupus | Ana.lup | 0.1 | 7.0 | 77 | 1 | 1.0 | 0.8772 | 98.93 |
| Cottunculus microps | Cot.mic | 0.2 | 5.8 | 64 | 1 | 0.1 | 0.8946 | 99.51 |
| Icelus | Ice. | 0.2 | 4.0 | 43 | 1 | 0.1 | 0.9297 | 99.90 |
| Aspidophoroides olrikii | Asp.olr | 0.2 | 2.6 | 27 | 1 | 0.1 | 0.9328 | 99.32 |
| Triglops pingelii | Tri.pin | 0.2 | 2.8 | 29 | 1 | 0.6 | 0.9674 | 99.12 |
| Triglops nybelini | Tri.nyb | 0.1 | 4.6 | 51 | 1 | 0.1 | 0.9686 | 99.03 |
| Liparidae | Lip. | 0.2 | 7.0 | 77 | 1 | 0.1 | 0.9828 | 99.81 |
| Argentina silus | Arg.sil | 0.4 | 3.1 | 31 | 1 | 0.8 | 0.9839 | 98.64 |
| Clupea harengus | Clu.har | 0.2 | 5.6 | 60 | 1 | 2.6 | 0.9859 | 99.81 |
| Melanogrammus aeglefinus | Mel.aeg | 0.2 | 6.2 | 69 | 1 | 0.9 | 0.9906 | 99.81 |
| Gadiculus argenteus | Gad.arg | 0.4 | 3.4 | 34 | 1 | 0.9 | 0.9992 | 99.12 |
| Sebastes viviparus | Seb.viv | 0.4 | 3.4 | 34 | 1 | 0.8 | 0.9999 | 99.22 |
| Trisopterus esmarkii | Tri.esm | 0.2 | 5.9 | 65 | 1 | 0.9 | 0.9999 | 99.81 |
| Pollachius virens | Pol.vir | 0.2 | 5.1 | 55 | 1 | 0.9 | 1.0000 | 98.74 |

days of ice

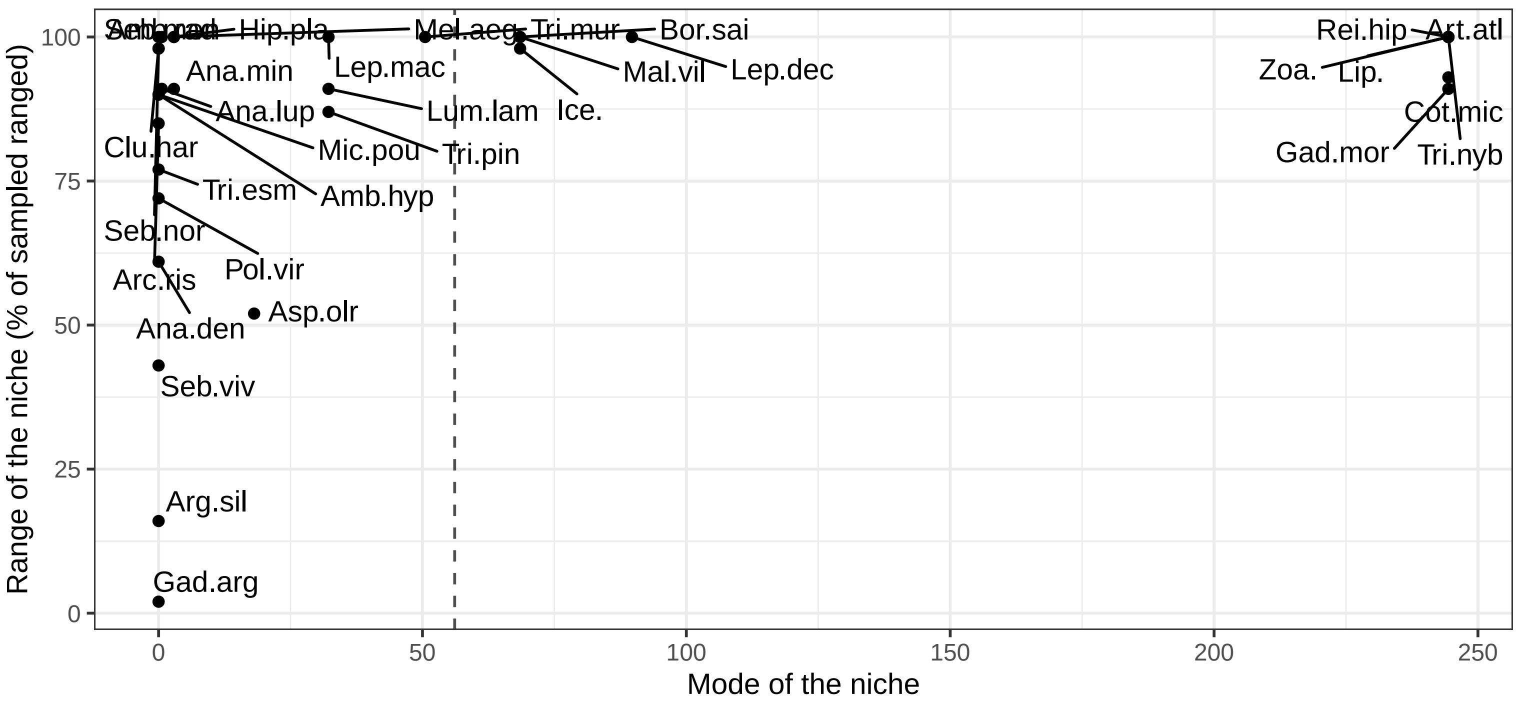


| Species | Code | Min | Max | Range | BS mean | Sp mean | Contrast | Data under the model |
|---|---|---|---|---|---|---|---|---|
| Leptoclinus maculatus | Lep.mac | 0 | 362.6 | 100 | 56.1 | 32.2 | 0.3548 | 98.64 |
| Amblyraja hyperborea | Amb.hyp | 0 | 327.2 | 90 | 56.1 | 0.0 | 0.4352 | 99.51 |
| Zoarcidae | Zoa. | 0 | 362.6 | 100 | 56.1 | 244.4 | 0.4386 | 99.90 |
| Leptagonus decagonus | Lep.dec | 0 | 362.6 | 100 | 56.1 | 89.7 | 0.5165 | 99.51 |
| Lumpenus lampretaeformis | Lum.lam | 0 | 330.6 | 91 | 56.1 | 32.2 | 0.6375 | 99.61 |
| Hippoglossoides platessoides | Hip.pla | 0 | 362.6 | 100 | 56.1 | 2.9 | 0.6692 | 99.42 |
| Reinhardtius hippoglossoides | Rei.hip | 0 | 362.6 | 100 | 56.1 | 244.4 | 0.7071 | 99.71 |
| Mallotus villosus | Mal.vil | 0 | 362.6 | 100 | 56.1 | 68.5 | 0.7956 | 99.22 |
| Artediellus atlanticus | Art.atl | 0 | 362.6 | 100 | 56.1 | 244.4 | 0.8129 | 99.51 |
| Triglops murrayi | Tri.mur | 0 | 362.6 | 100 | 56.1 | 50.5 | 0.8154 | 98.35 |
| Arctozenus risso | Arc.ris | 0 | 306.6 | 85 | 56.1 | 0.0 | 0.8177 | 99.32 |
| Gadus morhua | Gad.mor | 0 | 330.6 | 91 | 56.1 | 244.4 | 0.8345 | 99.61 |
| Anarhichas minor | Ana.min | 0 | 330.6 | 91 | 56.1 | 2.9 | 0.8349 | 98.93 |
| Amblyraja radiata | Amb.rad | 0 | 362.6 | 100 | 56.1 | 0.6 | 0.8362 | 99.32 |
| Triglops pingelii | Tri.pin | 0 | 316.9 | 87 | 56.1 | 32.2 | 0.8978 | 99.32 |
| Icelus | Ice. | 0 | 355.3 | 98 | 56.1 | 68.5 | 0.9136 | 99.81 |
| Anarhichas denticulatus | Ana.den | 0 | 220.1 | 61 | 56.1 | 0.0 | 0.9140 | 98.35 |
| Boreogadus saida | Bor.sai | 0 | 362.6 | 100 | 56.1 | 68.5 | 0.9240 | 99.81 |
| Cottunculus microps | Cot.mic | 0 | 336.3 | 93 | 56.1 | 244.4 | 0.9445 | 99.61 |
| Anarhichas lupus | Ana.lup | 0 | 330.6 | 91 | 56.1 | 0.6 | 0.9522 | 99.03 |
| Sebastes mentella | Seb.men | 0 | 362.6 | 100 | 56.1 | 0.0 | 0.9691 | 99.42 |
| Triglops nybelini | Tri.nyb | 0 | 362.6 | 100 | 56.1 | 244.4 | 0.9695 | 99.81 |
| Liparidae | Lip. | 0 | 362.6 | 100 | 56.1 | 244.4 | 0.9800 | 100.00 |
| Clupea harengus | Clu.har | 0 | 355.2 | 98 | 56.1 | 0.0 | 0.9844 | 99.51 |
| Melanogrammus aeglefinus | Mel.aeg | 0 | 362.6 | 100 | 56.1 | 2.9 | 0.9857 | 99.81 |
| Sebastes norvegicus | Seb.nor | 0 | 355.2 | 98 | 56.1 | 0.0 | 0.9935 | 98.83 |
| Aspidophoroides olrikii | Asp.olr | 0 | 188.5 | 52 | 56.1 | 18.1 | 0.9960 | 99.12 |
| Micromesistius poutassou | Mic.pou | 0 | 328.1 | 90 | 56.1 | 0.0 | 0.9994 | 99.90 |
| Gadiculus argenteus | Gad.arg | 0 | 7.9 | 2 | 56.1 | 0.0 | 0.9998 | 99.22 |
| Argentina silus | Arg.sil | 0 | 56.8 | 16 | 56.1 | 0.0 | 1.0000 | 99.12 |
| Pollachius virens | Pol.vir | 0 | 260.4 | 72 | 56.1 | 0.0 | 1.0000 | 98.64 |
| Sebastes viviparus | Seb.viv | 0 | 154.9 | 43 | 56.1 | 0.0 | 1.0000 | 99.42 |
| Trisopterus esmarkii | Tri.esm | 0 | 279.5 | 77 | 56.1 | 0.0 | 1.0000 | 99.90 |

depth

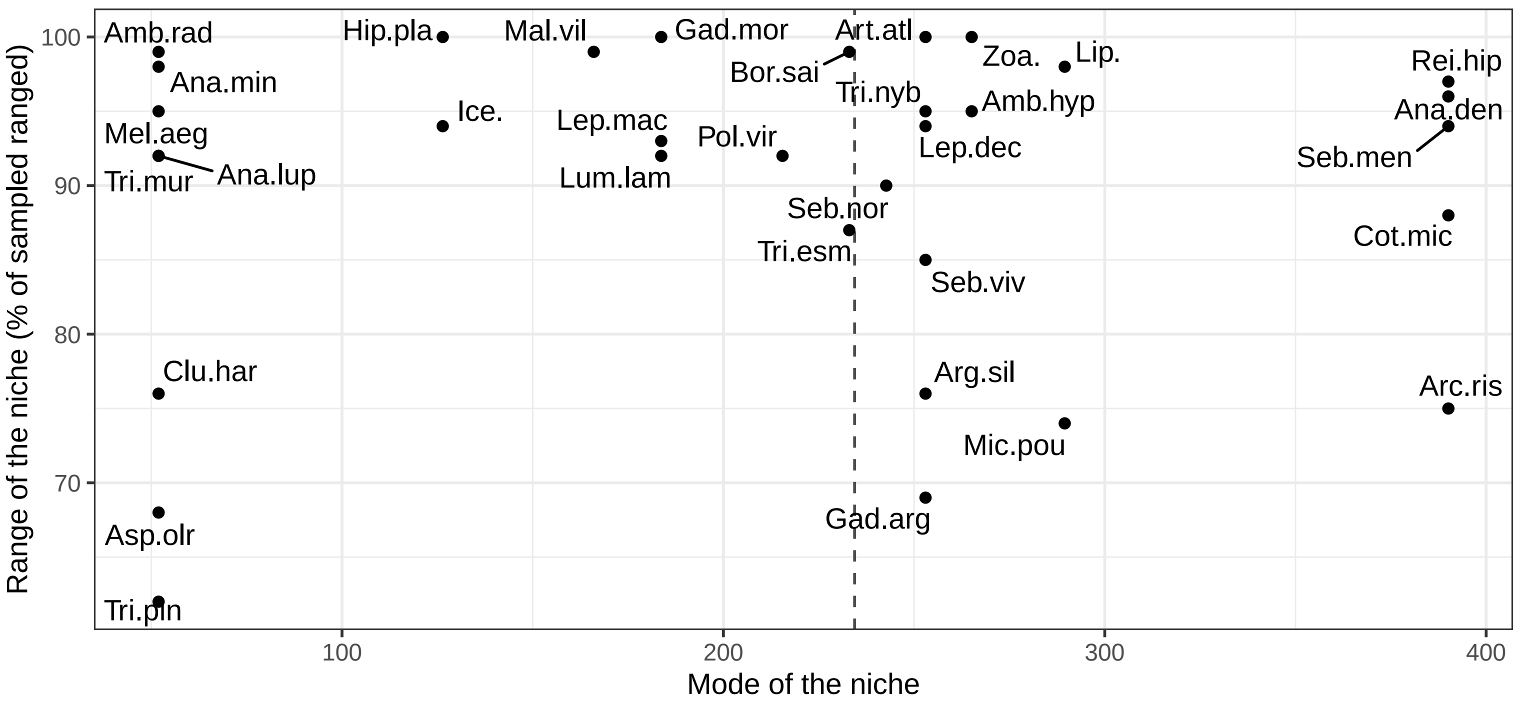


| Species | Code | Min | Max | Range | BS mean | Sp mean | Contrast | Data under the model |
|---|---|---|---|---|---|---|---|---|
| Boreogadus saida | Bor.sai | 51.9 | 490.6 | 99 | 234.4 | 233.0 | 0.4718 | 99.71 |
| Amblyraja radiata | Amb.rad | 51.9 | 490.6 | 99 | 234.4 | 51.9 | 0.4949 | 99.61 |
| Anarhichas denticulatus | Ana.den | 63.5 | 490.6 | 96 | 234.4 | 390.1 | 0.5280 | 98.15 |
| Zoarcidae | Zoa. | 53.3 | 494.8 | 100 | 234.4 | 265.1 | 0.5621 | 100.00 |
| Amblyraja hyperborea | Amb.hyp | 63.5 | 485.9 | 95 | 234.4 | 265.1 | 0.7104 | 99.61 |
| Anarhichas minor | Ana.min | 58.6 | 490.6 | 98 | 234.4 | 51.9 | 0.8258 | 98.64 |
| Leptagonus decagonus | Lep.dec | 51.9 | 468.0 | 94 | 234.4 | 253.0 | 0.8758 | 99.13 |
| Hippoglossoides platessoides | Hip.pla | 51.9 | 494.8 | 100 | 234.4 | 126.4 | 0.8781 | 99.51 |
| Triglops nybelini | Tri.nyb | 67.0 | 485.9 | 95 | 234.4 | 253.0 | 0.8782 | 99.32 |
| Mallotus villosus | Mal.vil | 52.1 | 490.6 | 99 | 234.4 | 166.0 | 0.8905 | 99.32 |
| Gadus morhua | Gad.mor | 51.9 | 494.8 | 100 | 234.4 | 183.7 | 0.9139 | 99.90 |
| Reinhardtius hippoglossoides | Rei.hip | 63.5 | 494.8 | 97 | 234.4 | 390.1 | 0.9234 | 99.81 |
| Liparidae | Lip. | 51.9 | 485.9 | 98 | 234.4 | 289.5 | 0.9417 | 99.81 |
| Sebastes norvegicus | Seb.nor | 68.3 | 467.8 | 90 | 234.4 | 242.7 | 0.9608 | 98.74 |
| Melanogrammus aeglefinus | Mel.aeg | 51.9 | 474.5 | 95 | 234.4 | 51.9 | 0.9684 | 100.00 |
| Pollachius virens | Pol.vir | 60.8 | 468.0 | 92 | 234.4 | 215.5 | 0.9693 | 98.64 |
| Artediellus atlanticus | Art.atl | 51.9 | 494.8 | 100 | 234.4 | 253.0 | 0.9711 | 99.71 |
| Clupea harengus | Clu.har | 58.6 | 397.1 | 76 | 234.4 | 51.9 | 0.9766 | 99.32 |
| Leptoclinus maculatus | Lep.mac | 51.9 | 463.4 | 93 | 234.4 | 183.7 | 0.9810 | 98.45 |
| Anarhichas lupus | Ana.lup | 57.9 | 463.9 | 92 | 234.4 | 51.9 | 0.9877 | 98.35 |
| Lumpenus lampretaeformis | Lum.lam | 53.3 | 458.6 | 92 | 234.4 | 183.7 | 0.9916 | 99.71 |
| Triglops murrayi | Tri.mur | 51.9 | 458.6 | 92 | 234.4 | 51.9 | 0.9926 | 99.22 |
| Icelus | Ice. | 53.3 | 471.0 | 94 | 234.4 | 126.4 | 0.9937 | 99.90 |
| Cottunculus microps | Cot.mic | 95.8 | 485.9 | 88 | 234.4 | 390.1 | 0.9941 | 99.51 |
| Arctozenus risso | Arc.ris | 141.0 | 474.5 | 75 | 234.4 | 390.1 | 0.9969 | 99.22 |
| Trisopterus esmarkii | Tri.esm | 74.0 | 458.6 | 87 | 234.4 | 233.0 | 0.9975 | 99.32 |
| Gadiculus argenteus | Gad.arg | 80.3 | 385.5 | 69 | 234.4 | 253.0 | 0.9988 | 98.93 |
| Sebastes viviparus | Seb.viv | 84.5 | 459.0 | 85 | 234.4 | 253.0 | 0.9988 | 99.03 |
| Argentina silus | Arg.sil | 123.5 | 459.0 | 76 | 234.4 | 253.0 | 0.9993 | 97.67 |
| Triglops pingelii | Tri.pin | 53.3 | 327.9 | 62 | 234.4 | 51.9 | 0.9997 | 99.32 |
| Sebastes mentella | Seb.men | 79.2 | 494.8 | 94 | 234.4 | 390.1 | 0.9998 | 97.76 |
| Aspidophoroides olrikii | Asp.olr | 51.9 | 353.7 | 68 | 234.4 | 51.9 | 0.9999 | 99.81 |
| Micromesistius poutassou | Mic.pou | 163.6 | 490.6 | 74 | 234.4 | 289.5 | 1.0000 | 99.42 |

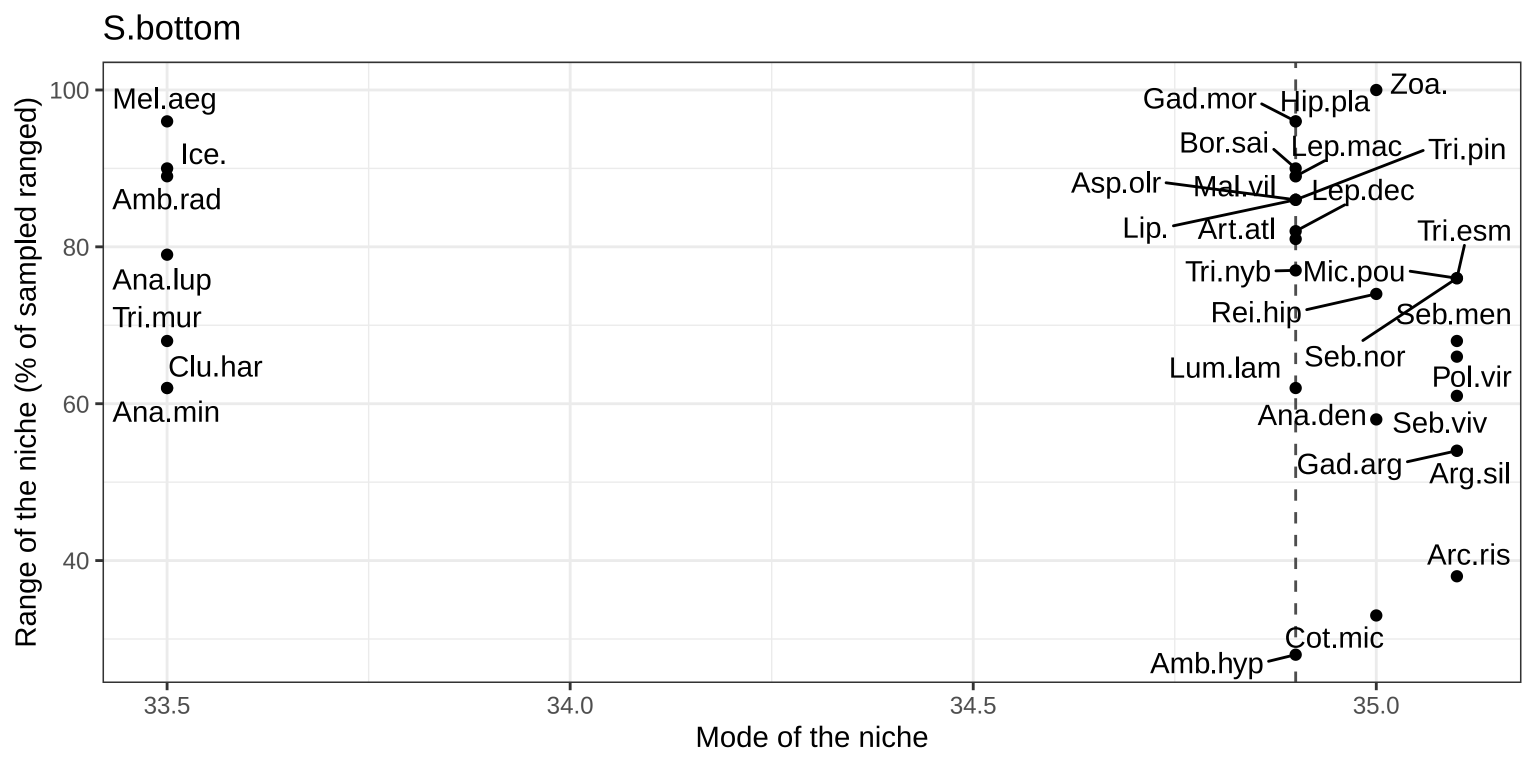


| Species | Code | Min | Max | Range | BS mean | Sp mean | Contrast | Data under the model |
|---|---|---|---|---|---|---|---|---|
| Amblyraja radiata | Amb.rad | 33.6 | 35.5 | 89 | 34.9 | 33.5 | 0.3769 | 99.41 |
| Hippoglossoides platessoides | Hip.pla | 33.5 | 35.6 | 96 | 34.9 | 34.9 | 0.4088 | 99.31 |
| Anarhichas minor | Ana.min | 33.9 | 35.2 | 62 | 34.9 | 33.5 | 0.5593 | 99.02 |
| Leptoclinus maculatus | Lep.mac | 33.5 | 35.3 | 89 | 34.9 | 34.9 | 0.5848 | 98.82 |
| Anarhichas denticulatus | Ana.den | 33.9 | 35.2 | 58 | 34.9 | 35.0 | 0.6914 | 98.24 |
| Lumpenus lampretaeformis | Lum.lam | 33.9 | 35.2 | 62 | 34.9 | 34.9 | 0.7005 | 99.51 |
| Reinhardtius hippoglossoides | Rei.hip | 33.9 | 35.5 | 74 | 34.9 | 35.0 | 0.7164 | 100.00 |
| Zoarcidae | Zoa. | 33.5 | 35.6 | 100 | 34.9 | 35.0 | 0.7394 | 99.71 |
| Aspidophoroides olrikii | Asp.olr | 33.5 | 35.4 | 86 | 34.9 | 34.9 | 0.7714 | 99.22 |
| Artediellus atlanticus | Art.atl | 33.9 | 35.6 | 81 | 34.9 | 34.9 | 0.8425 | 99.71 |
| Triglops pingelii | Tri.pin | 33.5 | 35.4 | 86 | 34.9 | 34.9 | 0.8786 | 99.22 |
| Anarhichas lupus | Ana.lup | 33.9 | 35.6 | 79 | 34.9 | 33.5 | 0.8809 | 98.24 |
| Gadus morhua | Gad.mor | 33.5 | 35.6 | 96 | 34.9 | 34.9 | 0.8923 | 99.61 |
| Boreogadus saida | Bor.sai | 33.5 | 35.4 | 90 | 34.9 | 34.9 | 0.9066 | 99.61 |
| Sebastes norvegicus | Seb.nor | 33.9 | 35.6 | 76 | 34.9 | 35.1 | 0.9084 | 98.63 |
| Leptagonus decagonus | Lep.dec | 33.6 | 35.4 | 82 | 34.9 | 34.9 | 0.9120 | 98.53 |
| Cottunculus microps | Cot.mic | 34.5 | 35.2 | 33 | 34.9 | 35.0 | 0.9218 | 99.51 |
| Melanogrammus aeglefinus | Mel.aeg | 33.5 | 35.6 | 96 | 34.9 | 33.5 | 0.9352 | 99.61 |
| Liparidae | Lip. | 33.5 | 35.4 | 86 | 34.9 | 34.9 | 0.9390 | 99.51 |
| Triglops murrayi | Tri.mur | 33.9 | 35.3 | 68 | 34.9 | 33.5 | 0.9613 | 98.92 |
| Trisopterus esmarkii | Tri.esm | 33.9 | 35.6 | 76 | 34.9 | 35.1 | 0.9672 | 99.22 |
| Icelus | Ice. | 33.5 | 35.4 | 90 | 34.9 | 33.5 | 0.9691 | 100.00 |
| Mallotus villosus | Mal.vil | 33.6 | 35.5 | 89 | 34.9 | 34.9 | 0.9728 | 99.31 |
| Gadiculus argenteus | Gad.arg | 34.4 | 35.6 | 54 | 34.9 | 35.1 | 0.9885 | 97.75 |
| Micromesistius poutassou | Mic.pou | 33.9 | 35.6 | 76 | 34.9 | 35.1 | 0.9899 | 99.12 |
| Pollachius virens | Pol.vir | 34.2 | 35.6 | 66 | 34.9 | 35.1 | 0.9934 | 98.24 |
| Clupea harengus | Clu.har | 33.9 | 35.2 | 62 | 34.9 | 33.5 | 0.9944 | 98.73 |
| Triglops nybelini | Tri.nyb | 33.5 | 35.1 | 77 | 34.9 | 34.9 | 0.9948 | 99.41 |
| Sebastes mentella | Seb.men | 34.1 | 35.6 | 68 | 34.9 | 35.1 | 0.9950 | 98.82 |
| Arctozenus risso | Arc.ris | 34.4 | 35.2 | 38 | 34.9 | 35.1 | 0.9974 | 99.22 |
| Amblyraja hyperborea | Amb.hyp | 34.6 | 35.2 | 28 | 34.9 | 34.9 | 0.9983 | 99.41 |
| Sebastes viviparus | Seb.viv | 34.3 | 35.6 | 61 | 34.9 | 35.1 | 0.9999 | 98.24 |
| Argentina silus | Arg.sil | 34.4 | 35.6 | 54 | 34.9 | 35.1 | 1.0000 | 96.67 |

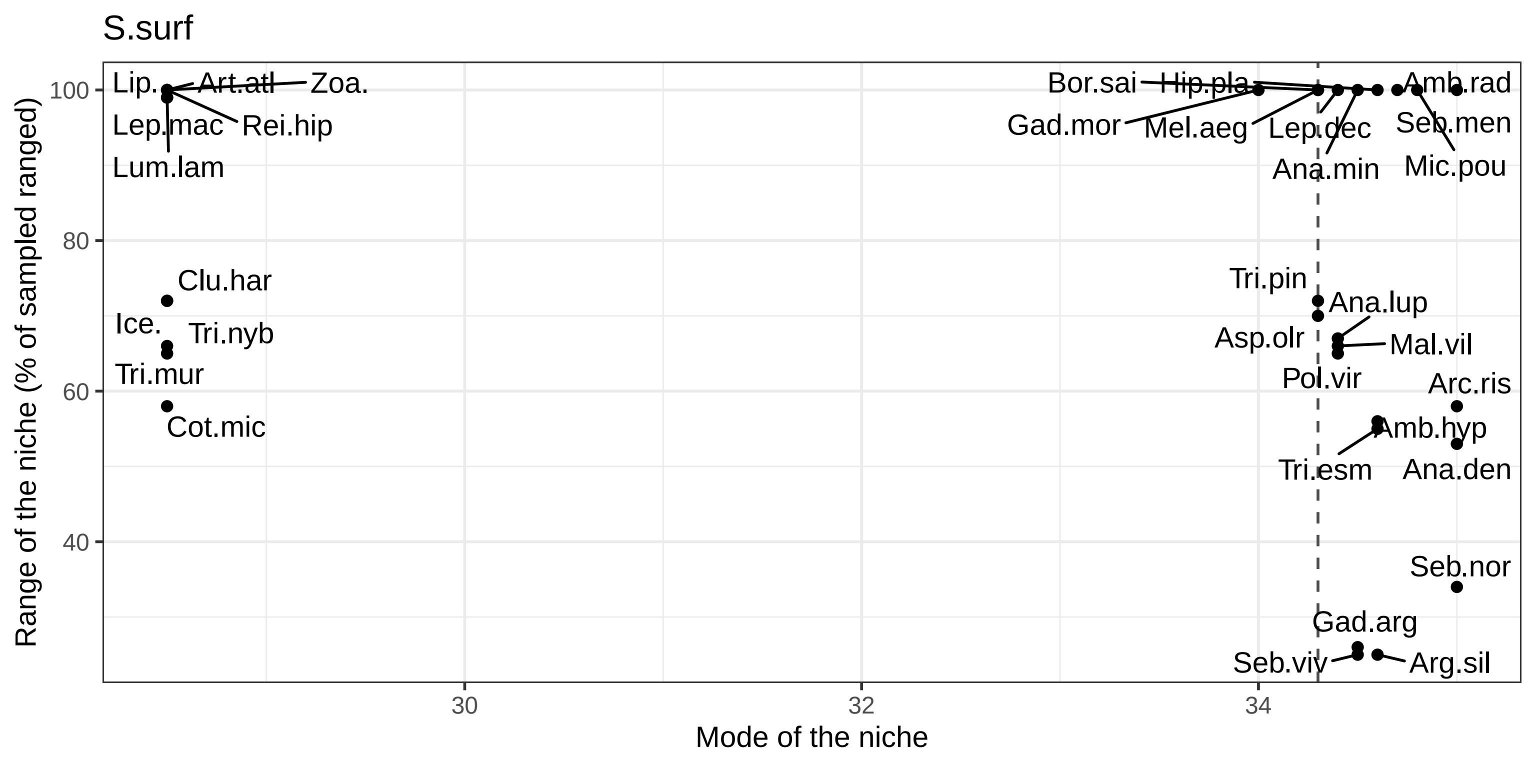


| Species | Code | Min | Max | Range | BS mean | Sp mean | Contrast | Data under the model |
|---|---|---|---|---|---|---|---|---|
| Leptagonus decagonus | Lep.dec | 28.5 | 35.1 | 100 | 34.3 | 34.4 | 0.0930 | 99.31 |
| Zoarcidae | Zoa. | 28.5 | 35.1 | 100 | 34.3 | 28.5 | 0.1295 | 99.90 |
| Sebastes norvegicus | Seb.nor | 32.9 | 35.1 | 34 | 34.3 | 35.0 | 0.1733 | 98.73 |
| Lumpenus lampretaeformis | Lum.lam | 28.5 | 35.1 | 99 | 34.3 | 28.5 | 0.3527 | 99.51 |
| Amblyraja radiata | Amb.rad | 28.5 | 35.1 | 100 | 34.3 | 34.7 | 0.4131 | 98.92 |
| Leptoclinus maculatus | Lep.mac | 28.5 | 35.1 | 99 | 34.3 | 28.5 | 0.4581 | 98.24 |
| Anarhichas minor | Ana.min | 28.5 | 35.1 | 100 | 34.3 | 34.5 | 0.4889 | 98.82 |
| Hippoglossoides platessoides | Hip.pla | 28.5 | 35.1 | 100 | 34.3 | 34.6 | 0.5371 | 99.51 |
| Amblyraja hyperborea | Amb.hyp | 31.3 | 35.0 | 56 | 34.3 | 34.6 | 0.5965 | 99.80 |
| Anarhichas denticulatus | Ana.den | 31.6 | 35.1 | 53 | 34.3 | 35.0 | 0.6650 | 98.33 |
| Reinhardtius hippoglossoides | Rei.hip | 28.5 | 35.1 | 100 | 34.3 | 28.5 | 0.7128 | 99.80 |
| Sebastes viviparus | Seb.viv | 33.4 | 35.1 | 25 | 34.3 | 34.5 | 0.7581 | 99.31 |
| Artediellus atlanticus | Art.atl | 28.5 | 35.1 | 100 | 34.3 | 28.5 | 0.7796 | 99.71 |
| Anarhichas lupus | Ana.lup | 30.7 | 35.1 | 67 | 34.3 | 34.4 | 0.8016 | 98.63 |
| Arctozenus risso | Arc.ris | 31.3 | 35.1 | 58 | 34.3 | 35.0 | 0.8518 | 99.41 |
| Gadiculus argenteus | Gad.arg | 33.3 | 35.1 | 26 | 34.3 | 34.5 | 0.8583 | 98.82 |
| Triglops murrayi | Tri.mur | 30.8 | 35.1 | 65 | 34.3 | 28.5 | 0.8766 | 98.82 |
| Gadus morhua | Gad.mor | 28.5 | 35.1 | 100 | 34.3 | 34.0 | 0.8851 | 99.41 |
| Trisopterus esmarkii | Tri.esm | 31.5 | 35.1 | 55 | 34.3 | 34.6 | 0.8869 | 99.41 |
| Mallotus villosus | Mal.vil | 30.8 | 35.1 | 66 | 34.3 | 34.4 | 0.8925 | 99.31 |
| Cottunculus microps | Cot.mic | 31.3 | 35.1 | 58 | 34.3 | 28.5 | 0.8990 | 99.71 |
| Melanogrammus aeglefinus | Mel.aeg | 28.5 | 35.1 | 100 | 34.3 | 34.3 | 0.9046 | 99.71 |
| Argentina silus | Arg.sil | 33.4 | 35.1 | 25 | 34.3 | 34.6 | 0.9069 | 98.14 |
| Icelus | Ice. | 30.3 | 35.0 | 72 | 34.3 | 28.5 | 0.9419 | 99.90 |
| Pollachius virens | Pol.vir | 30.8 | 35.1 | 65 | 34.3 | 34.4 | 0.9535 | 98.63 |
| Triglops pingelii | Tri.pin | 30.3 | 35.0 | 72 | 34.3 | 34.3 | 0.9782 | 99.12 |
| Clupea harengus | Clu.har | 30.3 | 35.1 | 72 | 34.3 | 28.5 | 0.9800 | 99.22 |
| Triglops nybelini | Tri.nyb | 30.7 | 35.0 | 66 | 34.3 | 28.5 | 0.9809 | 99.61 |
| Sebastes mentella | Seb.men | 28.5 | 35.1 | 100 | 34.3 | 35.0 | 0.9819 | 98.43 |
| Liparidae | Lip. | 28.5 | 35.1 | 100 | 34.3 | 28.5 | 0.9845 | 100.00 |
| Micromesistius poutassou | Mic.pou | 28.5 | 35.1 | 100 | 34.3 | 34.8 | 0.9888 | 99.51 |
| Boreogadus saida | Bor.sai | 28.5 | 35.1 | 100 | 34.3 | 34.3 | 0.9928 | 99.61 |
| Aspidophoroides olrikii | Asp.olr | 30.3 | 34.9 | 70 | 34.3 | 34.3 | 0.9990 | 99.22 |

sediment

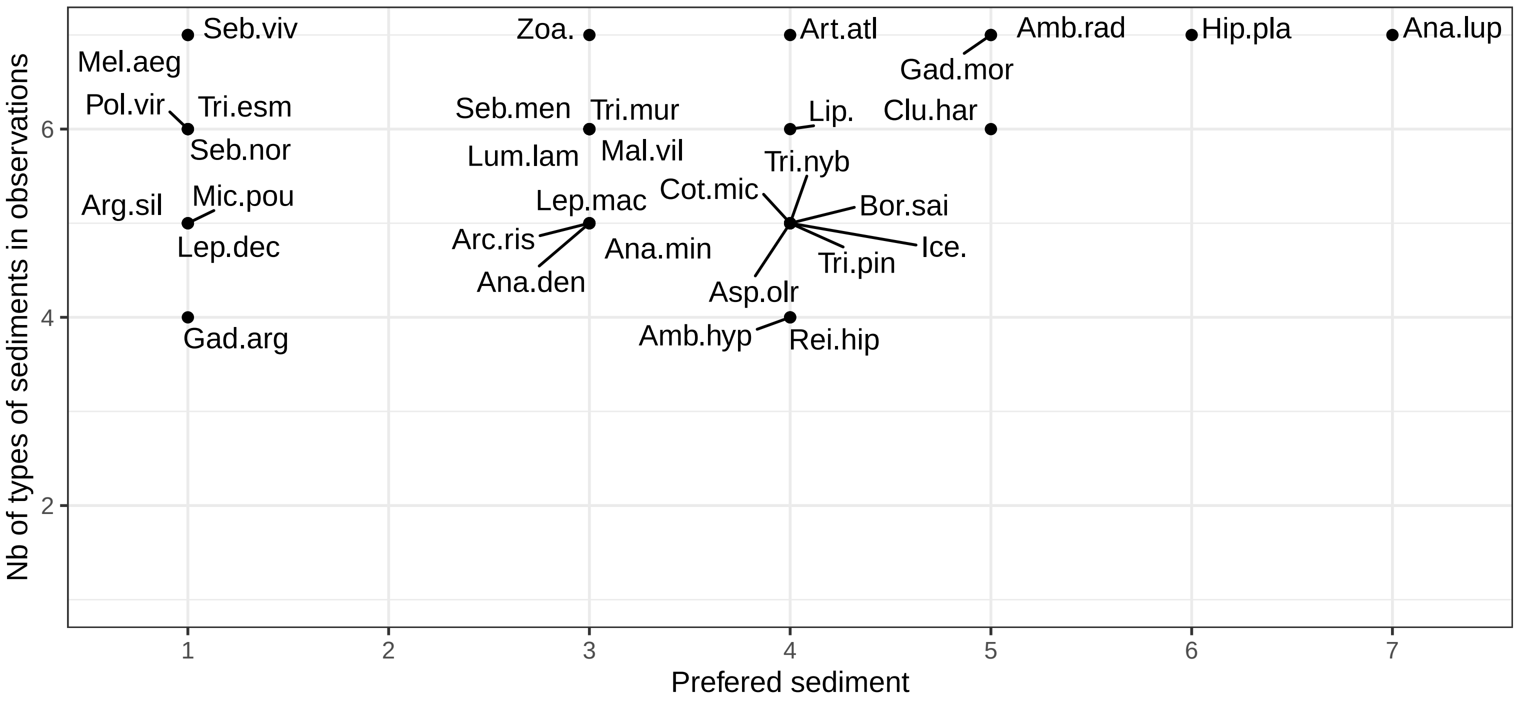


| Species | Code | Prefered sediment | Nb of sediment with obs | Contrast | Data under the model |
|---|---|---|---|---|---|
| Zoarcidae | Zoa. | Mixed sediment | 7 | 0.2683 | 99.49 |
| Amblyraja hyperborea | Amb.hyp | Mud, clay and sandy mud | 4 | 0.3990 | 98.36 |
| Amblyraja radiata | Amb.rad | Sand and muddy sand | 7 | 0.5275 | 98.57 |
| Anarhichas denticulatus | Ana.den | Mixed sediment | 5 | 0.5579 | 96.42 |
| Hippoglossoides platessoides | Hip.pla | Sand, gravel and pebbles | 7 | 0.6431 | 98.98 |
| Sebastes mentella | Seb.men | Mixed sediment | 6 | 0.6540 | 97.44 |
| Gadus morhua | Gad.mor | Sand and muddy sand | 7 | 0.6718 | 99.28 |
| Micromesistius poutassou | Mic.pou | Coarse sediment | 5 | 0.6801 | 98.57 |
| Artediellus atlanticus | Art.atl | Mud, clay and sandy mud | 7 | 0.7000 | 99.59 |
| Anarhichas minor | Ana.min | Mixed sediment | 5 | 0.7506 | 98.67 |
| Leptoclinus maculatus | Lep.mac | Mixed sediment | 5 | 0.7507 | 97.03 |
| Reinhardtius hippoglossoides | Rei.hip | Mud, clay and sandy mud | 4 | 0.7884 | 99.28 |
| Lumpenus lampretaeformis | Lum.lam | Mixed sediment | 6 | 0.8696 | 99.69 |
| Sebastes norvegicus | Seb.nor | Coarse sediment | 6 | 0.8727 | 97.96 |
| Anarhichas lupus | Ana.lup | Thin/discont. sedim. cover on bedrock | 7 | 0.8786 | 98.88 |
| Leptagonus decagonus | Lep.dec | Coarse sediment | 5 | 0.8973 | 99.08 |
| Trisopterus esmarkii | Tri.esm | Coarse sediment | 6 | 0.8976 | 99.90 |
| Triglops pingelii | Tri.pin | Mud, clay and sandy mud | 5 | 0.9095 | 99.18 |
| Icelus | Ice. | Mud, clay and sandy mud | 5 | 0.9248 | 99.39 |
| Boreogadus saida | Bor.sai | Mud, clay and sandy mud | 5 | 0.9329 | 99.39 |
| Cottunculus microps | Cot.mic | Mud, clay and sandy mud | 5 | 0.9471 | 98.57 |
| Aspidophoroides olrikii | Asp.olr | Mud, clay and sandy mud | 5 | 0.9571 | 99.28 |
| Mallotus villosus | Mal.vil | Mixed sediment | 6 | 0.9577 | 98.77 |
| Liparidae | Lip. | Mud, clay and sandy mud | 6 | 0.9598 | 99.39 |
| Melanogrammus aeglefinus | Mel.aeg | Coarse sediment | 7 | 0.9612 | 99.08 |
| Clupea harengus | Clu.har | Sand and muddy sand | 6 | 0.9632 | 99.49 |
| Triglops murrayi | Tri.mur | Mixed sediment | 6 | 0.9784 | 98.36 |
| Triglops nybelini | Tri.nyb | Mud, clay and sandy mud | 5 | 0.9796 | 98.88 |
| Argentina silus | Arg.sil | Coarse sediment | 5 | 0.9823 | 96.83 |
| Gadiculus argenteus | Gad.arg | Coarse sediment | 4 | 0.9849 | 97.75 |
| Arctozenus risso | Arc.ris | Mixed sediment | 5 | 0.9953 | 87.63 |
| Pollachius virens | Pol.vir | Coarse sediment | 6 | 0.9956 | 98.77 |
| Sebastes viviparus | Seb.viv | Coarse sediment | 7 | 0.9973 | 98.67 |

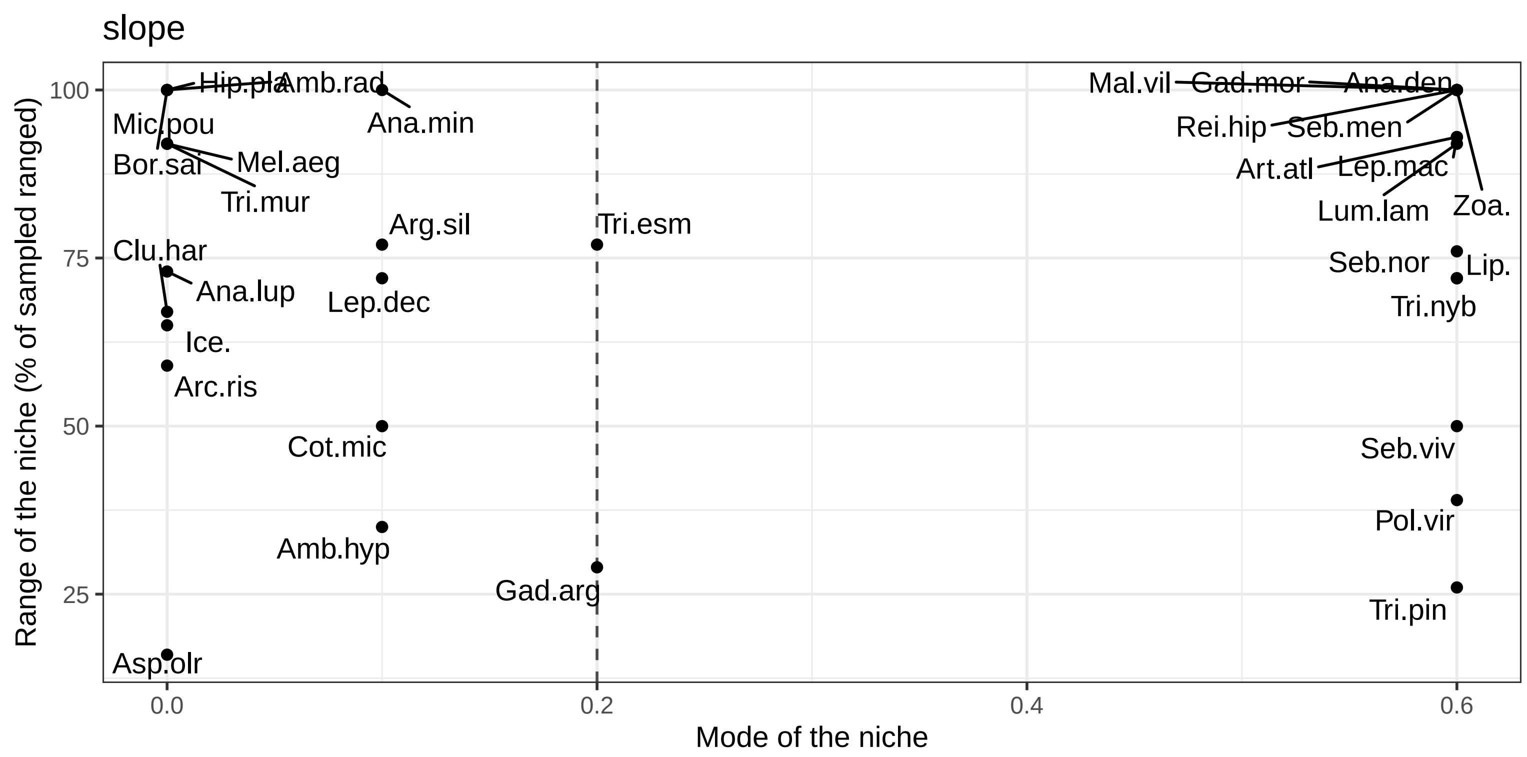


| Species | Code | Min | Max | Range | BS mean | Sp mean | Contrast | Data under the model |
|---|---|---|---|---|---|---|---|---|
| Boreogadus saida | Bor.sai | 0 | 4.5 | 100 | 0.2 | 0.0 | 0.0650 | 99.71 |
| Triglops pingelii | Tri.pin | 0 | 1.2 | 26 | 0.2 | 0.6 | 0.0674 | 99.32 |
| Leptagonus decagonus | Lep.dec | 0 | 3.2 | 72 | 0.2 | 0.1 | 0.1313 | 99.42 |
| Amblyraja radiata | Amb.rad | 0 | 4.5 | 100 | 0.2 | 0.0 | 0.1414 | 99.61 |
| Sebastes mentella | Seb.men | 0 | 4.5 | 100 | 0.2 | 0.6 | 0.1679 | 98.83 |
| Hippoglossoides platessoides | Hip.pla | 0 | 4.5 | 100 | 0.2 | 0.0 | 0.1874 | 99.32 |
| Amblyraja hyperborea | Amb.hyp | 0 | 1.6 | 35 | 0.2 | 0.1 | 0.2079 | 99.61 |
| Triglops murrayi | Tri.mur | 0 | 4.1 | 92 | 0.2 | 0.0 | 0.2154 | 98.54 |
| Arctozenus risso | Arc.ris | 0 | 2.7 | 59 | 0.2 | 0.0 | 0.2483 | 99.12 |
| Micromesistius poutassou | Mic.pou | 0 | 4.5 | 100 | 0.2 | 0.0 | 0.2655 | 99.51 |
| Anarhichas lupus | Ana.lup | 0 | 3.3 | 73 | 0.2 | 0.0 | 0.3060 | 98.74 |
| Reinhardtius hippoglossoides | Rei.hip | 0 | 4.5 | 100 | 0.2 | 0.6 | 0.3169 | 99.90 |
| Artediellus atlanticus | Art.atl | 0 | 4.2 | 93 | 0.2 | 0.6 | 0.3343 | 99.61 |
| Icelus | Ice. | 0 | 2.9 | 65 | 0.2 | 0.0 | 0.3400 | 99.71 |
| Anarhichas denticulatus | Ana.den | 0 | 4.5 | 100 | 0.2 | 0.6 | 0.4010 | 98.44 |
| Melanogrammus aeglefinus | Mel.aeg | 0 | 4.1 | 92 | 0.2 | 0.0 | 0.4275 | 99.61 |
| Mallotus villosus | Mal.vil | 0 | 4.5 | 100 | 0.2 | 0.6 | 0.4318 | 99.42 |
| Aspidophoroides olrikii | Asp.olr | 0 | 0.7 | 16 | 0.2 | 0.0 | 0.4580 | 99.03 |
| Triglops nybelini | Tri.nyb | 0 | 3.2 | 72 | 0.2 | 0.6 | 0.4644 | 99.32 |
| Cottunculus microps | Cot.mic | 0 | 2.2 | 50 | 0.2 | 0.1 | 0.4683 | 99.71 |
| Argentina silus | Arg.sil | 0 | 3.5 | 77 | 0.2 | 0.1 | 0.4752 | 97.96 |
| Leptoclinus maculatus | Lep.mac | 0 | 4.2 | 93 | 0.2 | 0.6 | 0.5378 | 98.54 |
| Clupea harengus | Clu.har | 0 | 3.0 | 67 | 0.2 | 0.0 | 0.5706 | 99.51 |
| Zoarcidae | Zoa. | 0 | 4.5 | 100 | 0.2 | 0.6 | 0.5747 | 99.81 |
| Anarhichas minor | Ana.min | 0 | 4.5 | 100 | 0.2 | 0.1 | 0.6028 | 98.93 |
| Liparidae | Lip. | 0 | 3.4 | 76 | 0.2 | 0.6 | 0.6356 | 99.81 |
| Gadus morhua | Gad.mor | 0 | 4.5 | 100 | 0.2 | 0.6 | 0.6518 | 99.81 |
| Lumpenus lampretaeformis | Lum.lam | 0 | 4.1 | 92 | 0.2 | 0.6 | 0.7155 | 99.61 |
| Sebastes norvegicus | Seb.nor | 0 | 3.2 | 72 | 0.2 | 0.6 | 0.7259 | 98.54 |
| Gadiculus argenteus | Gad.arg | 0 | 1.3 | 29 | 0.2 | 0.2 | 0.7437 | 98.83 |
| Trisopterus esmarkii | Tri.esm | 0 | 3.5 | 77 | 0.2 | 0.2 | 0.8130 | 99.42 |
| Pollachius virens | Pol.vir | 0 | 1.7 | 39 | 0.2 | 0.6 | 0.9753 | 97.96 |
| Sebastes viviparus | Seb.viv | 0 | 2.3 | 50 | 0.2 | 0.6 | 0.9807 | 98.74 |

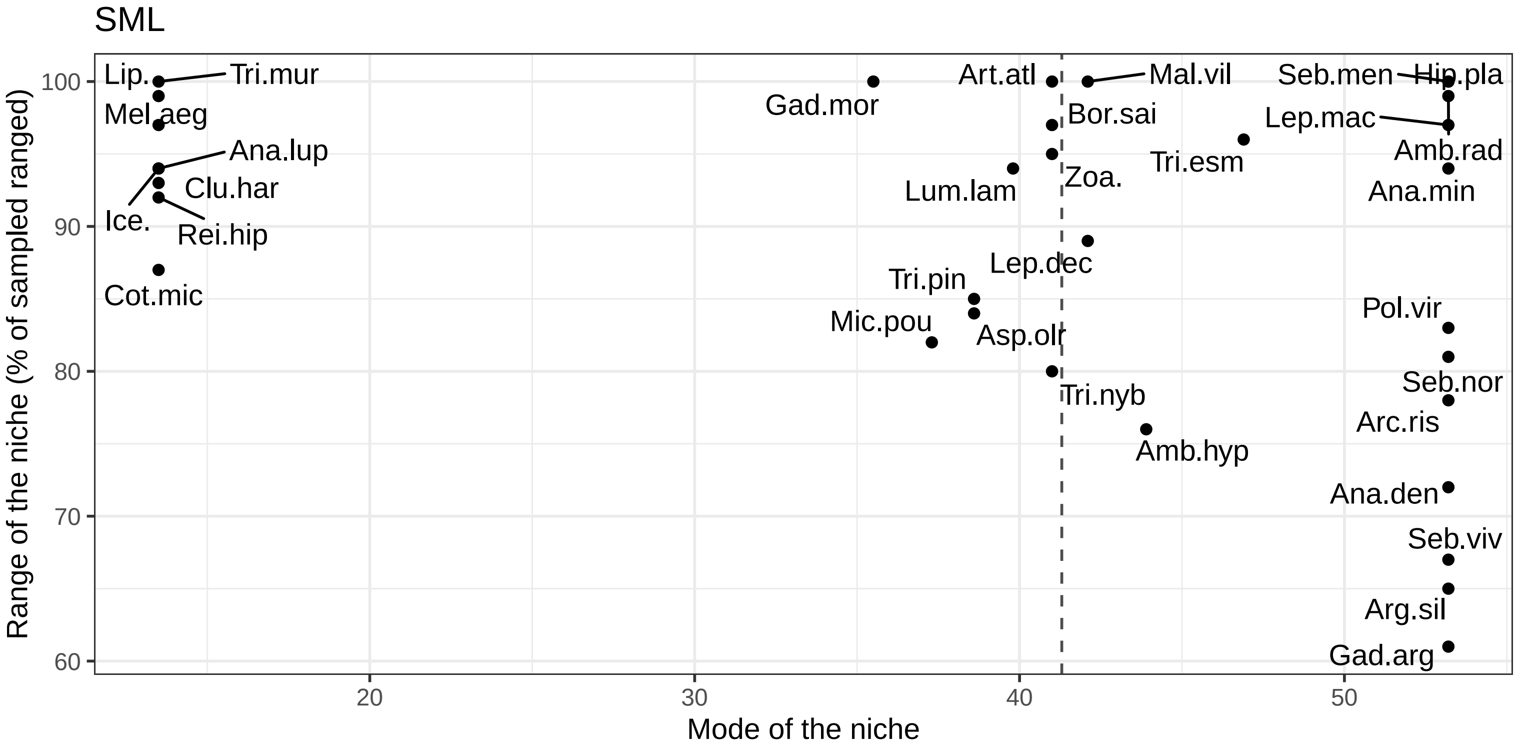


| Species | Code | Min | Max | Range | BS mean | Sp mean | Contrast | Data under the model |
|---|---|---|---|---|---|---|---|---|
| Anarhichas minor | Ana.min | 13.5 | 60.5 | 94 | 41.3 | 53.2 | 0.1202 | 98.73 |
| Amblyraja radiata | Amb.rad | 13.8 | 63.6 | 99 | 41.3 | 53.2 | 0.1343 | 99.61 |
| Leptoclinus maculatus | Lep.mac | 13.5 | 62.3 | 97 | 41.3 | 53.2 | 0.1391 | 98.63 |
| Micromesistius poutassou | Mic.pou | 18.9 | 60.2 | 82 | 41.3 | 37.3 | 0.1726 | 99.51 |
| Lumpenus lampretaeformis | Lum.lam | 13.5 | 60.5 | 94 | 41.3 | 39.8 | 0.2446 | 99.51 |
| Hippoglossoides platessoides | Hip.pla | 13.5 | 63.6 | 100 | 41.3 | 53.2 | 0.2451 | 99.41 |
| Reinhardtius hippoglossoides | Rei.hip | 16.2 | 62.3 | 92 | 41.3 | 13.5 | 0.2585 | 99.90 |
| Argentina silus | Arg.sil | 26.6 | 59.0 | 65 | 41.3 | 53.2 | 0.2749 | 98.04 |
| Anarhichas denticulatus | Ana.den | 24.2 | 60.3 | 72 | 41.3 | 53.2 | 0.4218 | 98.63 |
| Cottunculus microps | Cot.mic | 19.9 | 63.6 | 87 | 41.3 | 13.5 | 0.4572 | 99.61 |
| Zoarcidae | Zoa. | 13.7 | 61.1 | 95 | 41.3 | 41.0 | 0.4961 | 99.71 |
| Amblyraja hyperborea | Amb.hyp | 19.9 | 57.9 | 76 | 41.3 | 43.9 | 0.5019 | 99.61 |
| Leptagonus decagonus | Lep.dec | 15.9 | 60.3 | 89 | 41.3 | 42.1 | 0.5333 | 98.82 |
| Artediellus atlanticus | Art.atl | 13.5 | 63.6 | 100 | 41.3 | 41.0 | 0.5524 | 99.51 |
| Sebastes mentella | Seb.men | 13.7 | 63.6 | 100 | 41.3 | 53.2 | 0.5685 | 99.02 |
| Gadus morhua | Gad.mor | 13.7 | 63.6 | 100 | 41.3 | 35.5 | 0.5938 | 99.90 |
| Triglops murrayi | Tri.mur | 13.7 | 63.6 | 100 | 41.3 | 13.5 | 0.6209 | 98.53 |
| Triglops nybelini | Tri.nyb | 16.2 | 56.1 | 80 | 41.3 | 41.0 | 0.6569 | 99.12 |
| Sebastes viviparus | Seb.viv | 26.6 | 60.2 | 67 | 41.3 | 53.2 | 0.7116 | 99.02 |
| Arctozenus risso | Arc.ris | 19.6 | 58.5 | 78 | 41.3 | 53.2 | 0.7207 | 99.12 |
| Trisopterus esmarkii | Tri.esm | 13.7 | 61.7 | 96 | 41.3 | 46.9 | 0.7282 | 99.41 |
| Pollachius virens | Pol.vir | 20.0 | 61.6 | 83 | 41.3 | 53.2 | 0.7328 | 98.63 |
| Mallotus villosus | Mal.vil | 13.5 | 63.6 | 100 | 41.3 | 42.1 | 0.7574 | 99.12 |
| Anarhichas lupus | Ana.lup | 13.5 | 60.3 | 94 | 41.3 | 13.5 | 0.7709 | 98.24 |
| Icelus | Ice. | 13.5 | 60.3 | 94 | 41.3 | 13.5 | 0.7749 | 99.80 |
| Sebastes norvegicus | Seb.nor | 21.2 | 61.7 | 81 | 41.3 | 53.2 | 0.7972 | 98.33 |
| Liparidae | Lip. | 13.9 | 63.6 | 99 | 41.3 | 13.5 | 0.8019 | 99.80 |
| Melanogrammus aeglefinus | Mel.aeg | 13.7 | 62.3 | 97 | 41.3 | 13.5 | 0.8464 | 99.80 |
| Boreogadus saida | Bor.sai | 13.5 | 62.3 | 97 | 41.3 | 41.0 | 0.8951 | 99.41 |
| Clupea harengus | Clu.har | 13.5 | 60.2 | 93 | 41.3 | 13.5 | 0.9109 | 99.41 |
| Triglops pingelii | Tri.pin | 13.5 | 56.1 | 85 | 41.3 | 38.6 | 0.9110 | 98.63 |
| Gadiculus argenteus | Gad.arg | 28.5 | 59.0 | 61 | 41.3 | 53.2 | 0.9395 | 98.53 |
| Aspidophoroides olrikii | Asp.olr | 13.5 | 55.5 | 84 | 41.3 | 38.6 | 0.9581 | 98.63 |

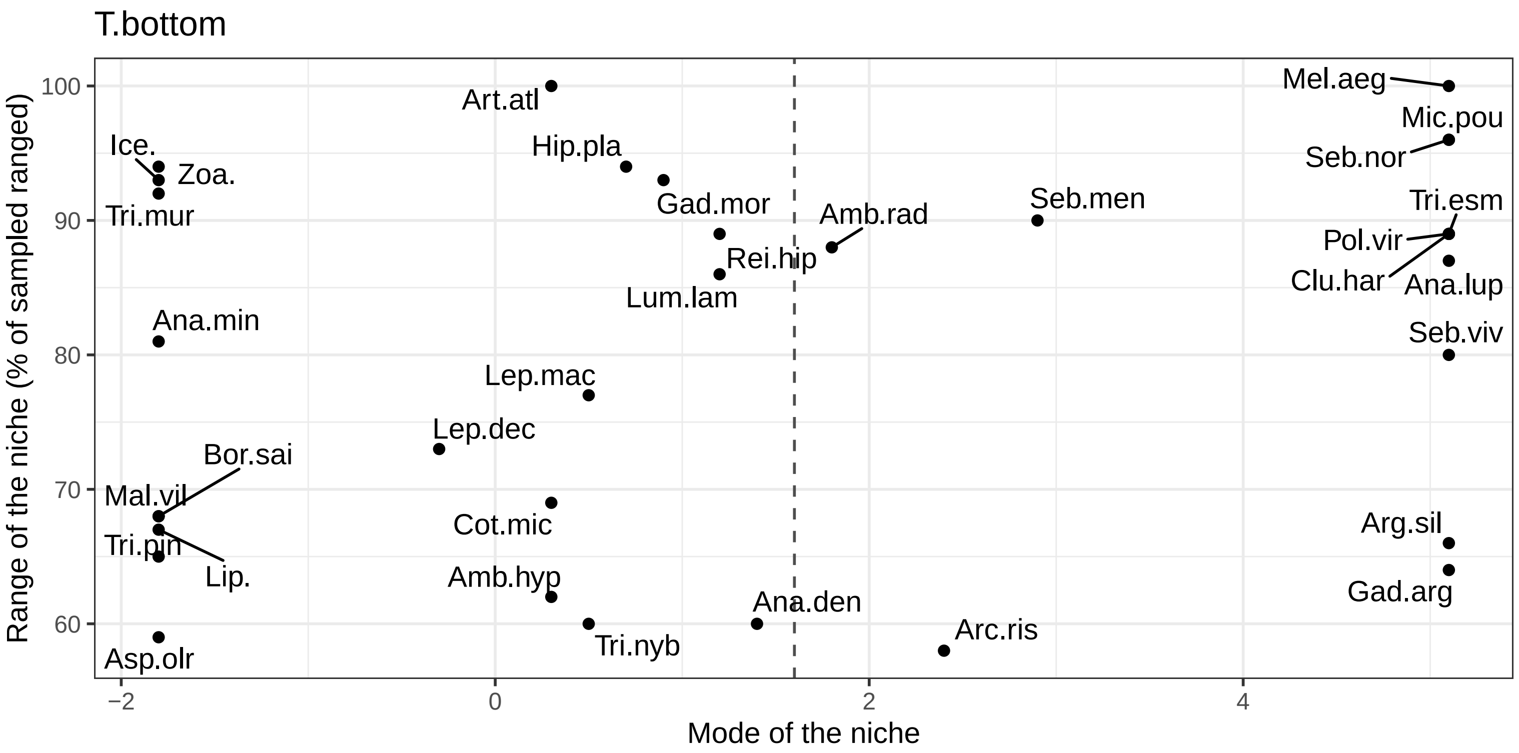


| Species | Code | Min | Max | Range | BS mean | Sp mean | Contrast | Data under the model |
|---|---|---|---|---|---|---|---|---|
| Anarhichas minor | Ana.min | −1.5 | 7.2 | 81 | 1.6 | −1.8 | 0.2713 | 98.92 |
| Lumpenus lampretaeformis | Lum.lam | −1.8 | 7.4 | 86 | 1.6 | 1.2 | 0.3861 | 99.51 |
| Anarhichas denticulatus | Ana.den | −1.0 | 5.5 | 60 | 1.6 | 1.4 | 0.4799 | 98.33 |
| Gadus morhua | Gad.mor | −1.8 | 8.2 | 93 | 1.6 | 0.9 | 0.6122 | 99.90 |
| Hippoglossoides platessoides | Hip.pla | −1.8 | 8.2 | 94 | 1.6 | 0.7 | 0.6749 | 99.51 |
| Cottunculus microps | Cot.mic | −0.7 | 6.8 | 69 | 1.6 | 0.3 | 0.7207 | 99.61 |
| Zoarcidae | Zoa. | −1.8 | 8.2 | 94 | 1.6 | −1.8 | 0.7225 | 99.61 |
| Amblyraja radiata | Amb.rad | −1.3 | 8.1 | 88 | 1.6 | 1.8 | 0.7561 | 99.61 |
| Reinhardtius hippoglossoides | Rei.hip | −1.3 | 8.2 | 89 | 1.6 | 1.2 | 0.7671 | 99.90 |
| Artediellus atlanticus | Art.atl | −1.8 | 8.9 | 100 | 1.6 | 0.3 | 0.8035 | 99.51 |
| Anarhichas lupus | Ana.lup | −1.2 | 8.2 | 87 | 1.6 | 5.1 | 0.8264 | 99.41 |
| Triglops murrayi | Tri.mur | −1.8 | 8.2 | 92 | 1.6 | −1.8 | 0.8524 | 98.33 |
| Leptoclinus maculatus | Lep.mac | −1.8 | 6.4 | 77 | 1.6 | 0.5 | 0.9588 | 98.73 |
| Leptagonus decagonus | Lep.dec | −1.8 | 6.0 | 73 | 1.6 | −0.3 | 0.9641 | 99.41 |
| Boreogadus saida | Bor.sai | −1.8 | 5.4 | 68 | 1.6 | −1.8 | 0.9717 | 99.90 |
| Mallotus villosus | Mal.vil | −1.8 | 5.5 | 68 | 1.6 | −1.8 | 0.9835 | 99.22 |
| Clupea harengus | Clu.har | −1.8 | 7.7 | 89 | 1.6 | 5.1 | 0.9863 | 99.41 |
| Arctozenus risso | Arc.ris | −0.7 | 5.6 | 58 | 1.6 | 2.4 | 0.9944 | 99.31 |
| Icelus | Ice. | −1.8 | 8.1 | 93 | 1.6 | −1.8 | 0.9957 | 99.90 |
| Triglops pingelii | Tri.pin | −1.8 | 5.2 | 65 | 1.6 | −1.8 | 0.9979 | 99.51 |
| Liparidae | Lip. | −1.8 | 5.3 | 67 | 1.6 | −1.8 | 0.9984 | 99.90 |
| Sebastes mentella | Seb.men | −1.4 | 8.2 | 90 | 1.6 | 2.9 | 0.9992 | 98.04 |
| Sebastes norvegicus | Seb.nor | −1.4 | 8.9 | 96 | 1.6 | 5.1 | 0.9992 | 99.41 |
| Amblyraja hyperborea | Amb.hyp | −1.8 | 4.9 | 62 | 1.6 | 0.3 | 0.9993 | 98.82 |
| Melanogrammus aeglefinus | Mel.aeg | −1.8 | 8.9 | 100 | 1.6 | 5.1 | 0.9993 | 99.71 |
| Aspidophoroides olrikii | Asp.olr | −1.8 | 4.5 | 59 | 1.6 | −1.8 | 0.9996 | 99.02 |
| Triglops nybelini | Tri.nyb | −1.8 | 4.6 | 60 | 1.6 | 0.5 | 0.9996 | 99.41 |
| Gadiculus argenteus | Gad.arg | 2.0 | 8.9 | 64 | 1.6 | 5.1 | 0.9999 | 99.41 |
| Argentina silus | Arg.sil | 1.8 | 8.9 | 66 | 1.6 | 5.1 | 1.0000 | 98.63 |
| Micromesistius poutassou | Mic.pou | −1.4 | 8.9 | 96 | 1.6 | 5.1 | 1.0000 | 99.41 |
| Pollachius virens | Pol.vir | −0.6 | 8.9 | 89 | 1.6 | 5.1 | 1.0000 | 99.22 |
| Sebastes viviparus | Seb.viv | 0.3 | 8.9 | 80 | 1.6 | 5.1 | 1.0000 | 99.71 |
| Trisopterus esmarkii | Tri.esm | −0.6 | 8.9 | 89 | 1.6 | 5.1 | 1.0000 | 99.80 |

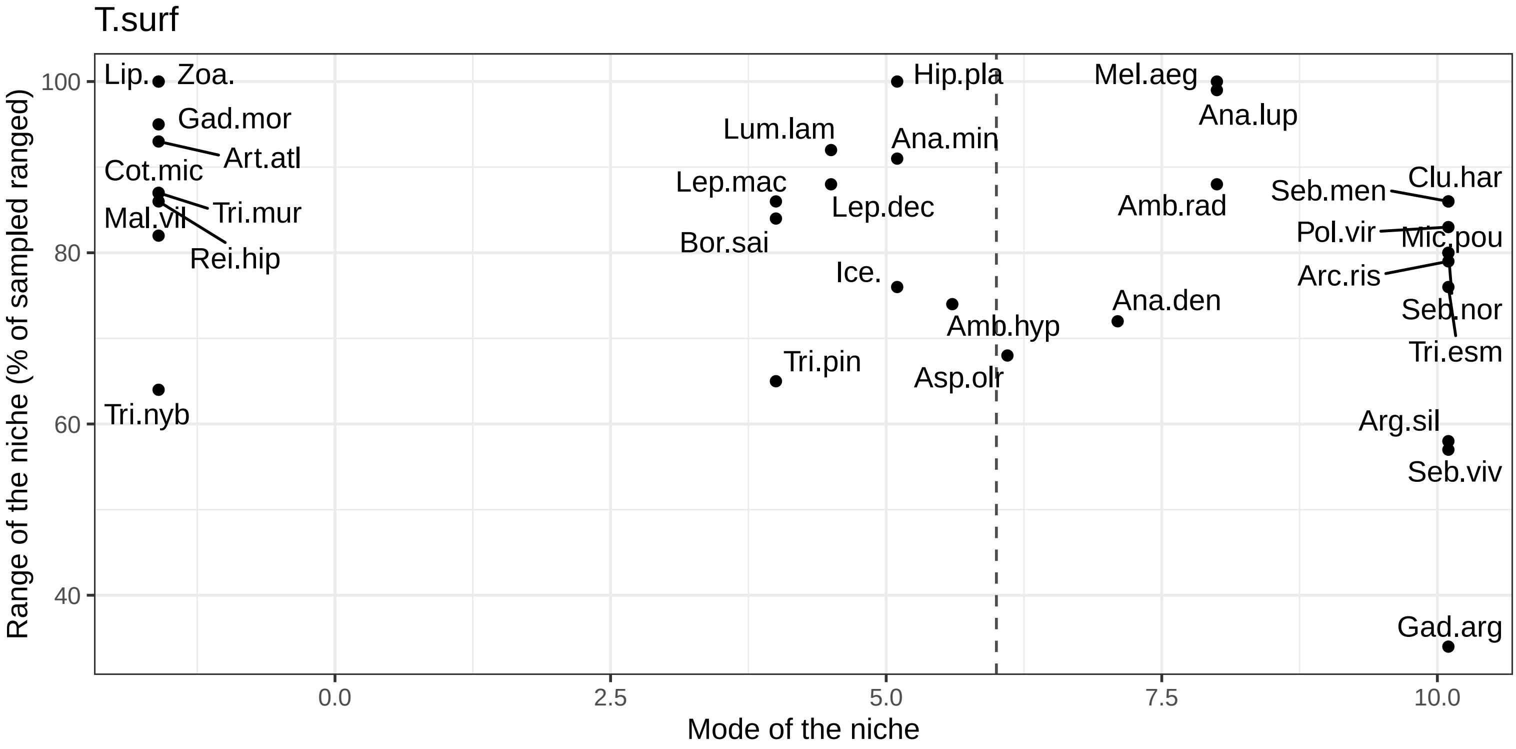


| Species | Code | Min | Max | Range | BS mean | Sp mean | Contrast | Data under the model |
|---|---|---|---|---|---|---|---|---|
| Anarhichas minor | Ana.min | −0.2 | 14.5 | 91 | 6 | 5.1 | 0.3943 | 98.82 |
| Reinhardtius hippoglossoides | Rei.hip | −1.6 | 12.3 | 86 | 6 | −1.6 | 0.6547 | 99.80 |
| Hippoglossoides platessoides | Hip.pla | −1.6 | 14.5 | 100 | 6 | 5.1 | 0.7265 | 99.51 |
| Zoarcidae | Zoa. | −1.6 | 14.5 | 100 | 6 | −1.6 | 0.7455 | 99.90 |
| Anarhichas denticulatus | Ana.den | 0.4 | 12.1 | 72 | 6 | 7.1 | 0.7761 | 98.43 |
| Lumpenus lampretaeformis | Lum.lam | −0.4 | 14.5 | 92 | 6 | 4.5 | 0.7857 | 99.61 |
| Amblyraja radiata | Amb.rad | −1.6 | 12.6 | 88 | 6 | 8.0 | 0.7885 | 98.92 |
| Gadus morhua | Gad.mor | −0.9 | 14.5 | 95 | 6 | −1.6 | 0.8273 | 99.90 |
| Arctozenus risso | Arc.ris | −1.1 | 11.7 | 79 | 6 | 10.1 | 0.8359 | 99.22 |
| Amblyraja hyperborea | Amb.hyp | −1.6 | 10.4 | 74 | 6 | 5.6 | 0.8482 | 99.22 |
| Anarhichas lupus | Ana.lup | −1.5 | 14.5 | 99 | 6 | 8.0 | 0.8937 | 98.63 |
| Artediellus atlanticus | Art.atl | −1.6 | 13.4 | 93 | 6 | −1.6 | 0.9169 | 99.61 |
| Melanogrammus aeglefinus | Mel.aeg | −1.6 | 14.5 | 100 | 6 | 8.0 | 0.9279 | 99.80 |
| Leptoclinus maculatus | Lep.mac | −1.6 | 12.3 | 86 | 6 | 4.0 | 0.9315 | 98.63 |
| Leptagonus decagonus | Lep.dec | −1.6 | 12.6 | 88 | 6 | 4.5 | 0.9323 | 98.82 |
| Cottunculus microps | Cot.mic | −1.5 | 12.6 | 87 | 6 | −1.6 | 0.9377 | 99.61 |
| Triglops murrayi | Tri.mur | −1.6 | 12.4 | 87 | 6 | −1.6 | 0.9407 | 98.73 |
| Sebastes mentella | Seb.men | −1.6 | 12.3 | 86 | 6 | 10.1 | 0.9785 | 99.02 |
| Mallotus villosus | Mal.vil | −1.6 | 11.7 | 82 | 6 | −1.6 | 0.9861 | 99.61 |
| Sebastes norvegicus | Seb.nor | 0.6 | 13.4 | 80 | 6 | 10.1 | 0.9922 | 98.63 |
| Aspidophoroides olrikii | Asp.olr | 1.0 | 12.0 | 68 | 6 | 6.1 | 0.9924 | 98.43 |
| Icelus | Ice. | −1.6 | 10.7 | 76 | 6 | 5.1 | 0.9942 | 99.71 |
| Boreogadus saida | Bor.sai | −1.6 | 12.0 | 84 | 6 | 4.0 | 0.9948 | 99.71 |
| Liparidae | Lip. | −1.6 | 14.5 | 100 | 6 | −1.6 | 0.9968 | 100.00 |
| Clupea harengus | Clu.har | 0.7 | 14.5 | 86 | 6 | 10.1 | 0.9984 | 99.02 |
| Micromesistius poutassou | Mic.pou | 0.5 | 13.4 | 80 | 6 | 10.1 | 0.9993 | 99.41 |
| Triglops pingelii | Tri.pin | −1.3 | 9.2 | 65 | 6 | 4.0 | 0.9997 | 99.22 |
| Argentina silus | Arg.sil | 4.0 | 13.4 | 58 | 6 | 10.1 | 1.0000 | 99.61 |
| Gadiculus argenteus | Gad.arg | 6.8 | 12.4 | 34 | 6 | 10.1 | 1.0000 | 99.90 |
| Pollachius virens | Pol.vir | 1.1 | 14.5 | 83 | 6 | 10.1 | 1.0000 | 98.73 |
| Sebastes viviparus | Seb.viv | 4.2 | 13.4 | 57 | 6 | 10.1 | 1.0000 | 99.71 |
| Triglops nybelini | Tri.nyb | −1.6 | 8.7 | 64 | 6 | −1.6 | 1.0000 | 100.00 |
| Trisopterus esmarkii | Tri.esm | 2.3 | 14.5 | 76 | 6 | 10.1 | 1.0000 | 99.80 |
