## Appendix 7 Habitat suitability maps for "Suitable habitats of fish species in the Barents Sea"

### Amblyraja hyperborea

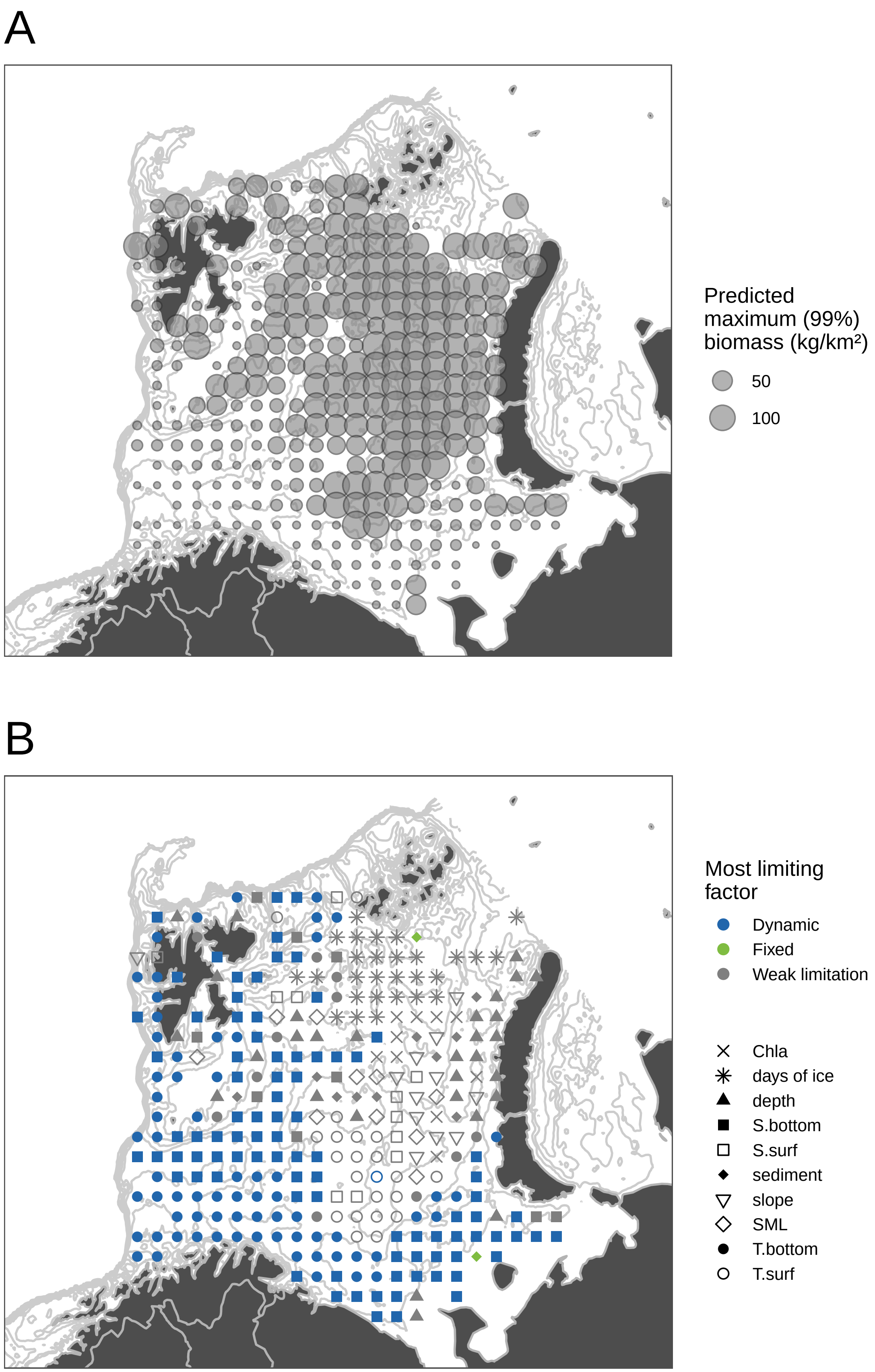

### *Amblyraja radiata*

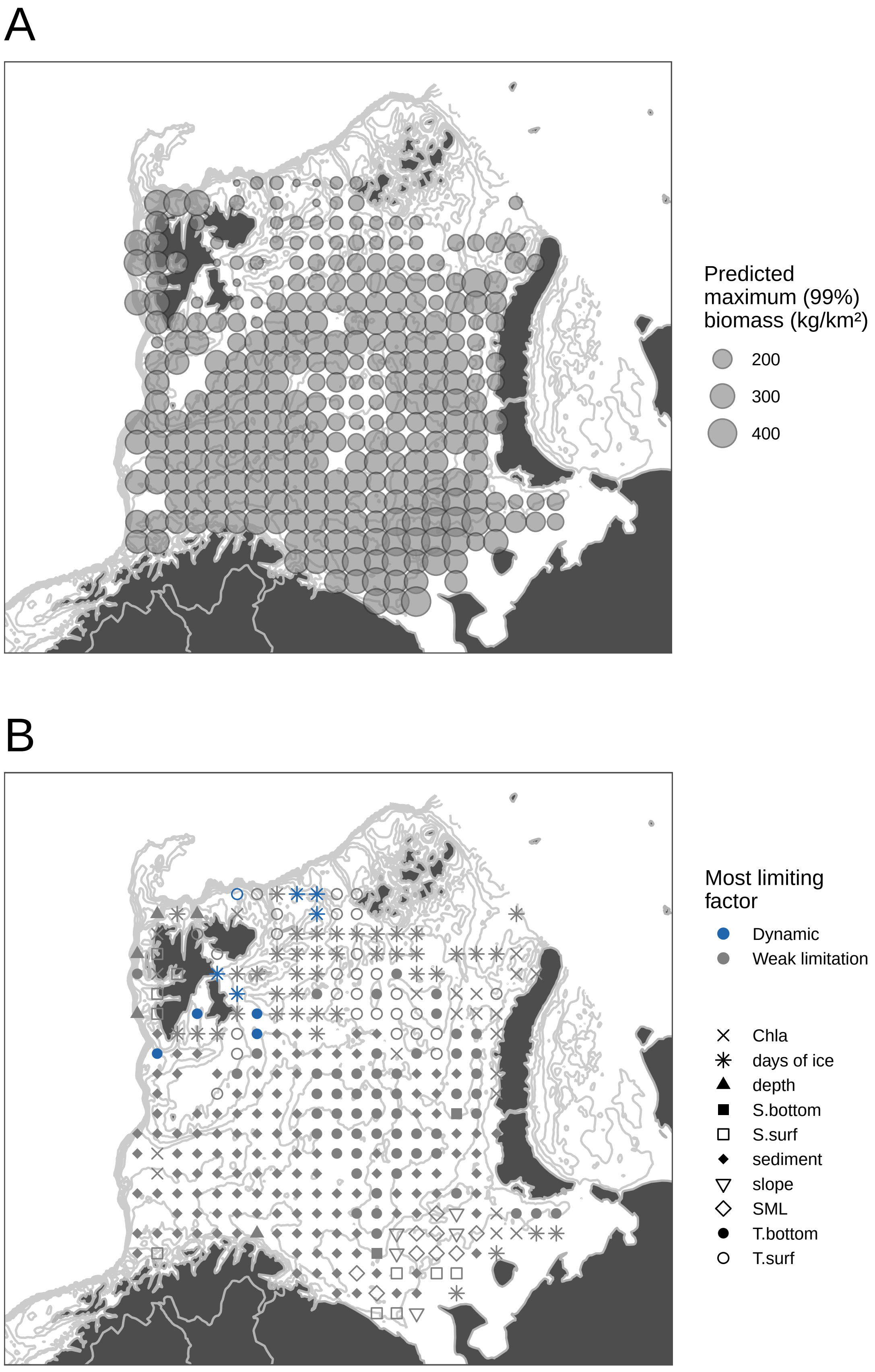

### Anarhichas denticulatus

A

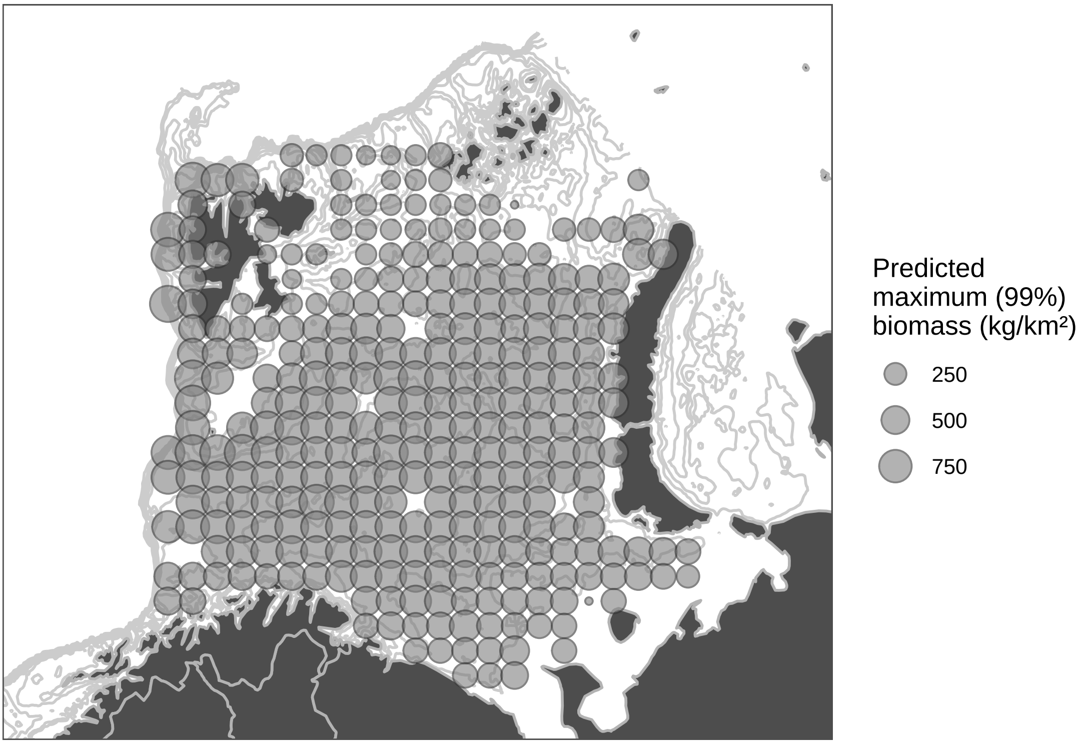


B

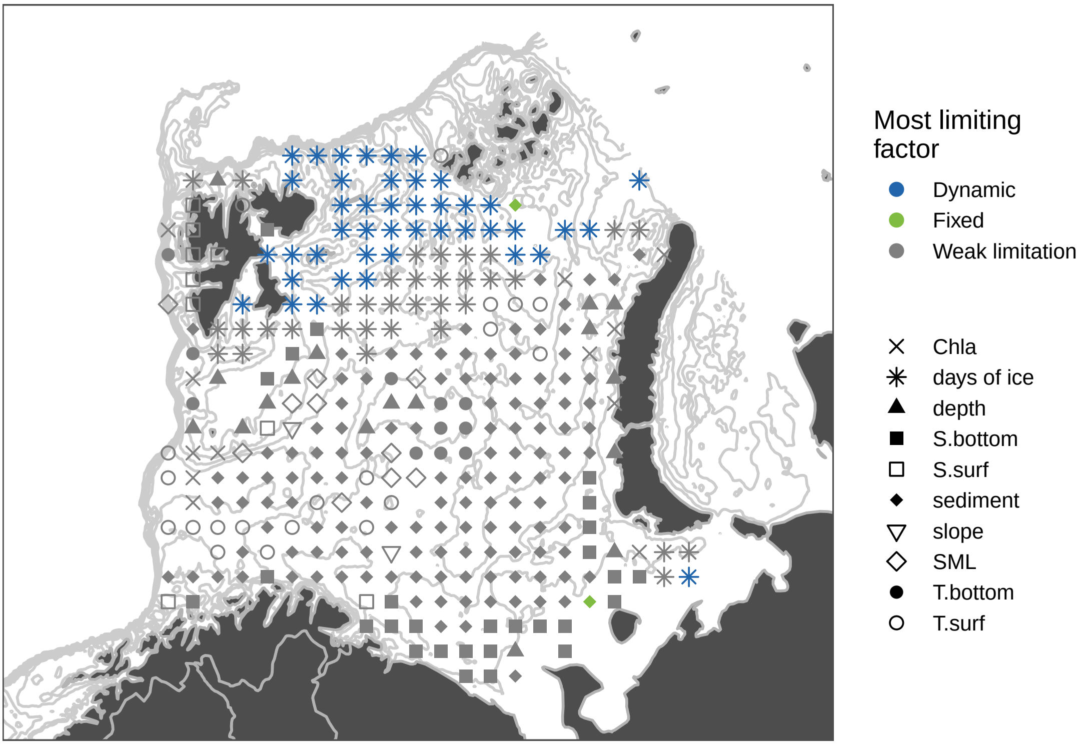

### Anarhichas lupus

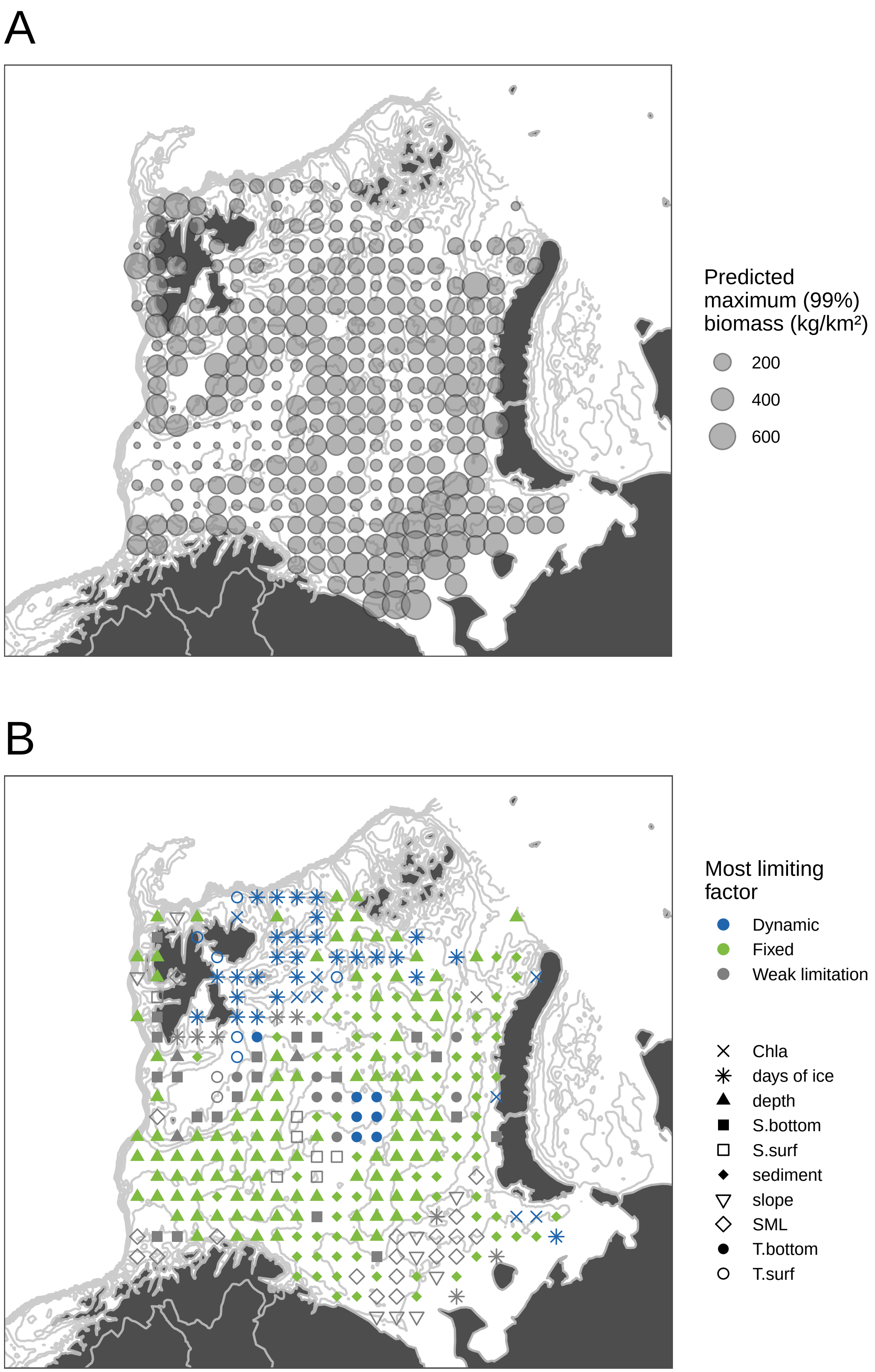

### Anarhichas minor

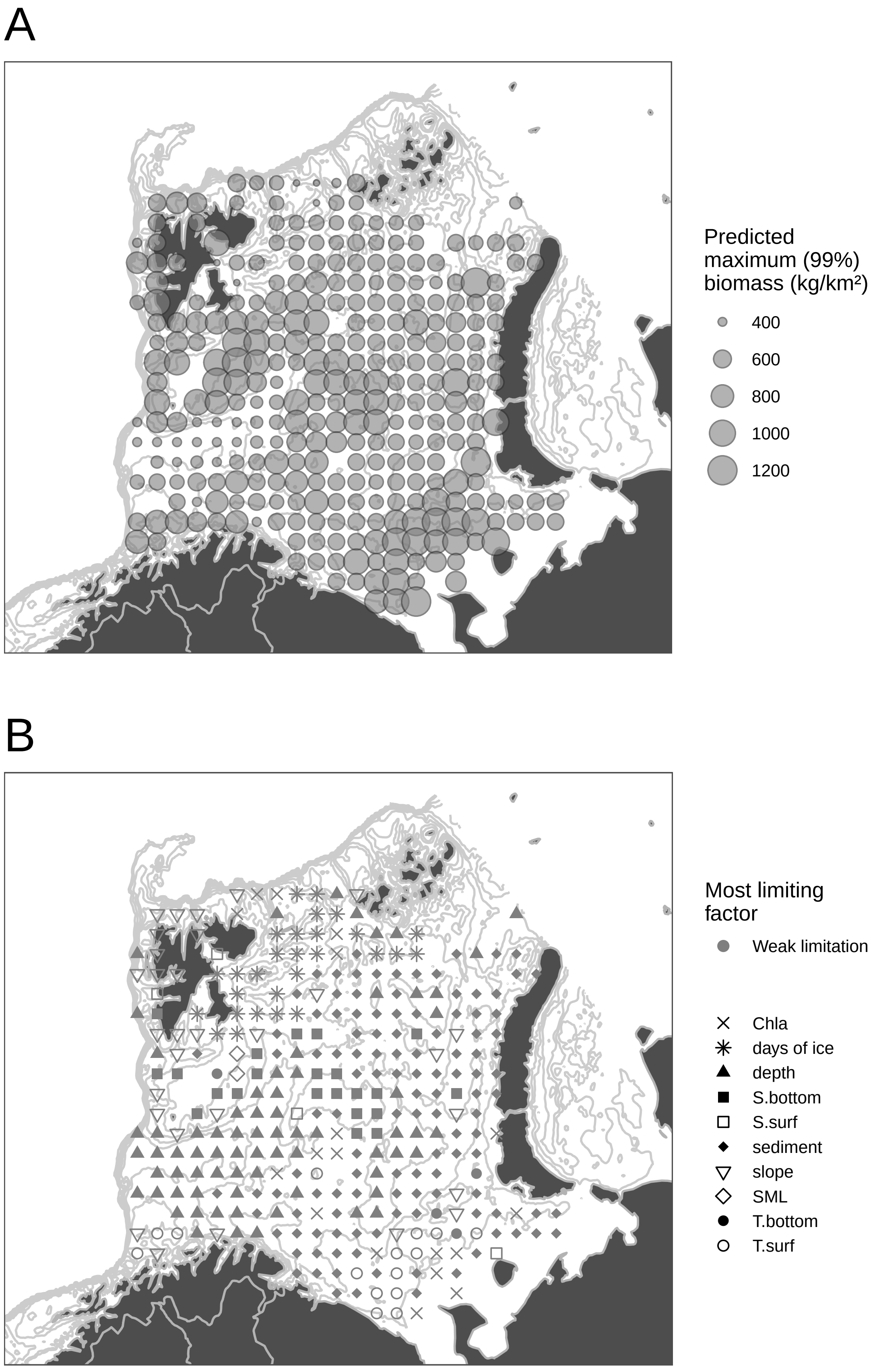

### *Arctozenus risso*

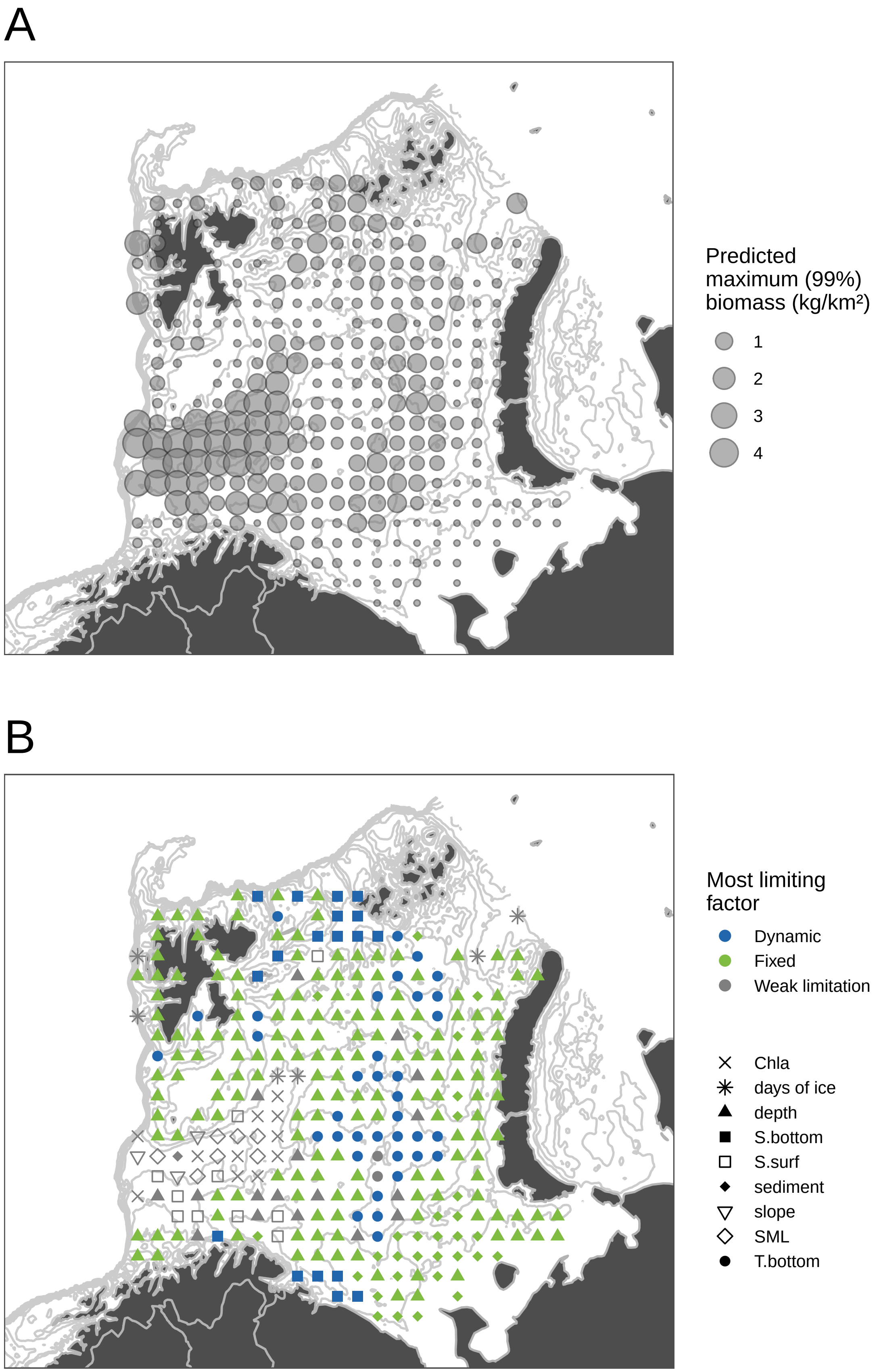

#### Argentina silus

A

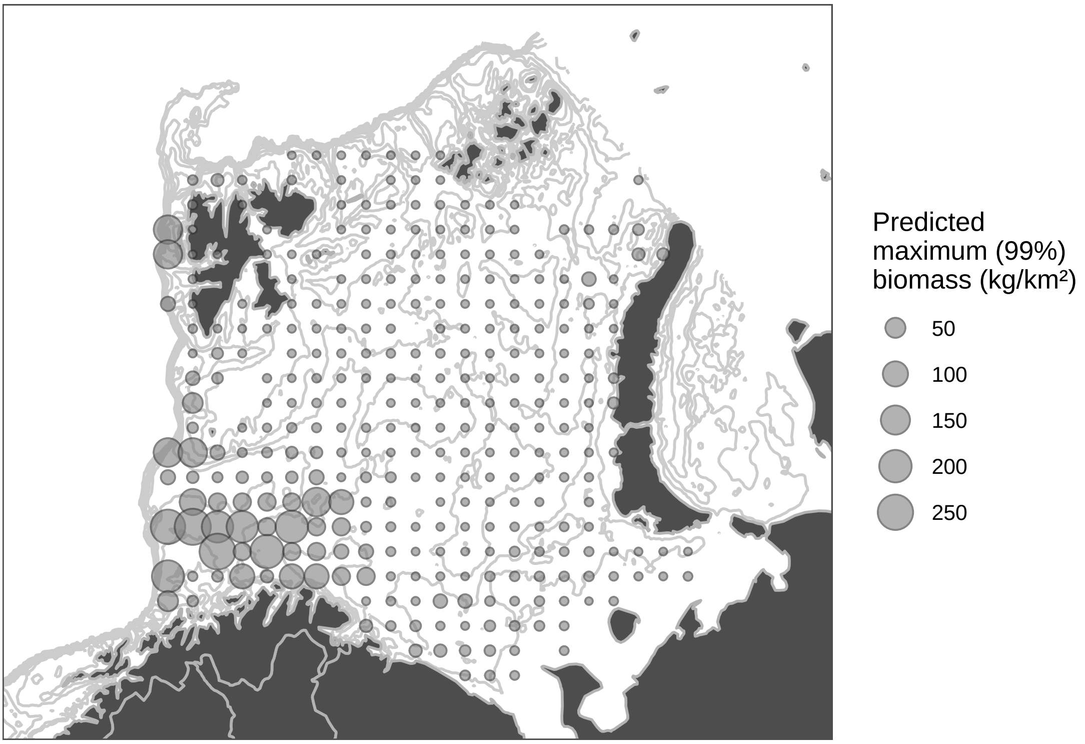


B

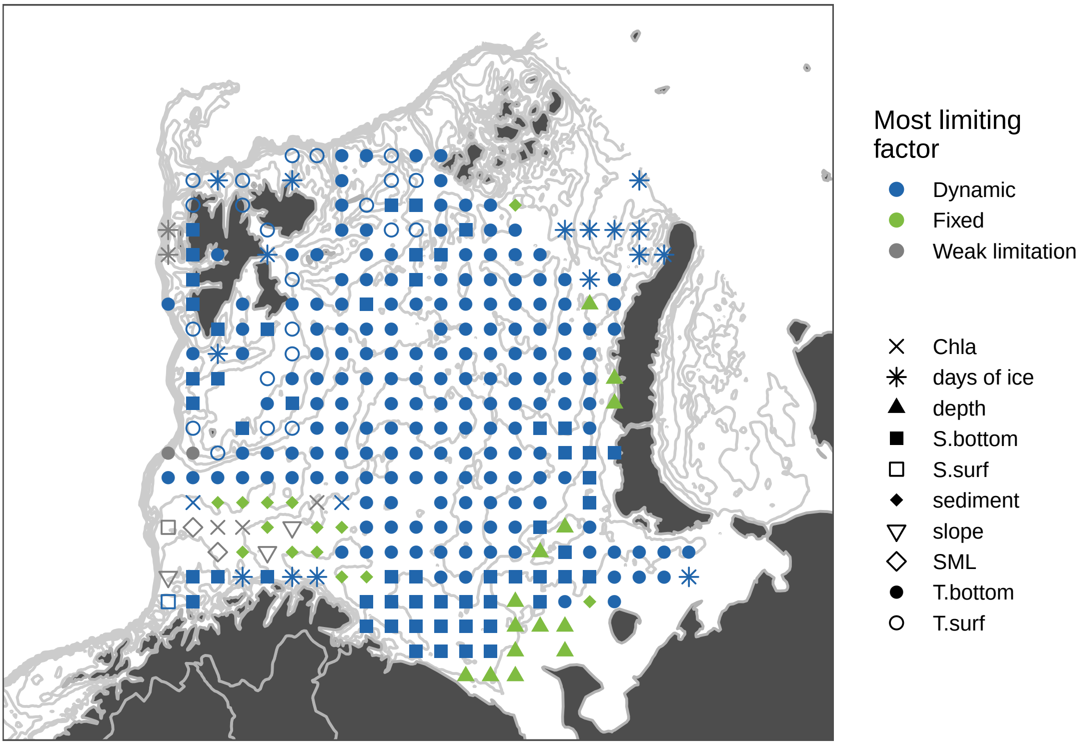

### Artediellus atlanticus

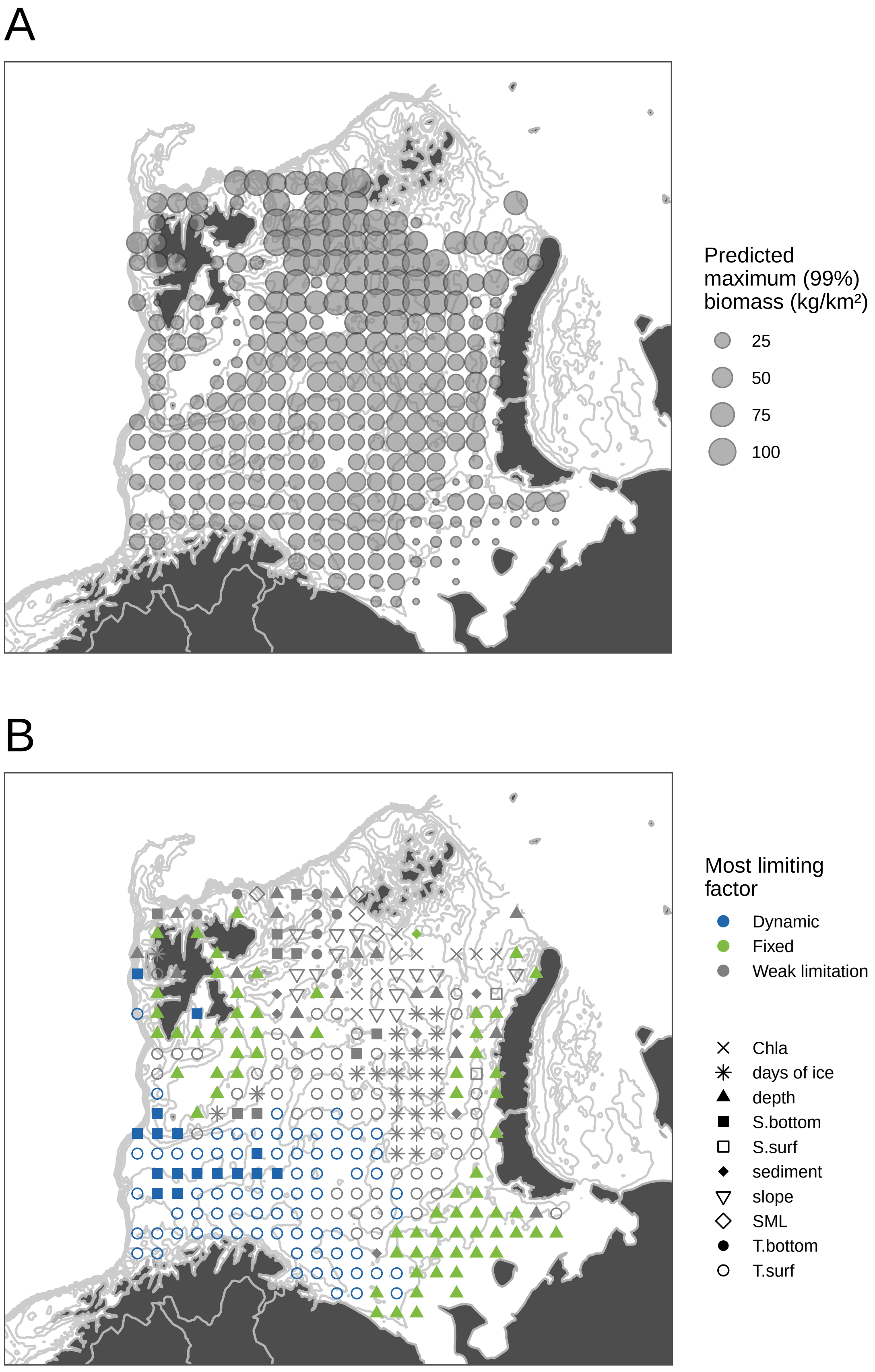

### Aspidophoroides olrikii

A

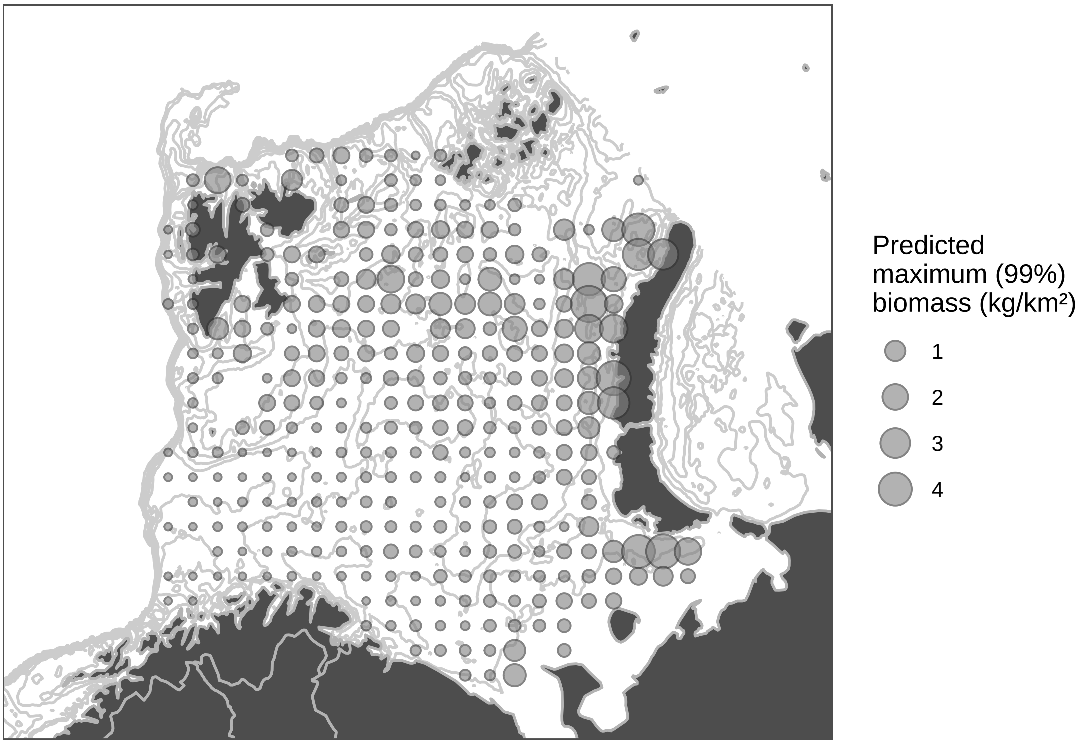


B

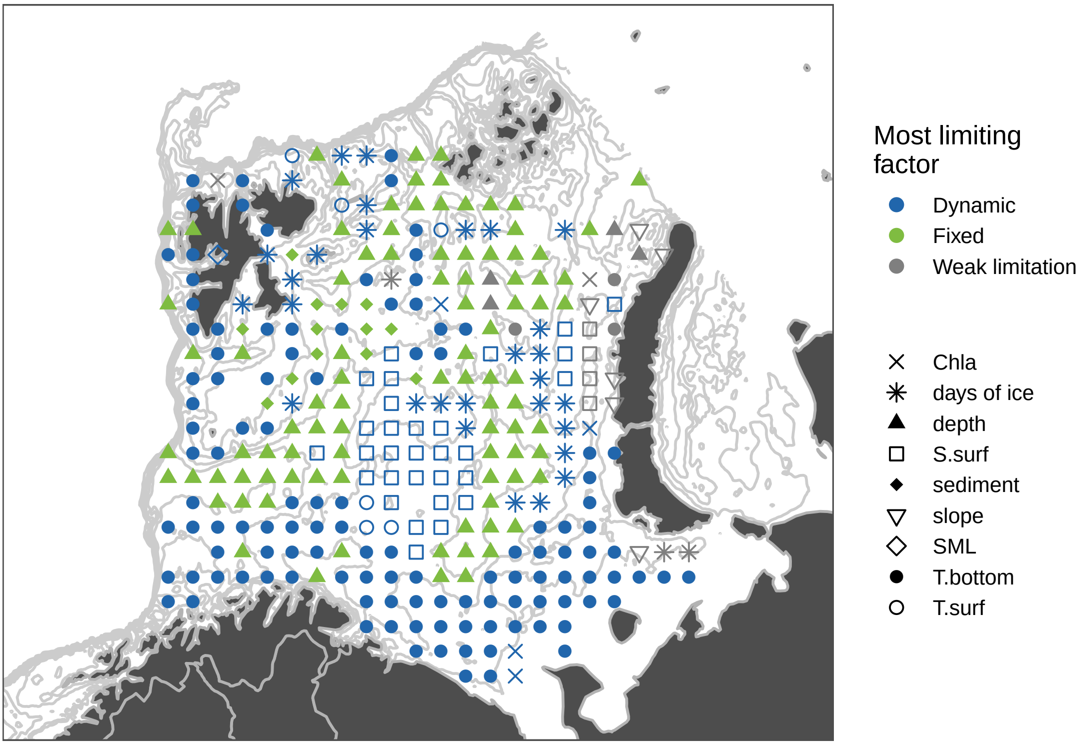

### Boreogadus saida

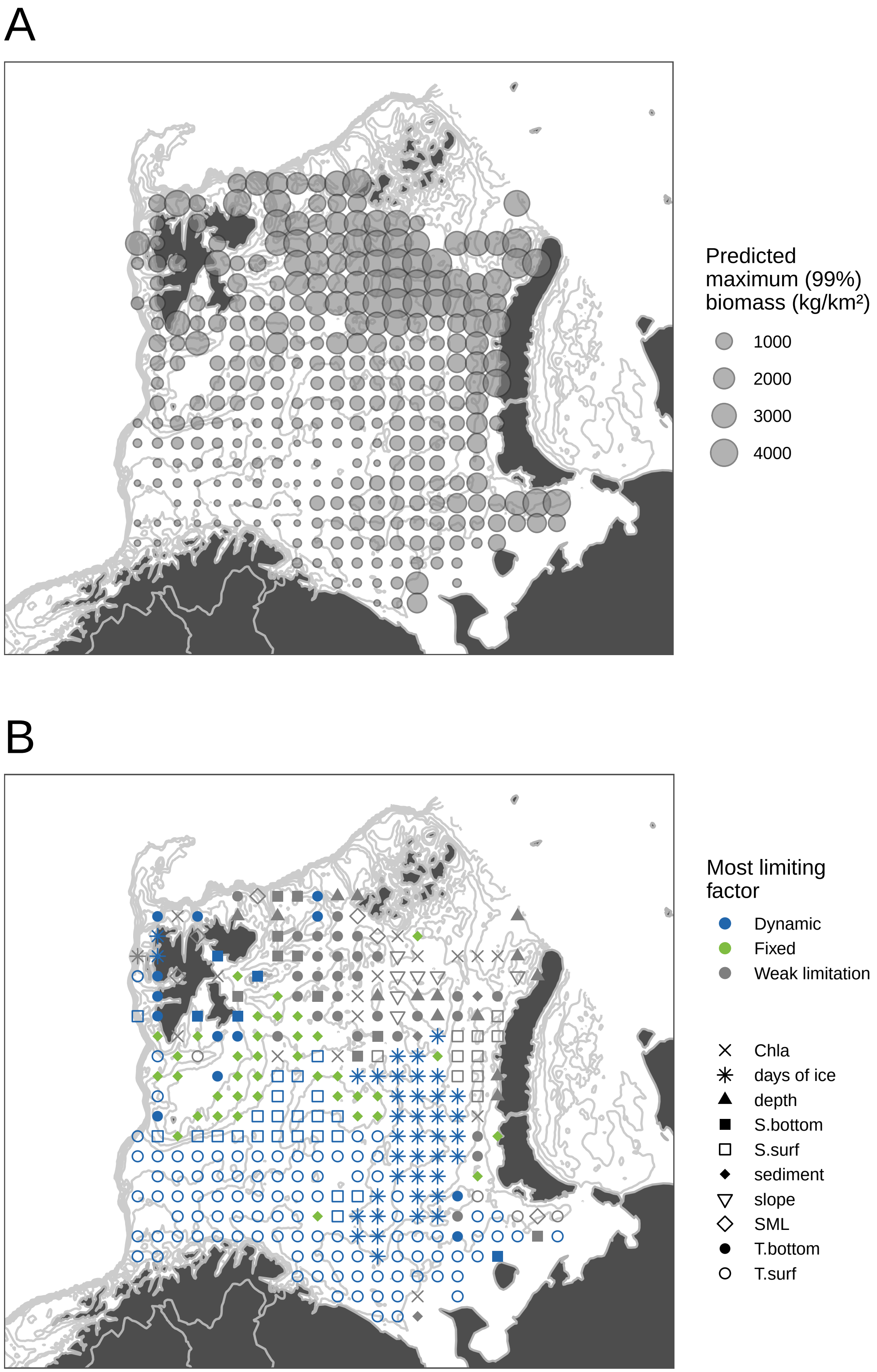

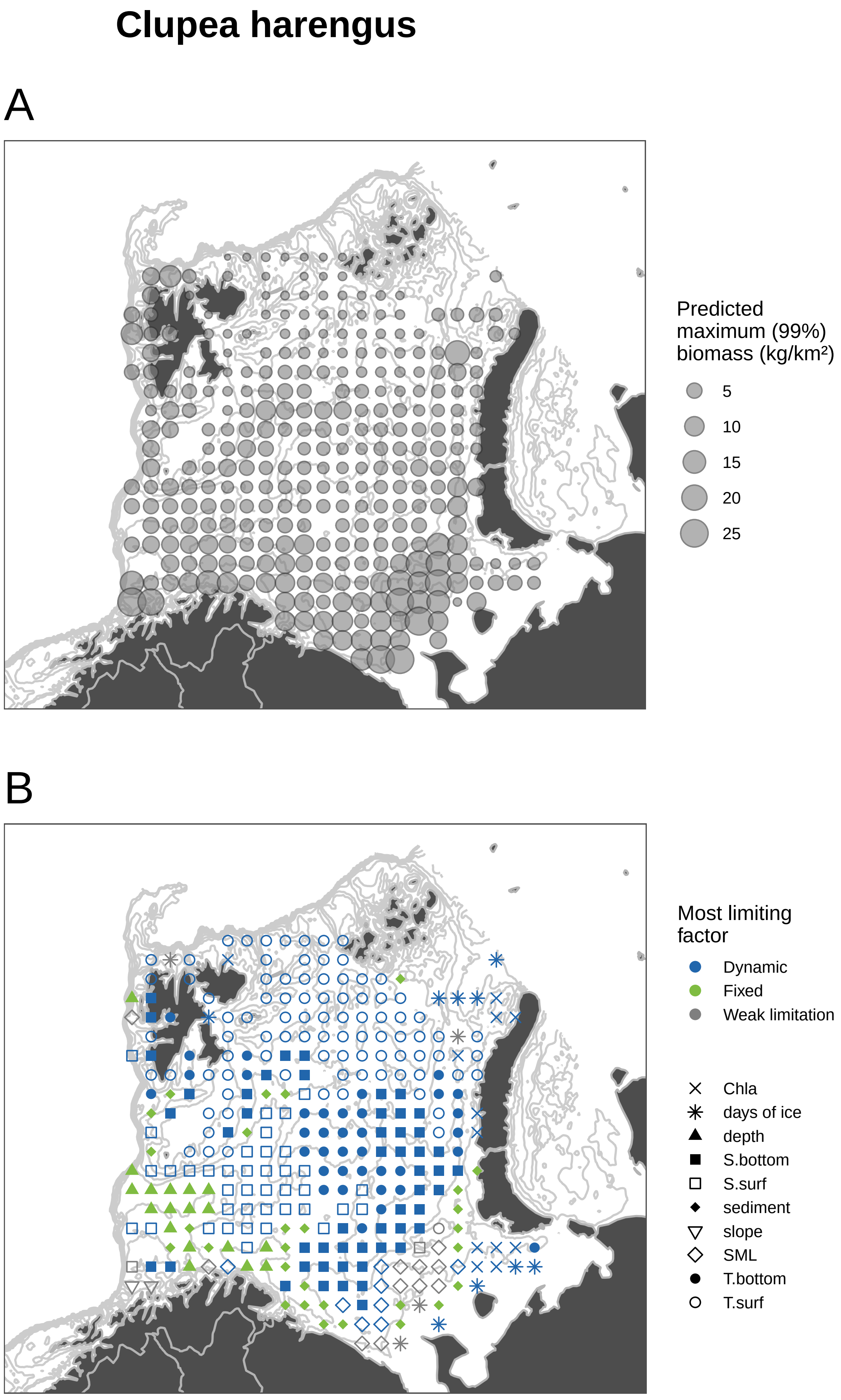
Clupea harengus
A
Predicted maximum (99%) biomass (kg/km²)
5
10
15
20
25
B
Most limiting factor
Dynamic
Fixed
Weak limitation
Chla
days of ice
depth
S.bottom
S.surf
sediment
slope
SML
T.bottom
T.surf

### Cottunculus microps

A

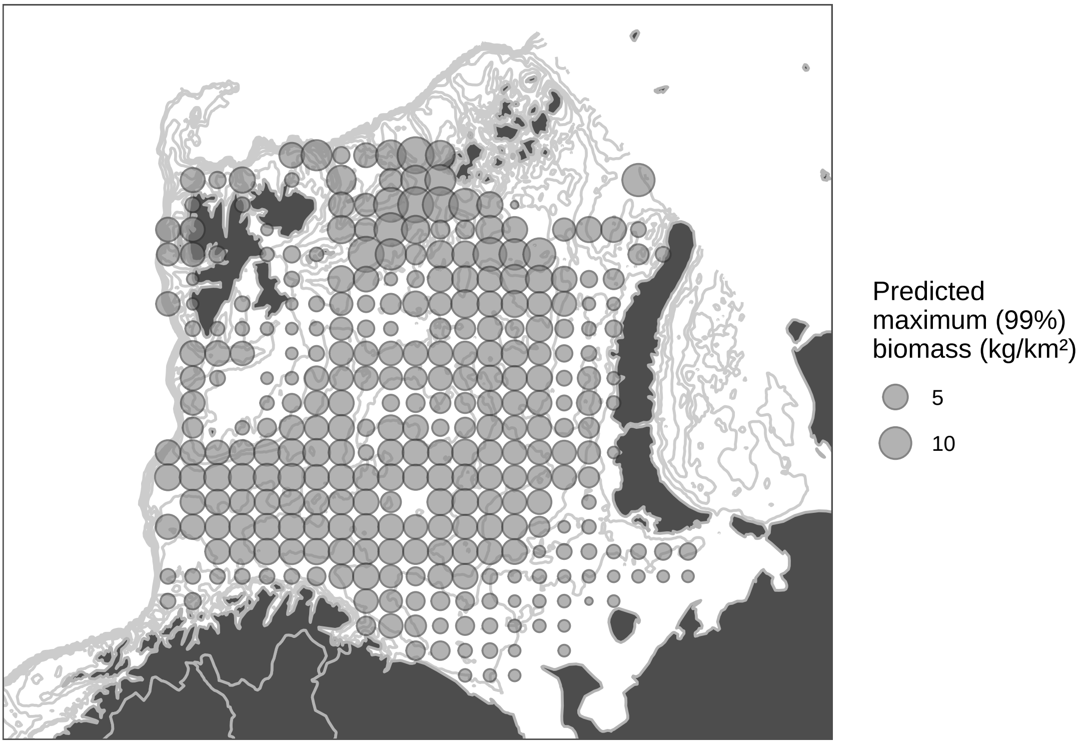


B

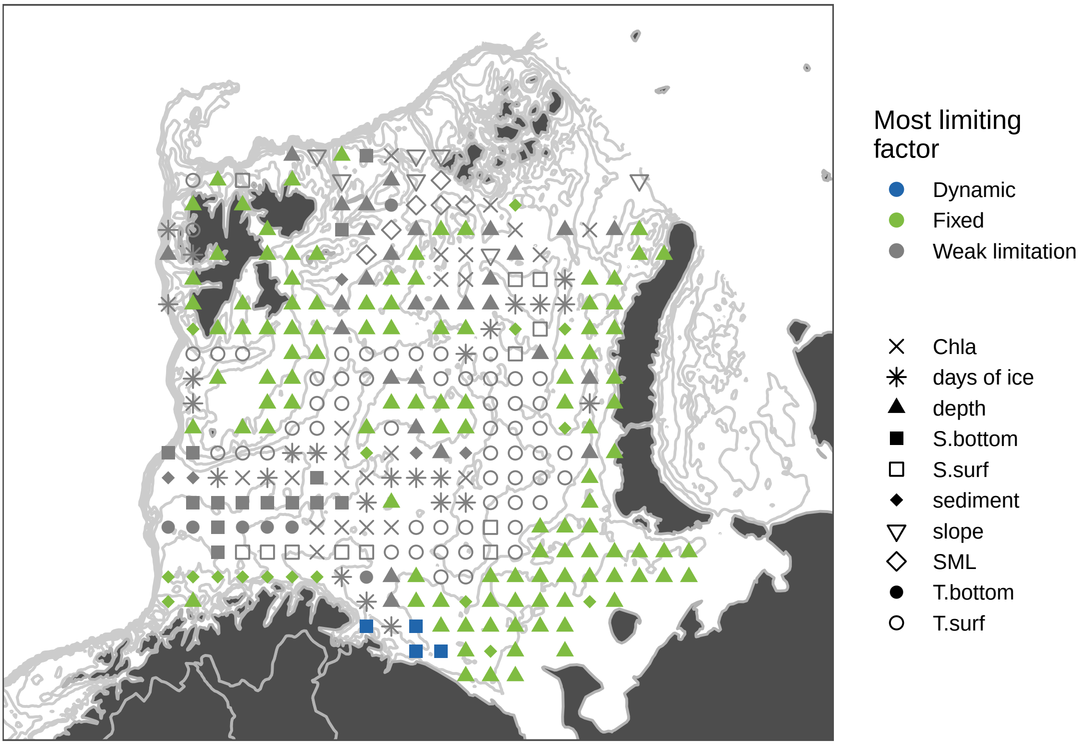

### *Gadiculus argenteus*

A

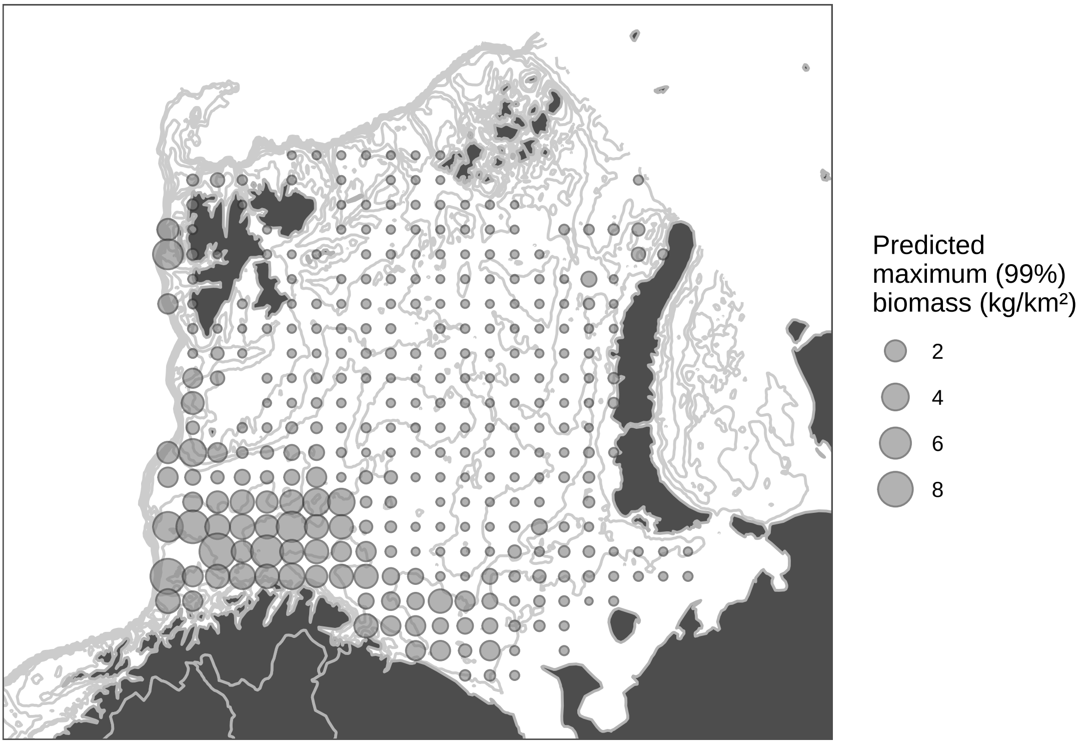


B

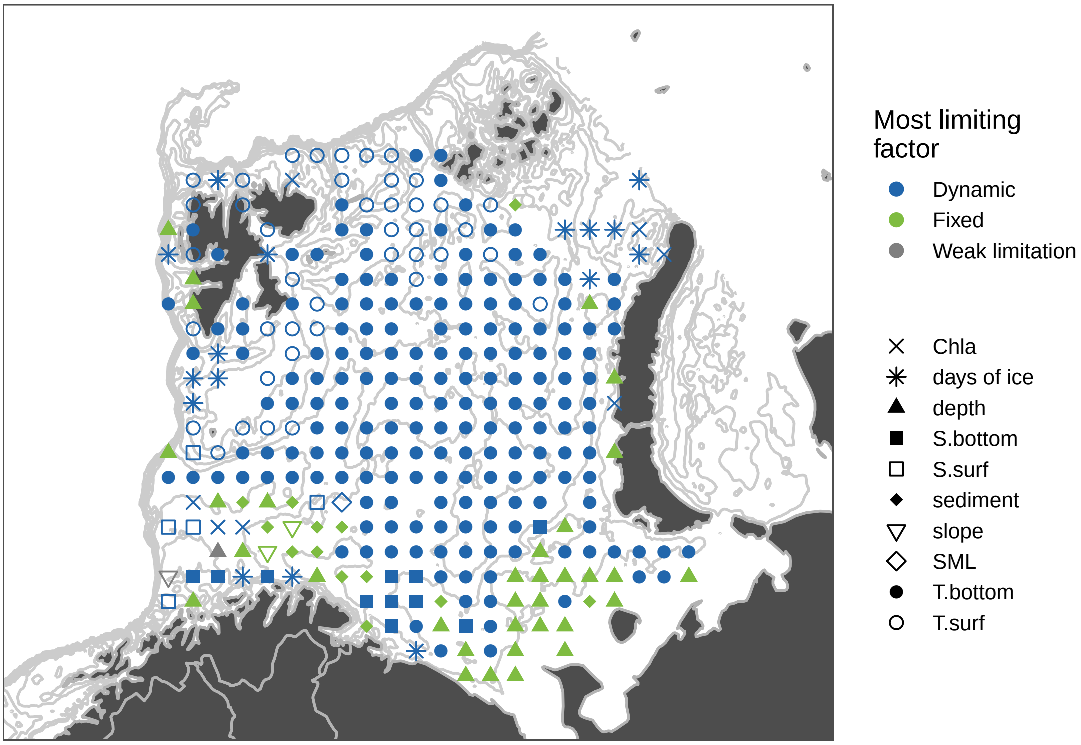

### Gadus morhua

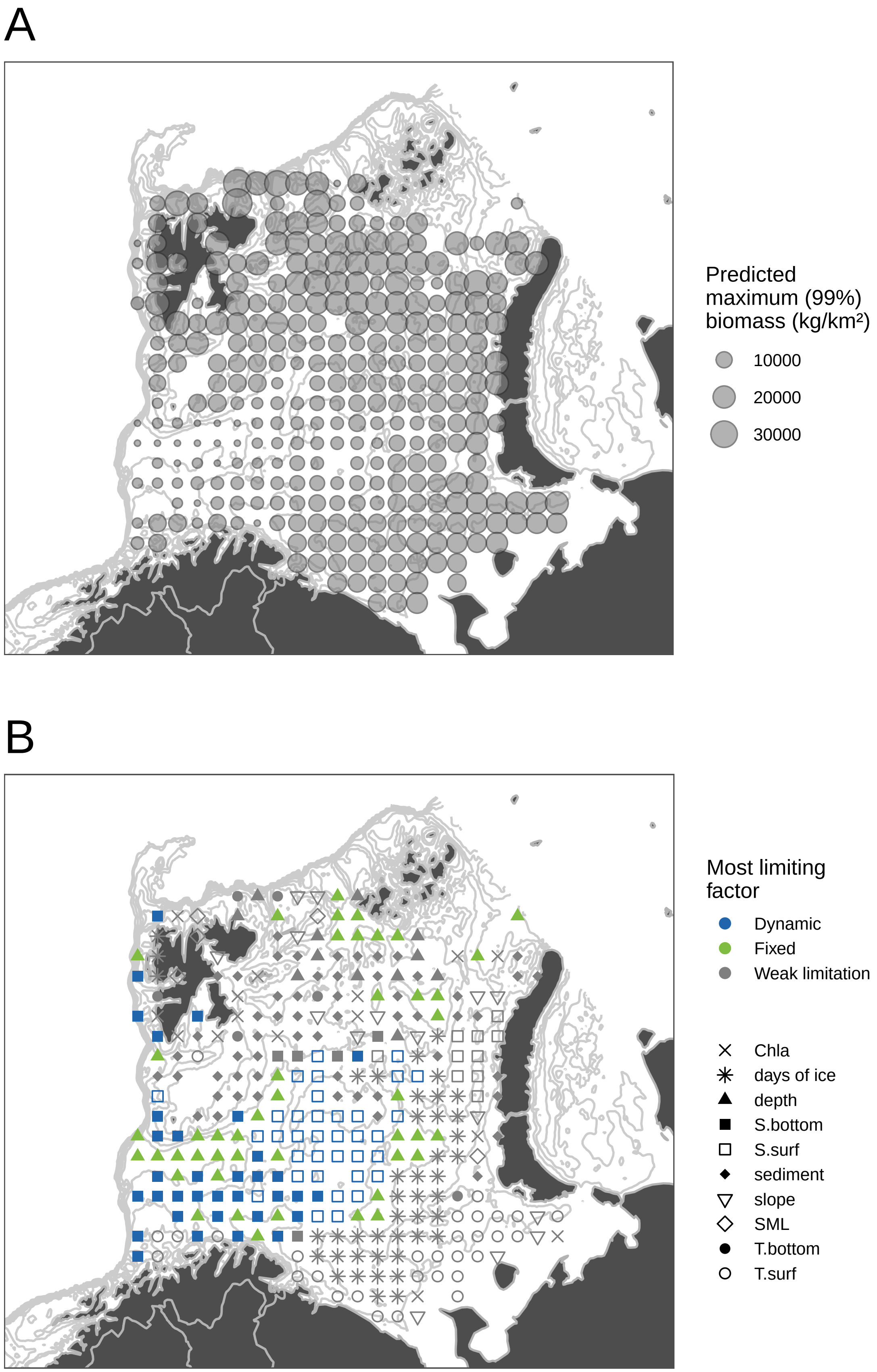

### Hippoglossoides platessoides

A

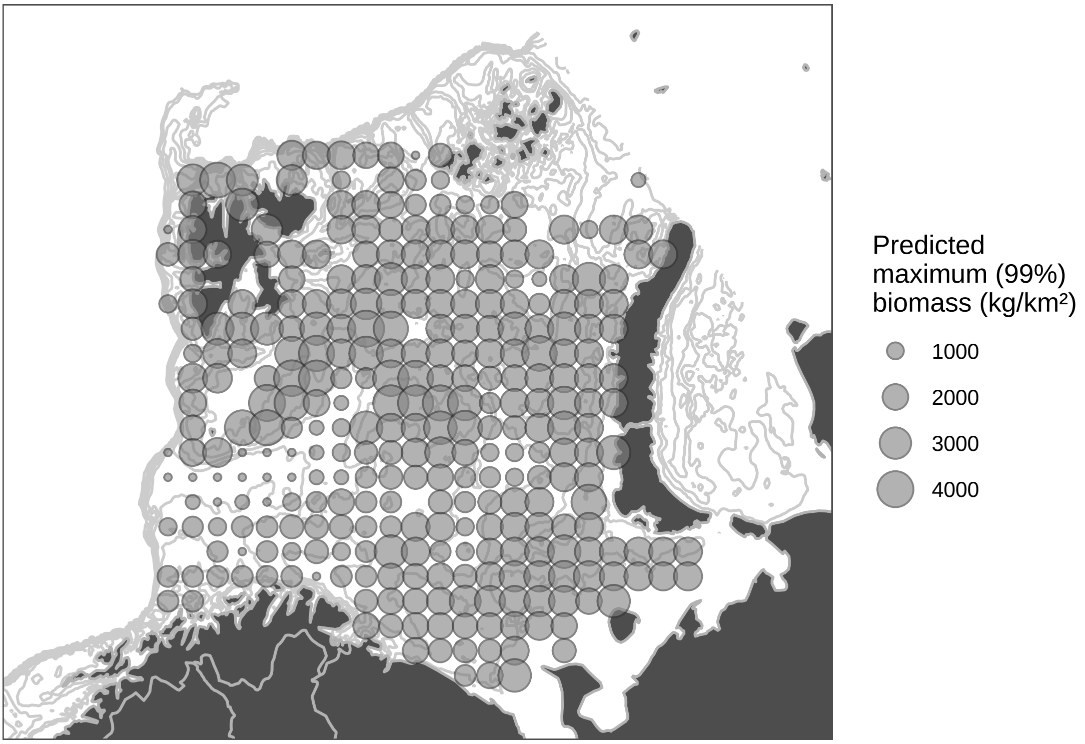


B

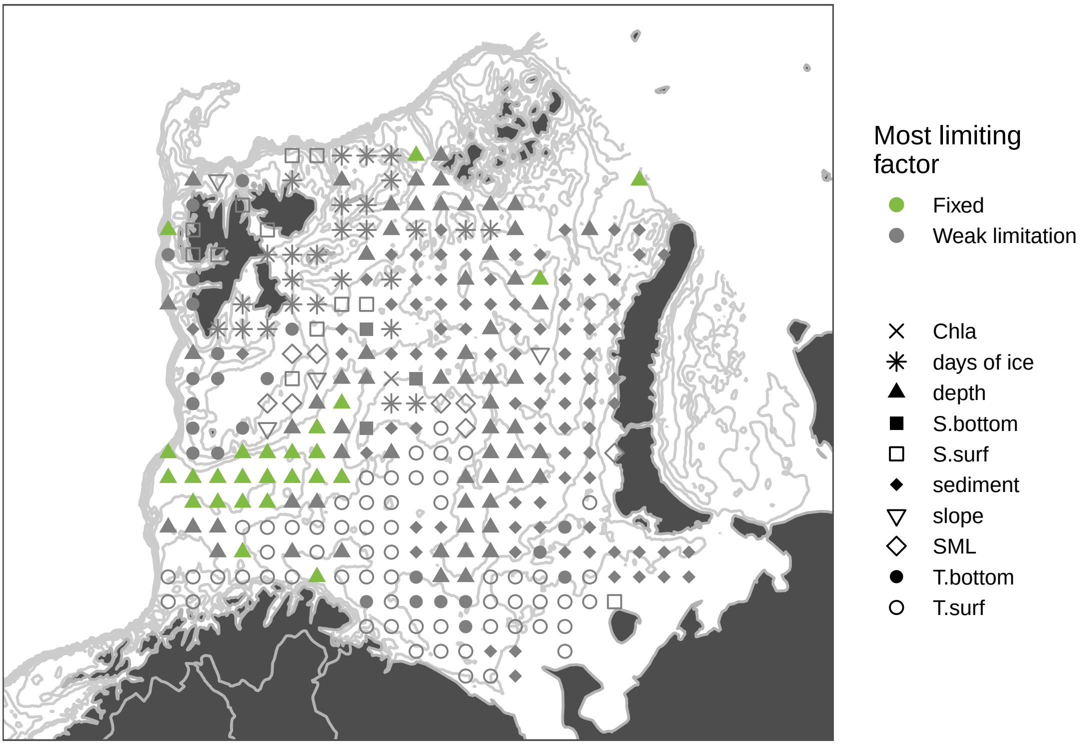

### Icelus

A

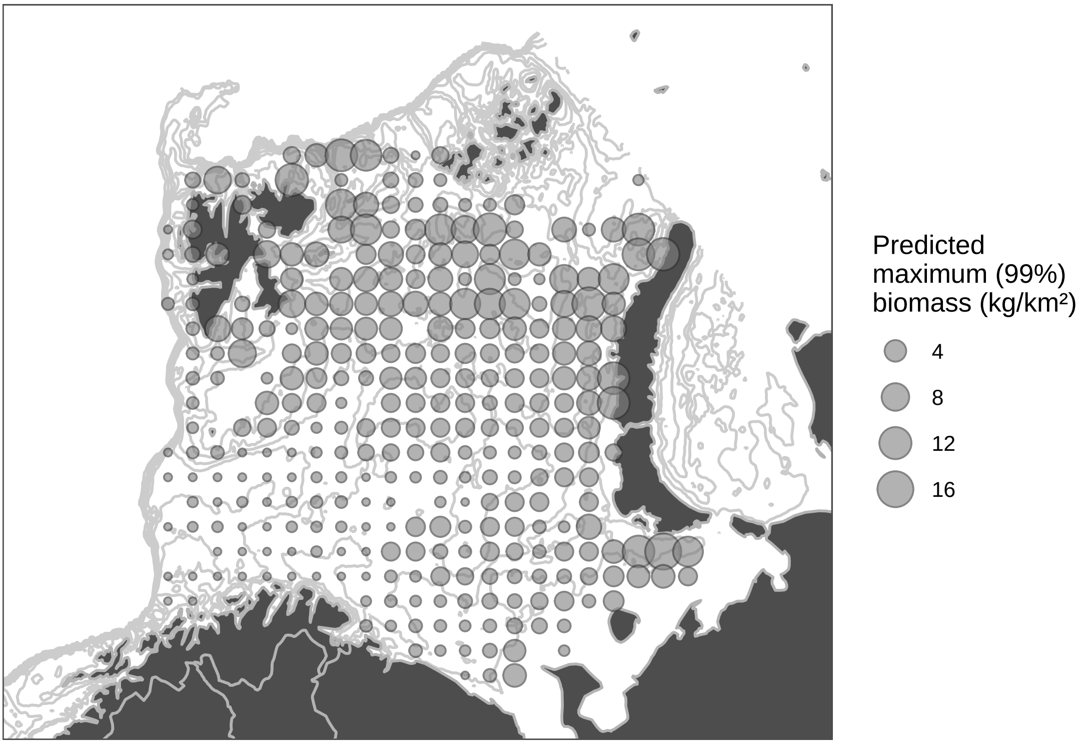
Predicted maximum (99%) biomass (kg/km²)
4
8
12
16

B

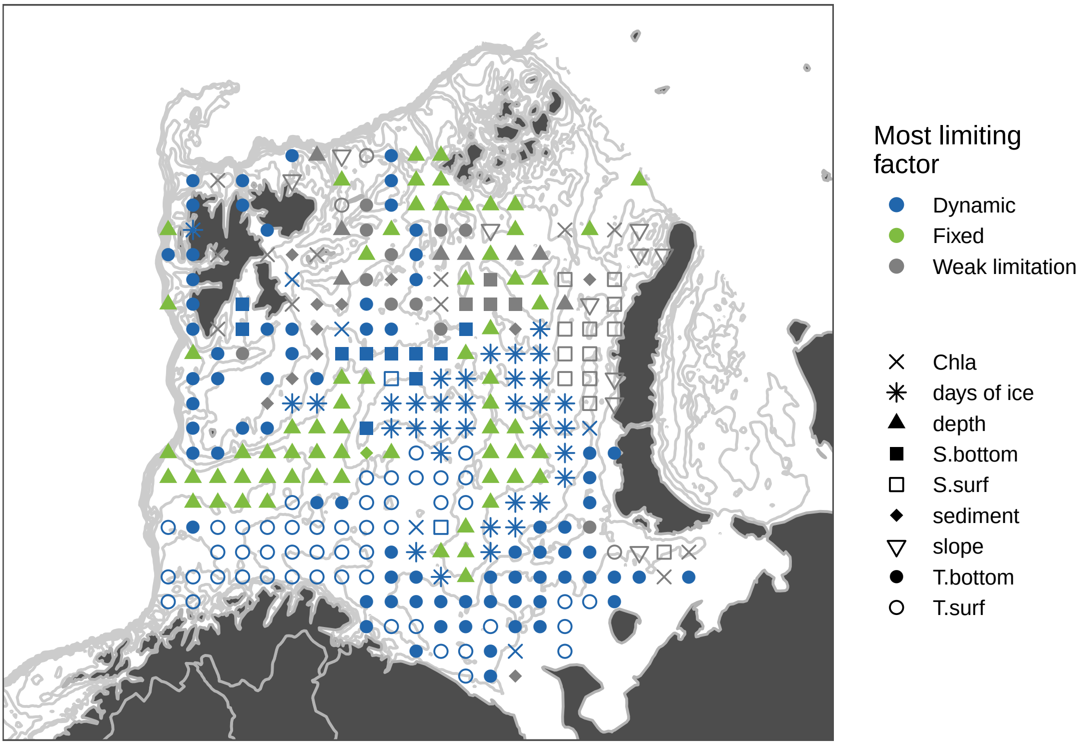
Most limiting factor
Dynamic
Fixed
Weak limitation
Chla
days of ice
depth
S.bottom
S.surf
sediment
slope
T.bottom
T.surf

### Leptagonus decagonus

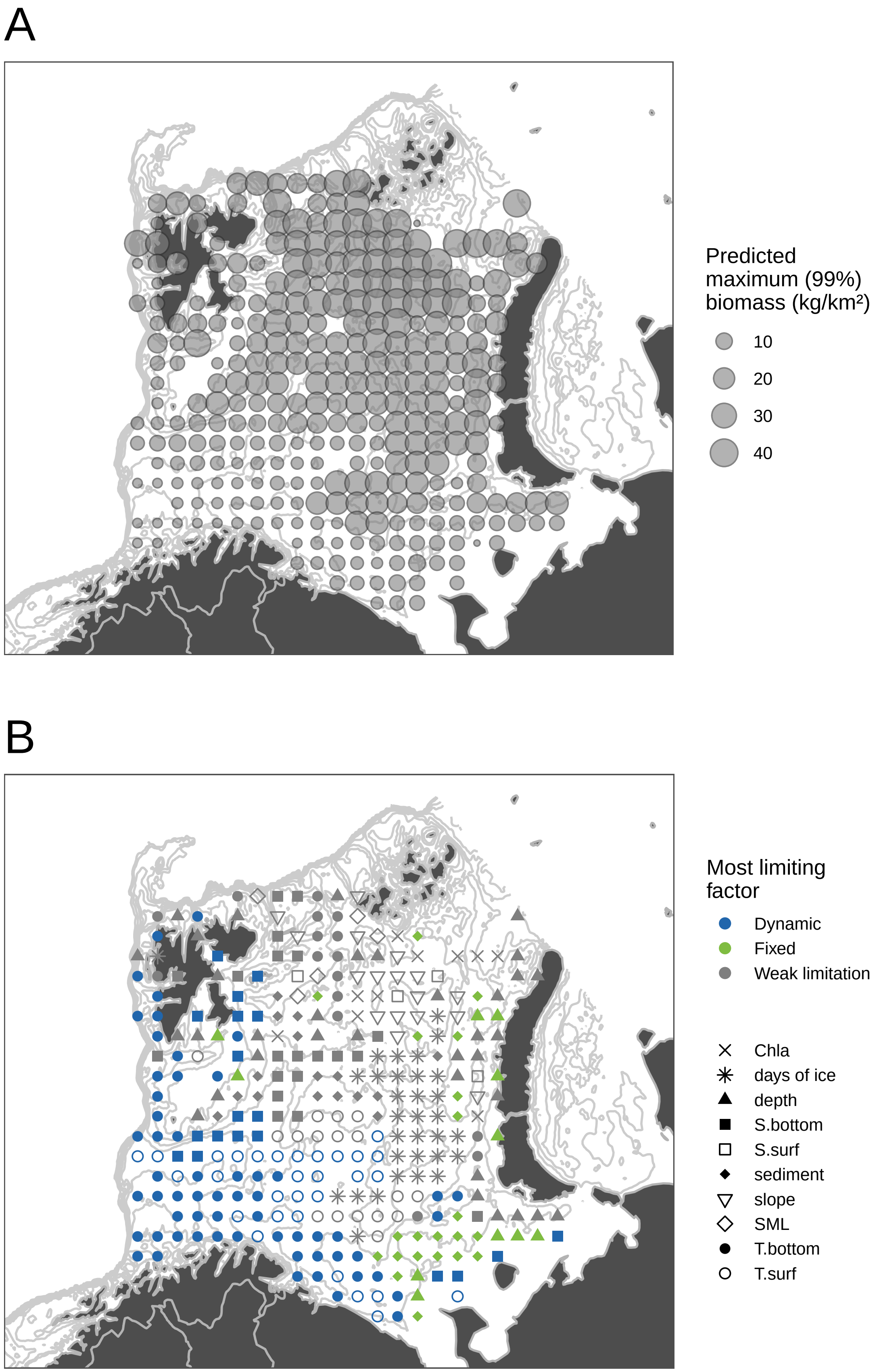

### Leptoclinus maculatus

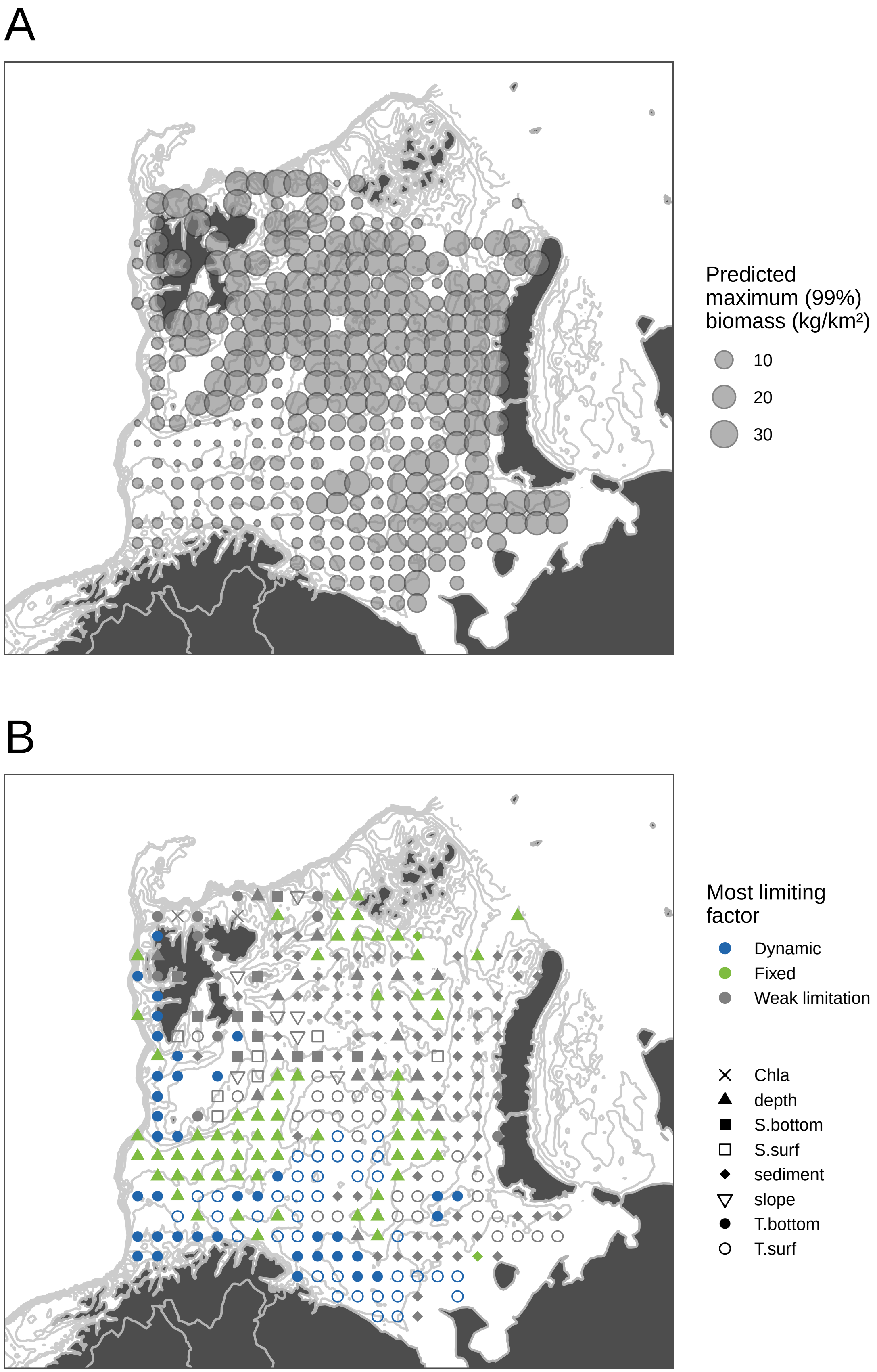

### Liparidae

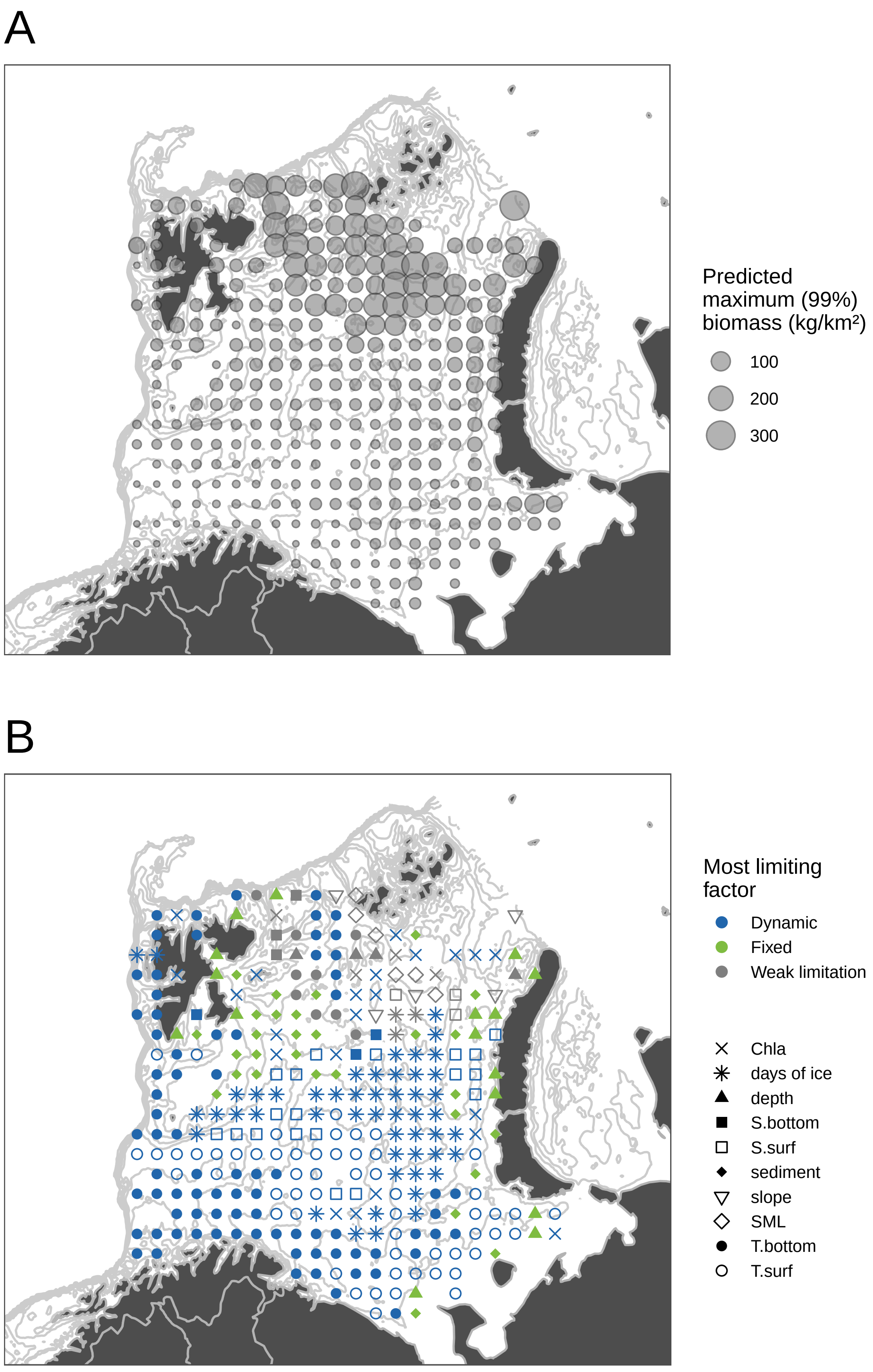

A
Predicted maximum (99%) biomass (kg/km²)
100
200
300
B
Most limiting factor
Dynamic
Fixed
Weak limitation
Chla
days of ice
depth
S.bottom
S.surf
sediment
slope
SML
T.bottom
T.surf

### Lumpenus lampretaeformis

A

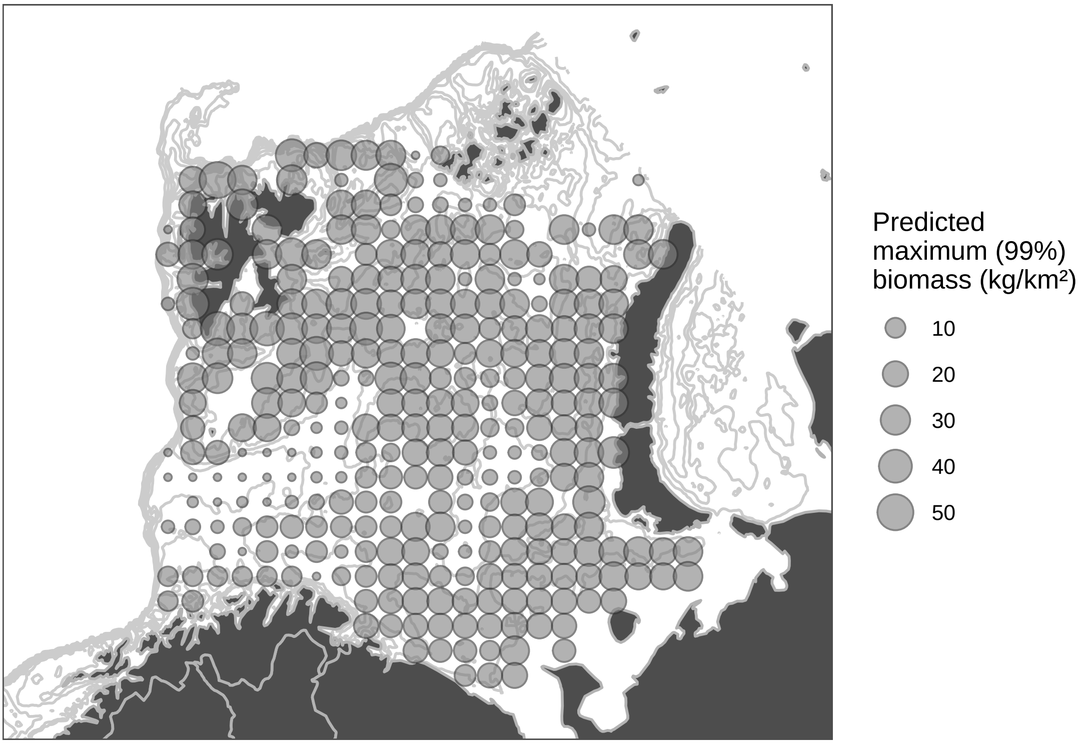


B

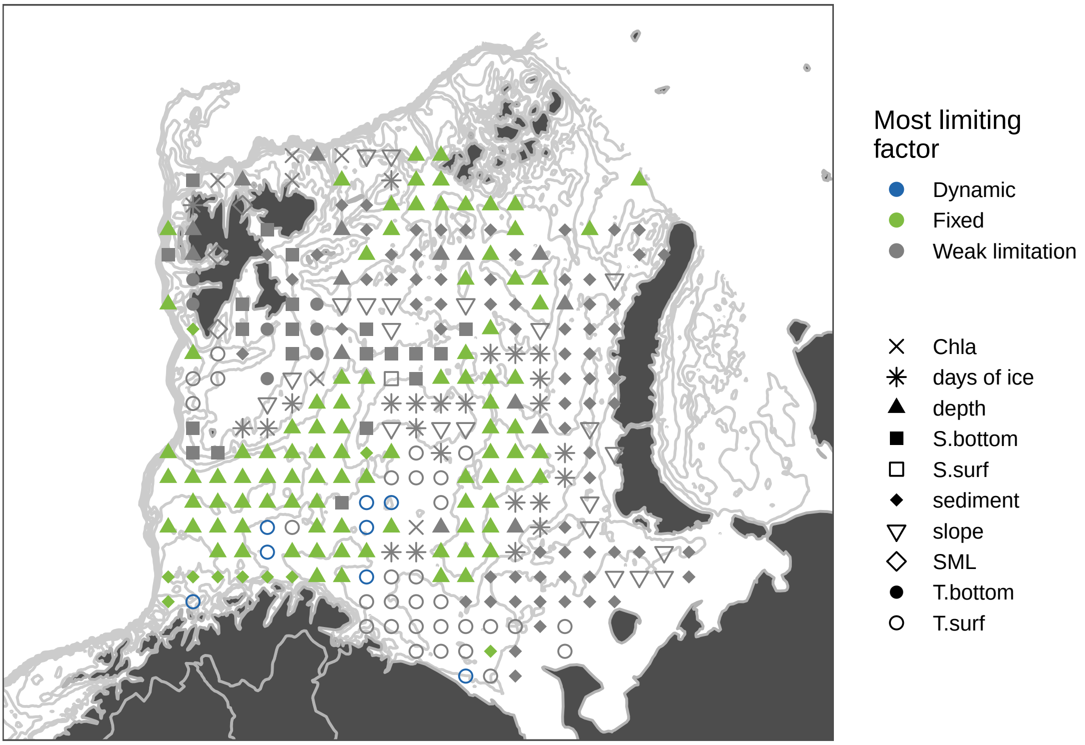

### Mallotus villosus

A

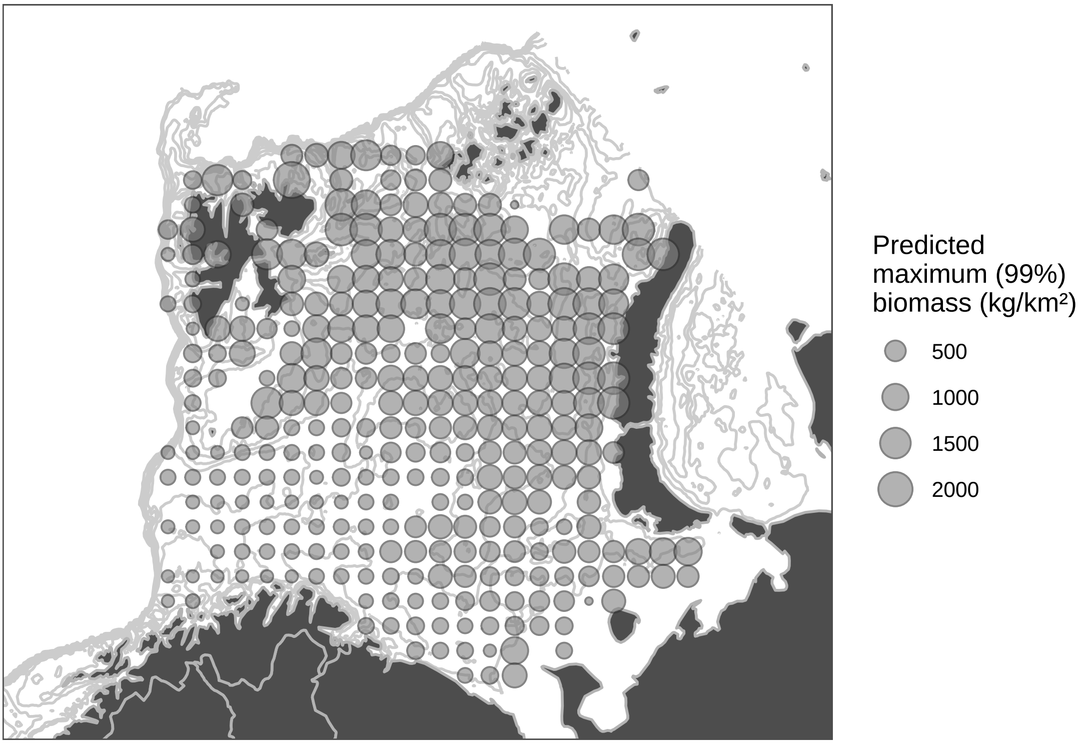


B

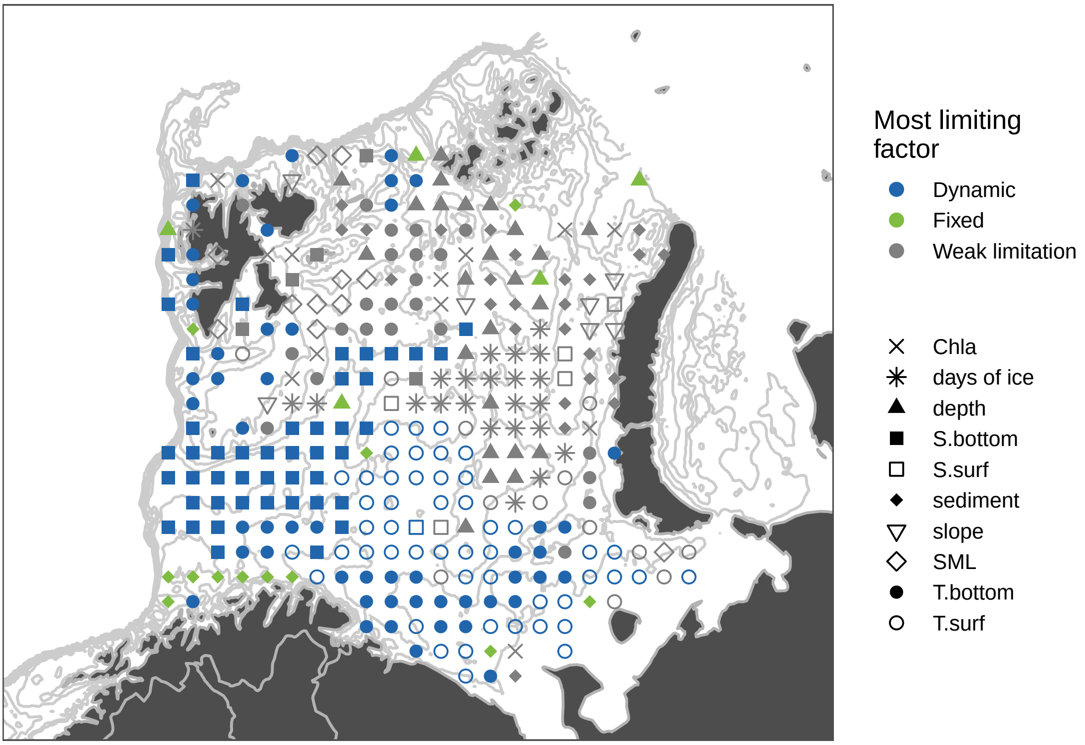

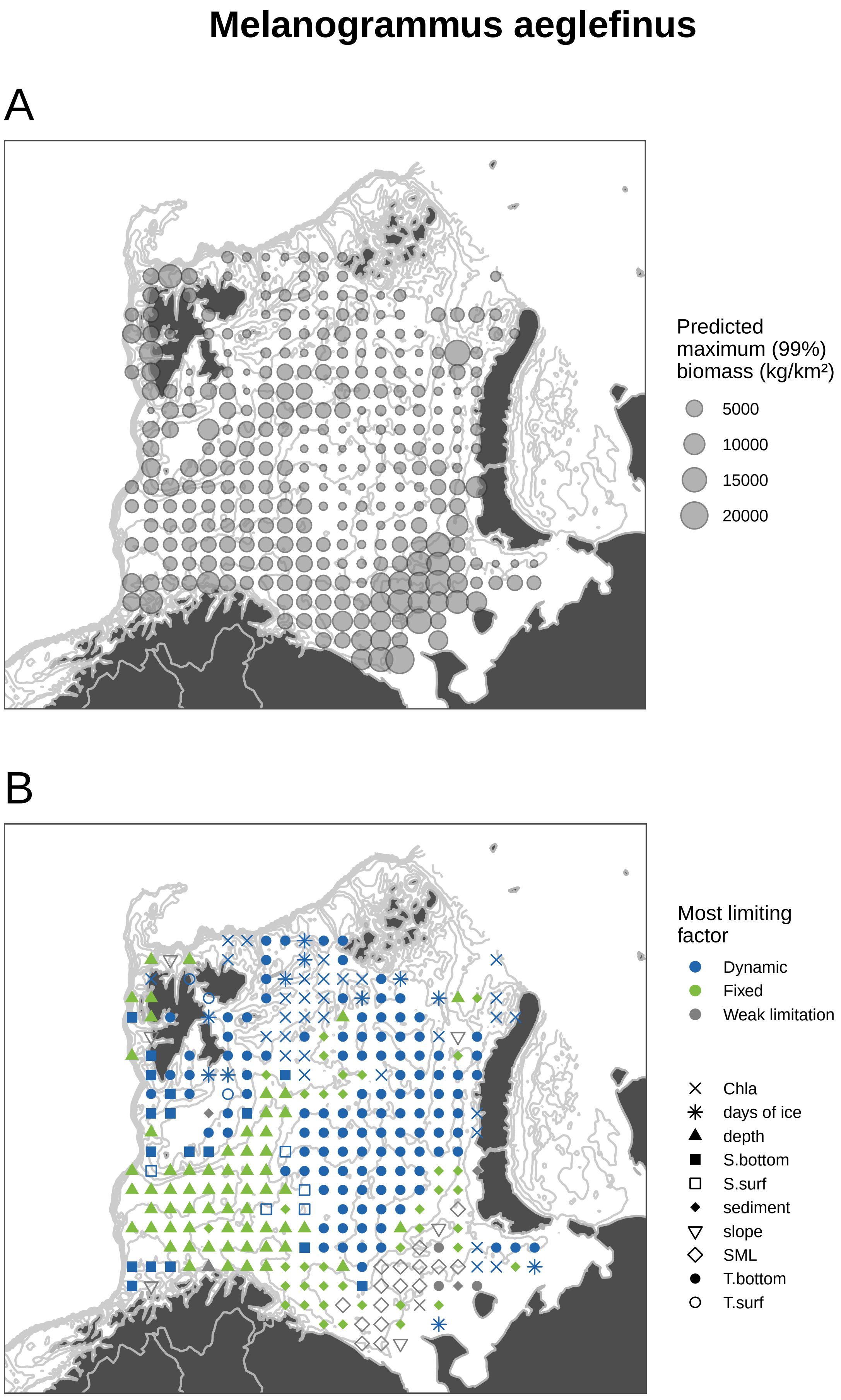
Melanogrammus aeglefinus
A
Predicted maximum (99%) biomass (kg/km²)
5000
10000
15000
20000
B
Most limiting factor
Dynamic
Fixed
Weak limitation
Chla
days of ice
depth
S.bottom
S.surf
sediment
slope
SML
T.bottom
T.surf

### Micromesistius poutassou

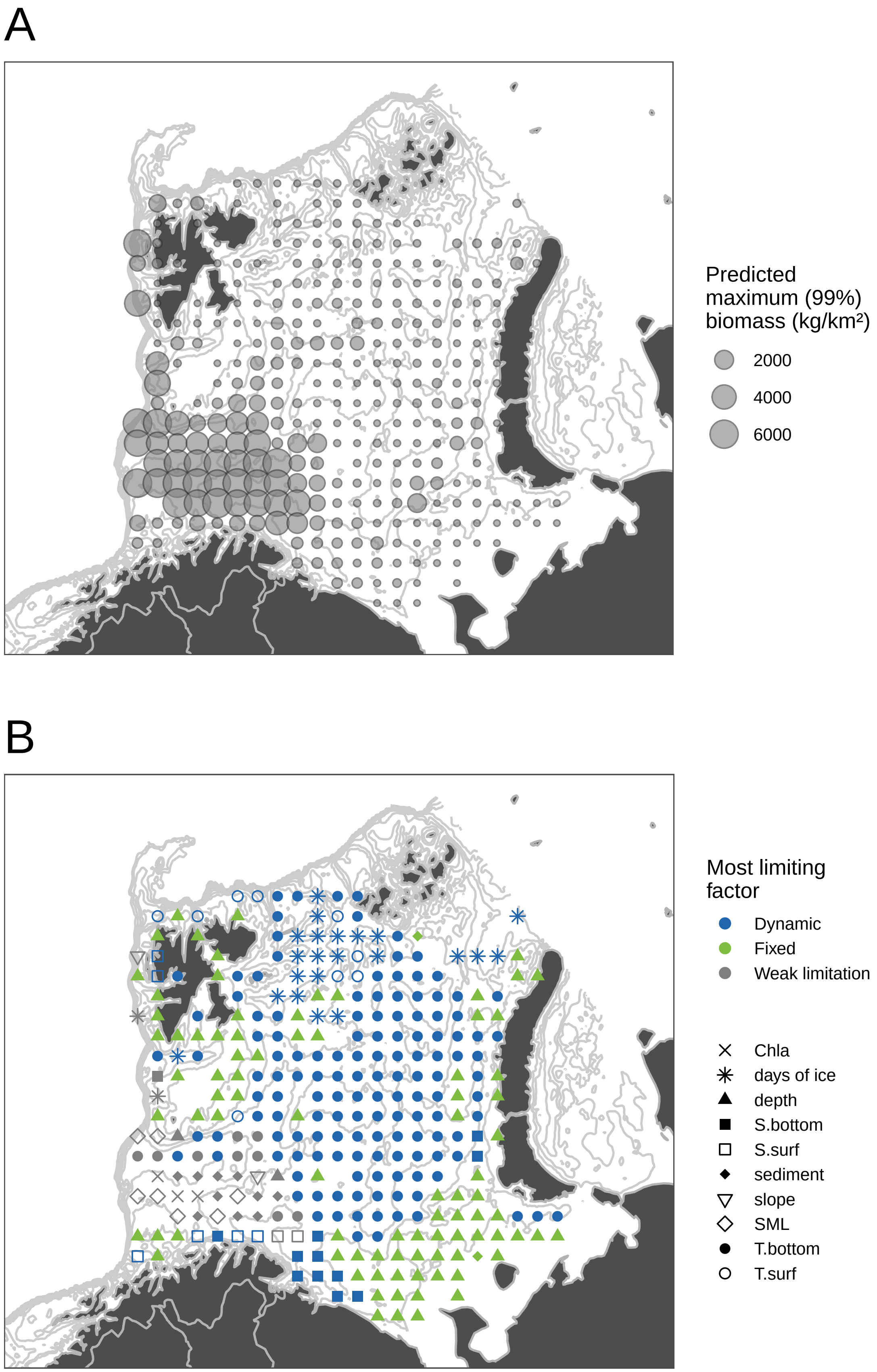

### Pollachius virens

A

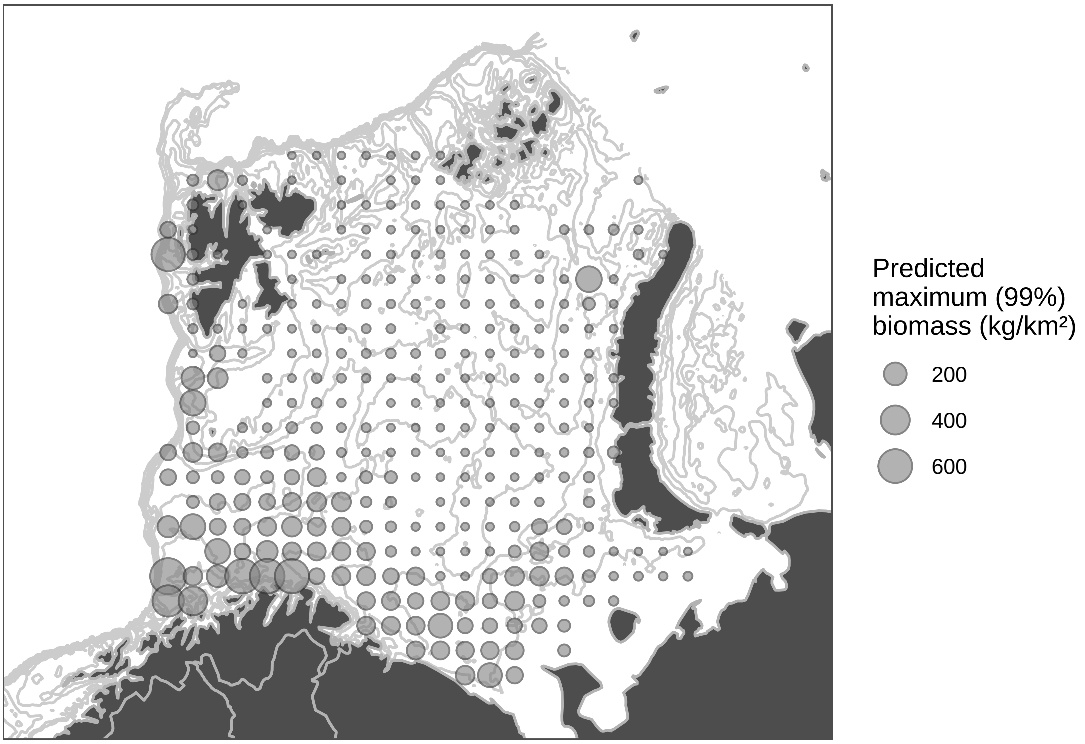


B

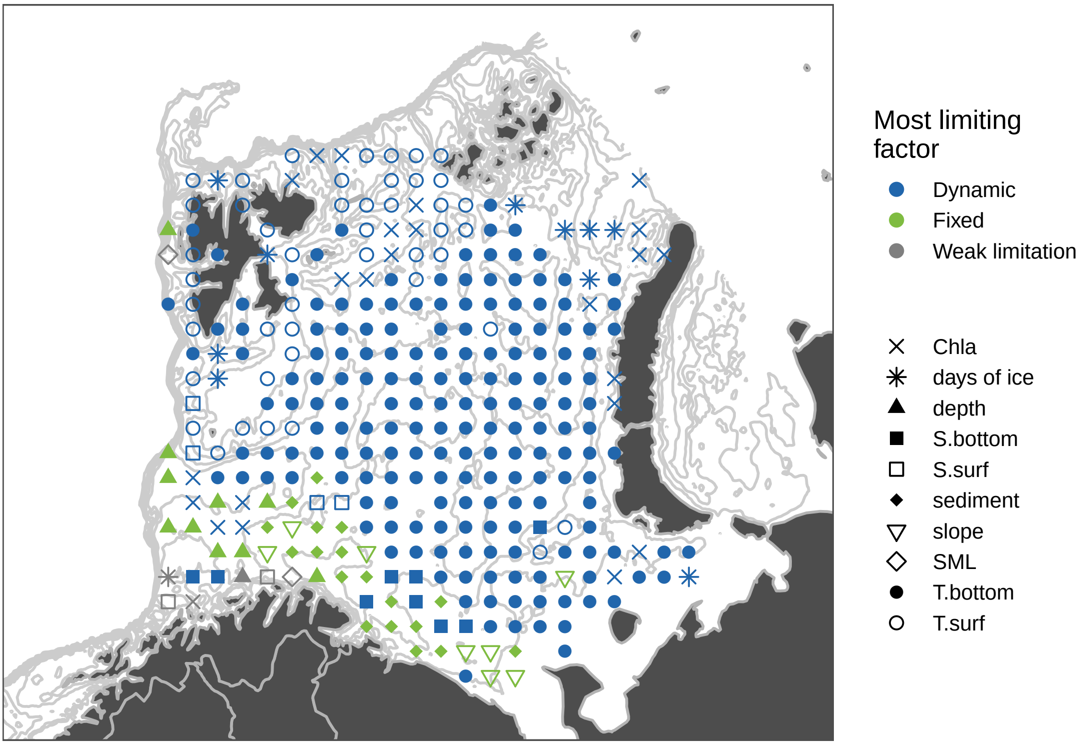

### Reinhardtius hippoglossoides

A

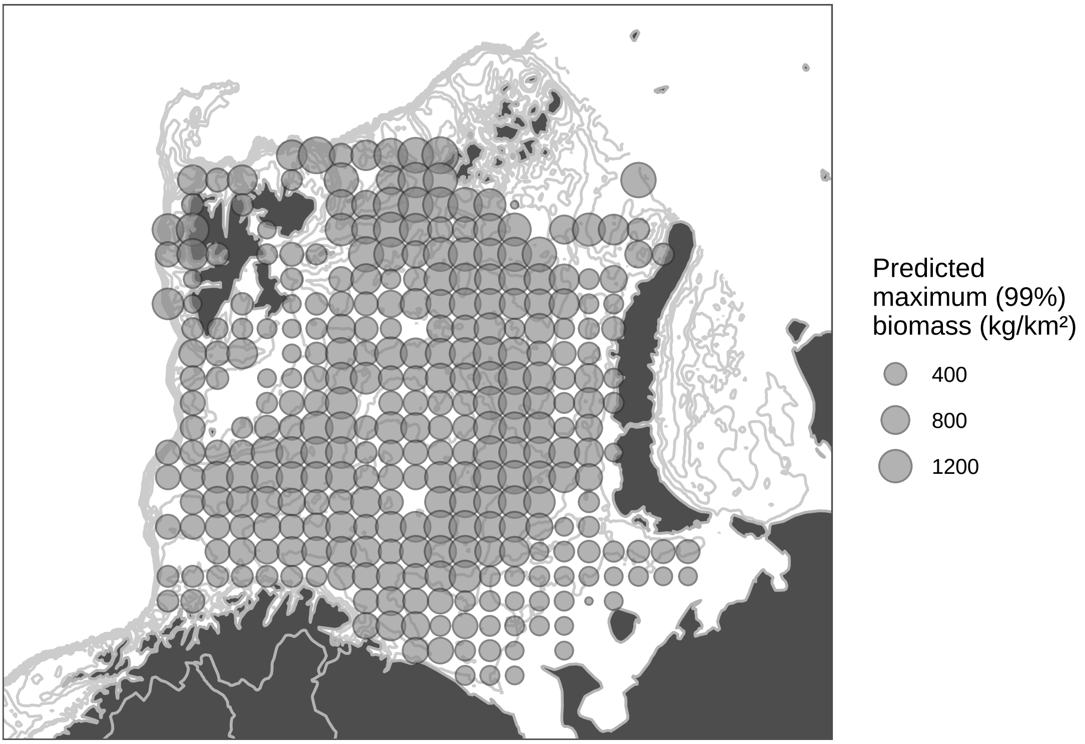


B

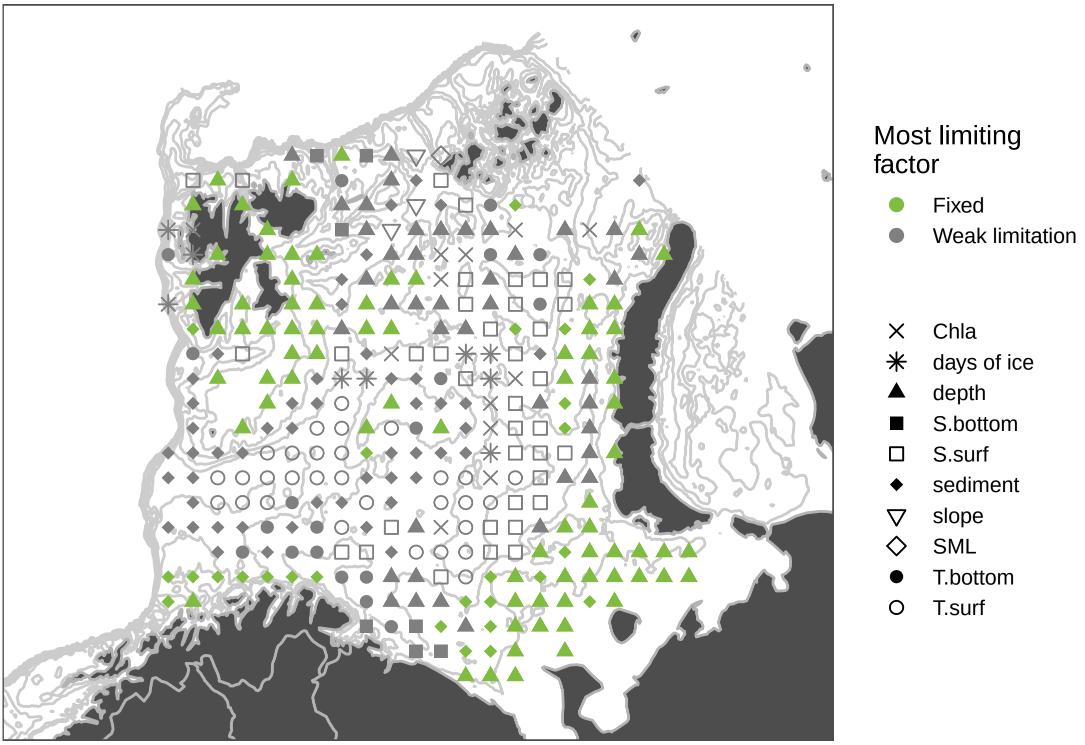

### Sebastes mentella

A

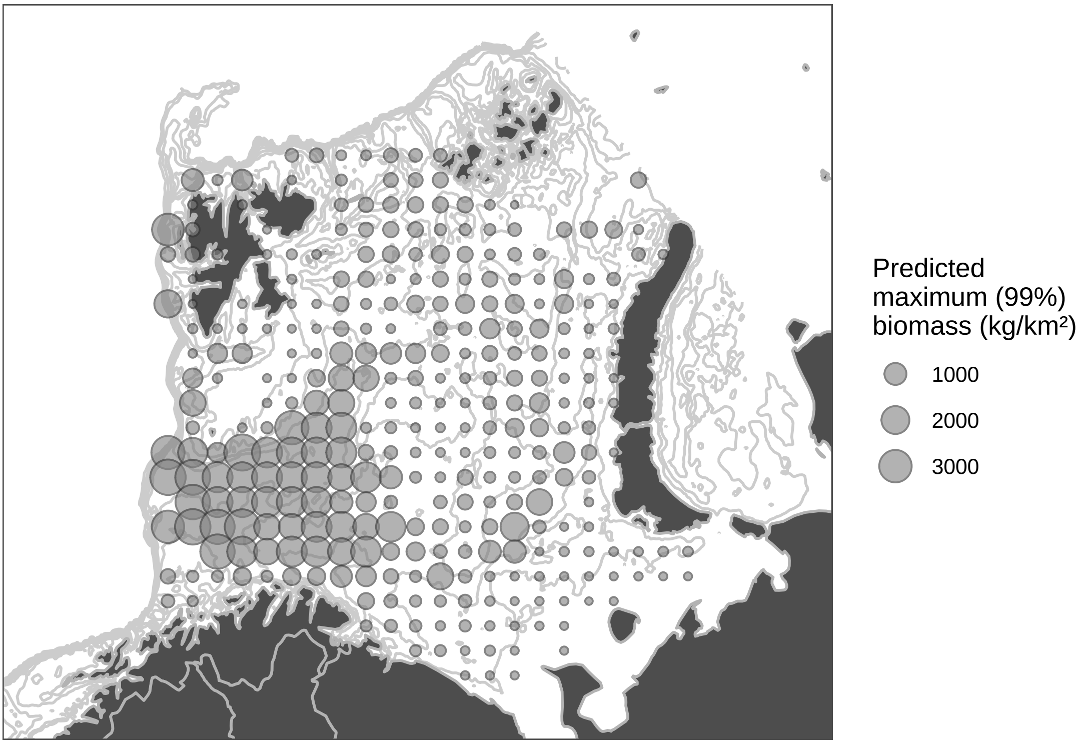


B

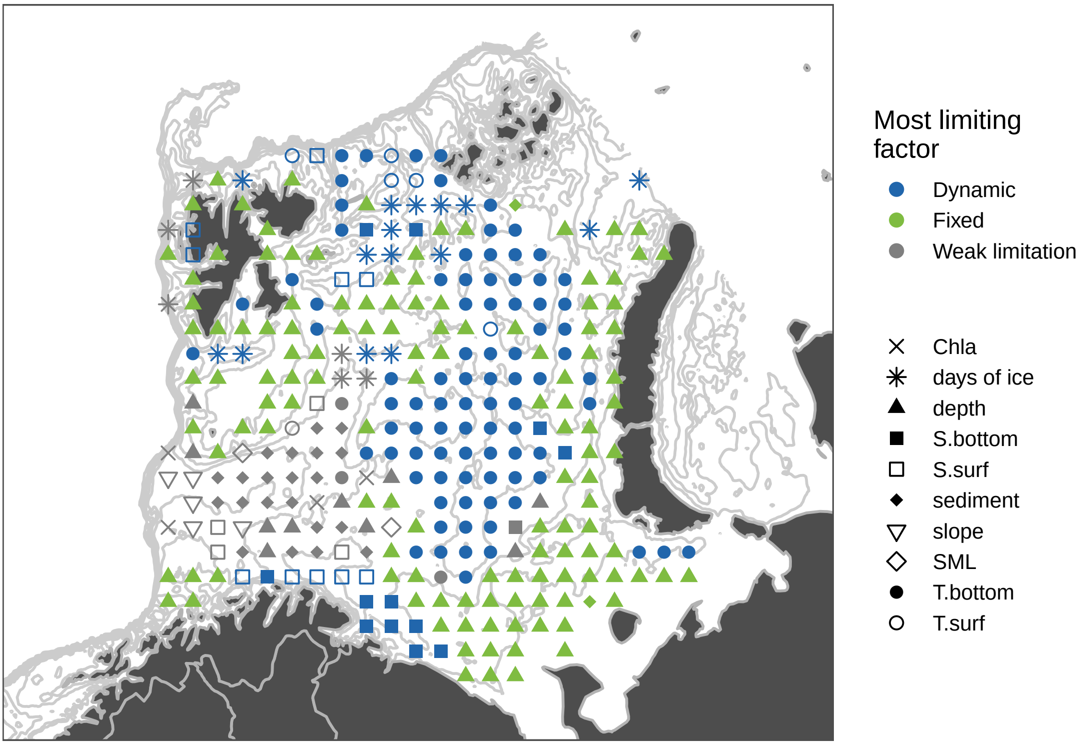

### *Sebastes norvegicus*

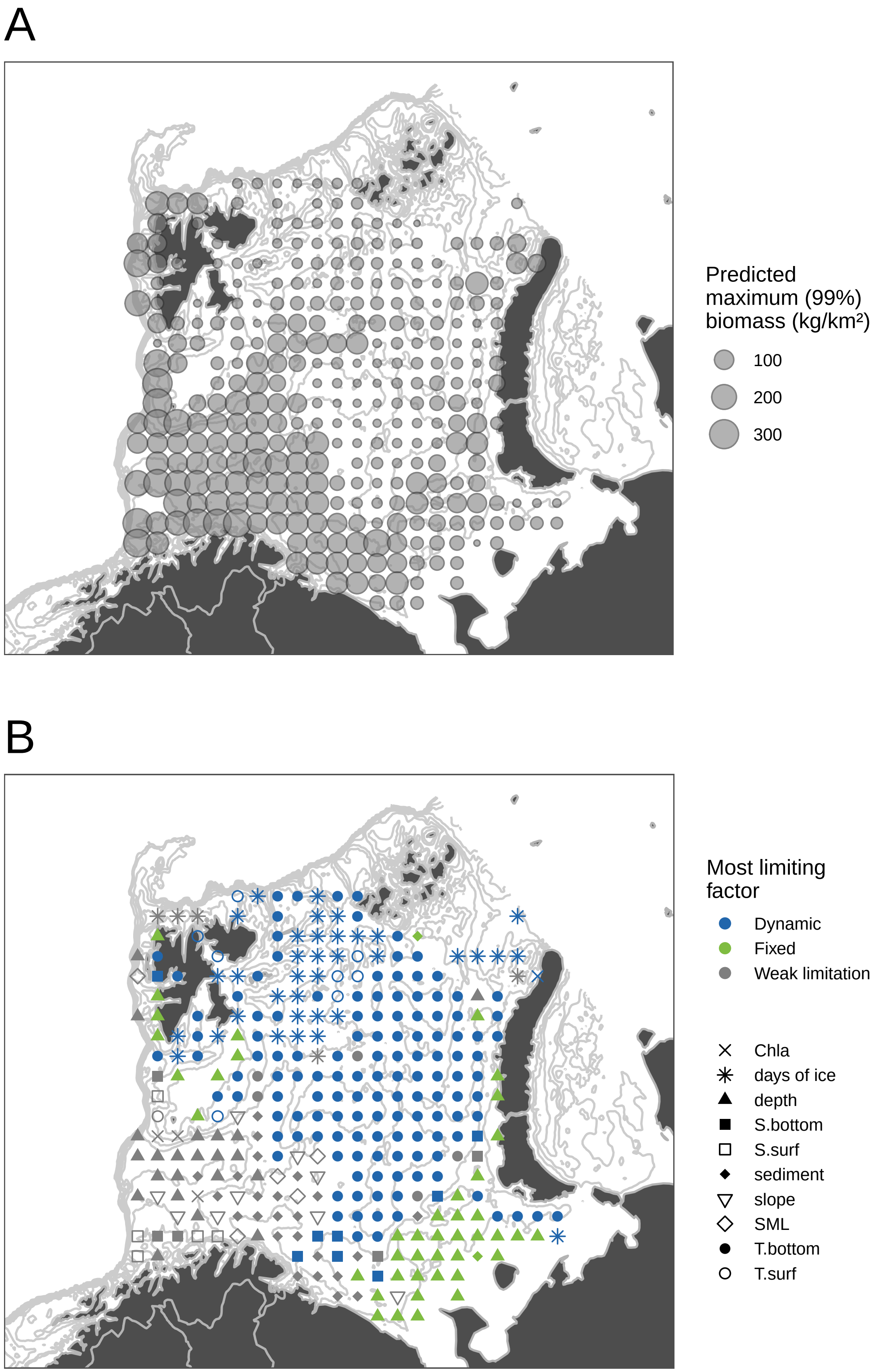

### Sebastes viviparus

A

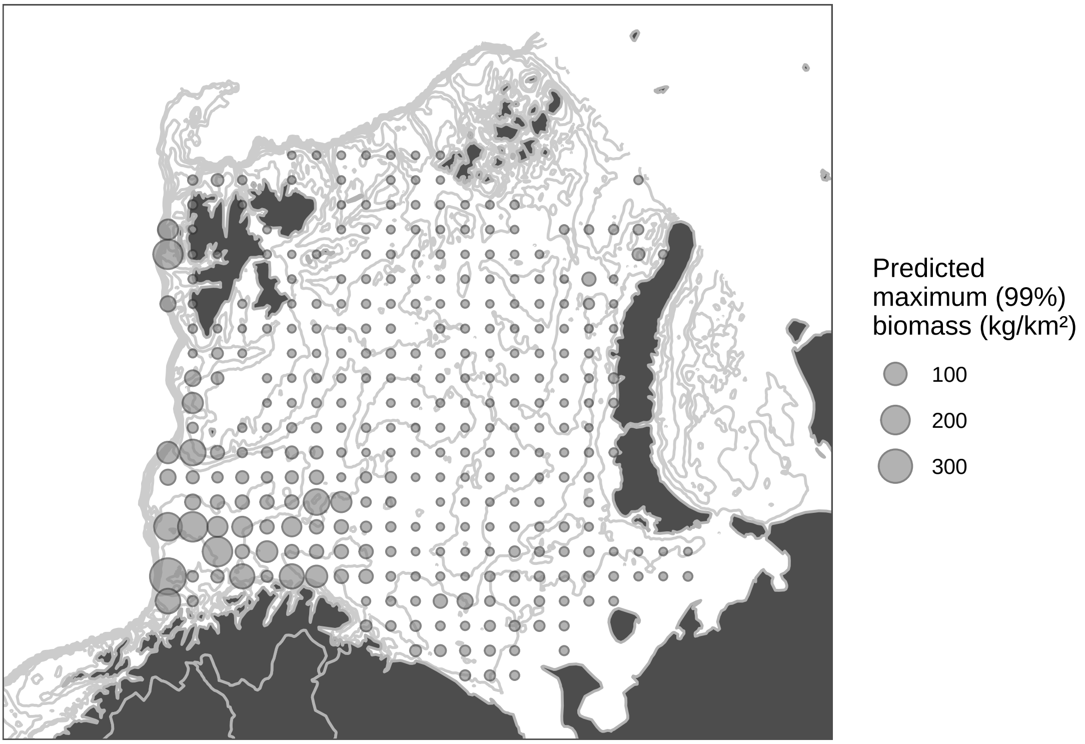


B

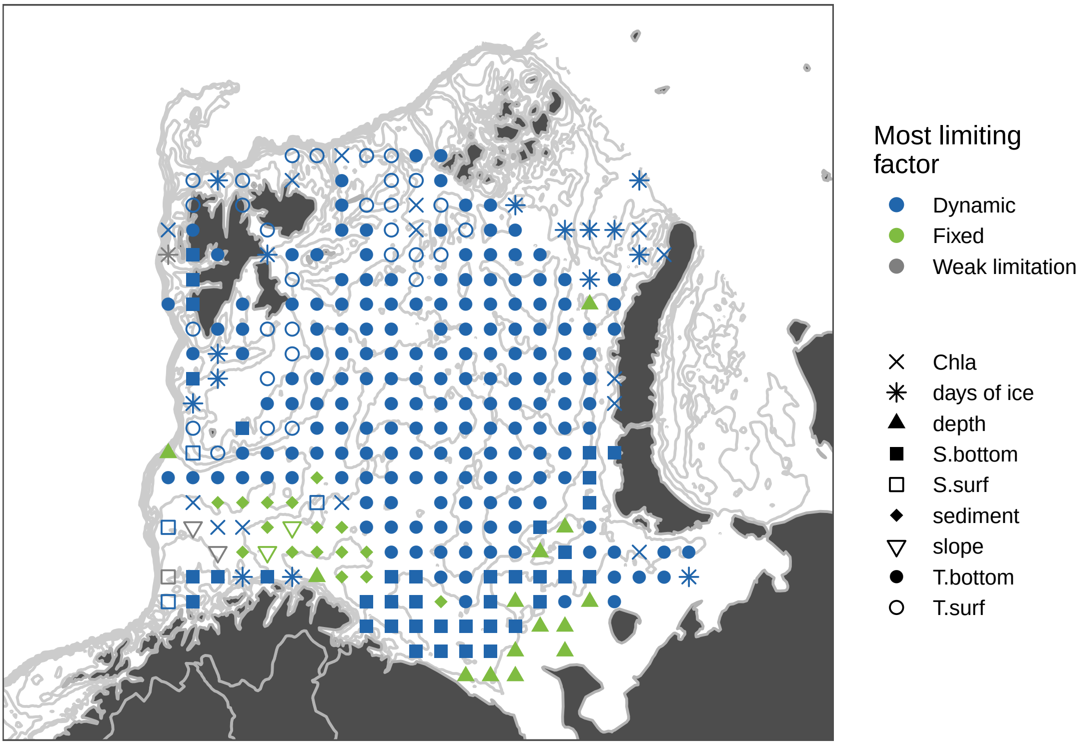

### Triglops murrayi

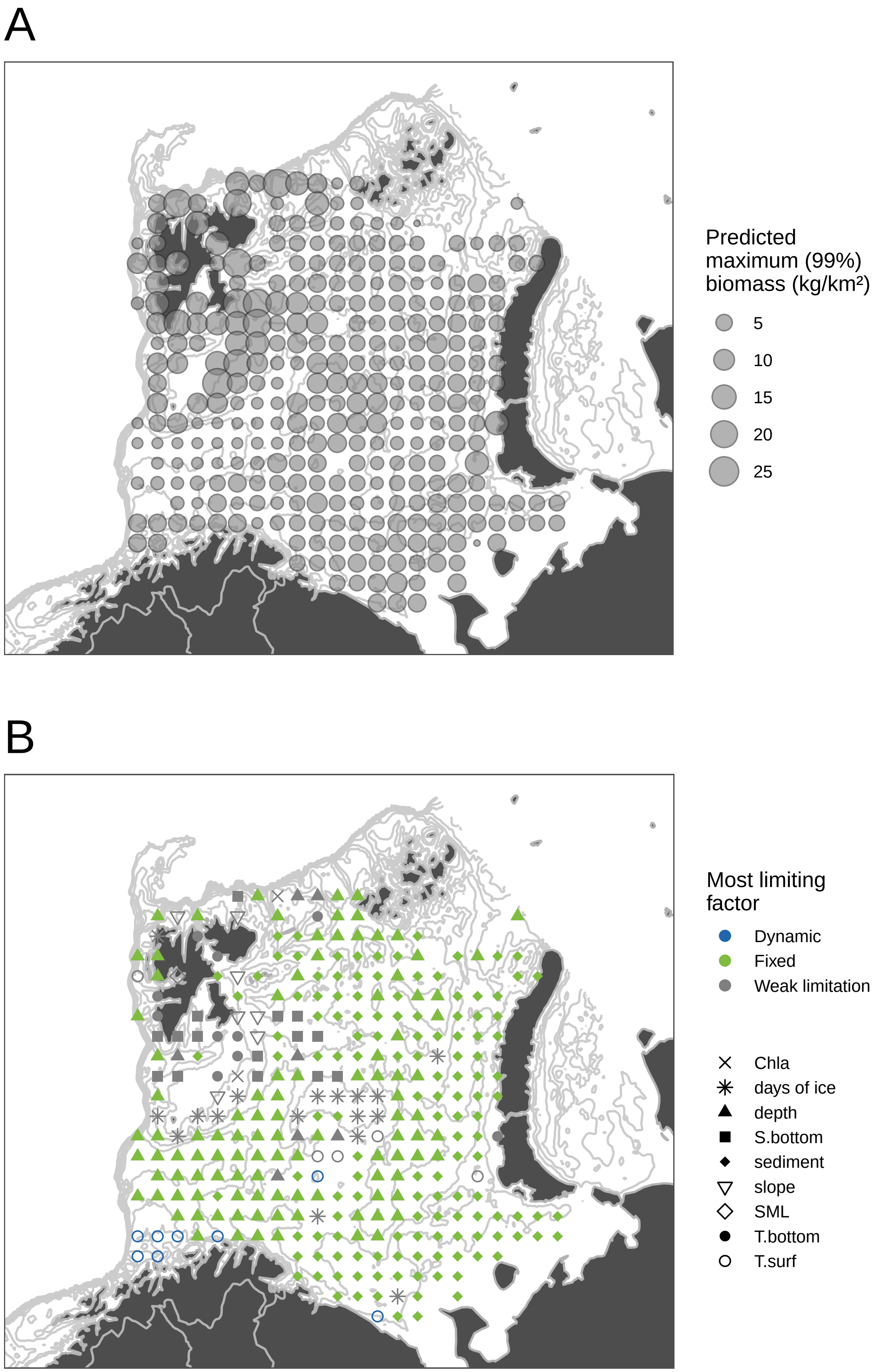

### Triglops nybelini

A

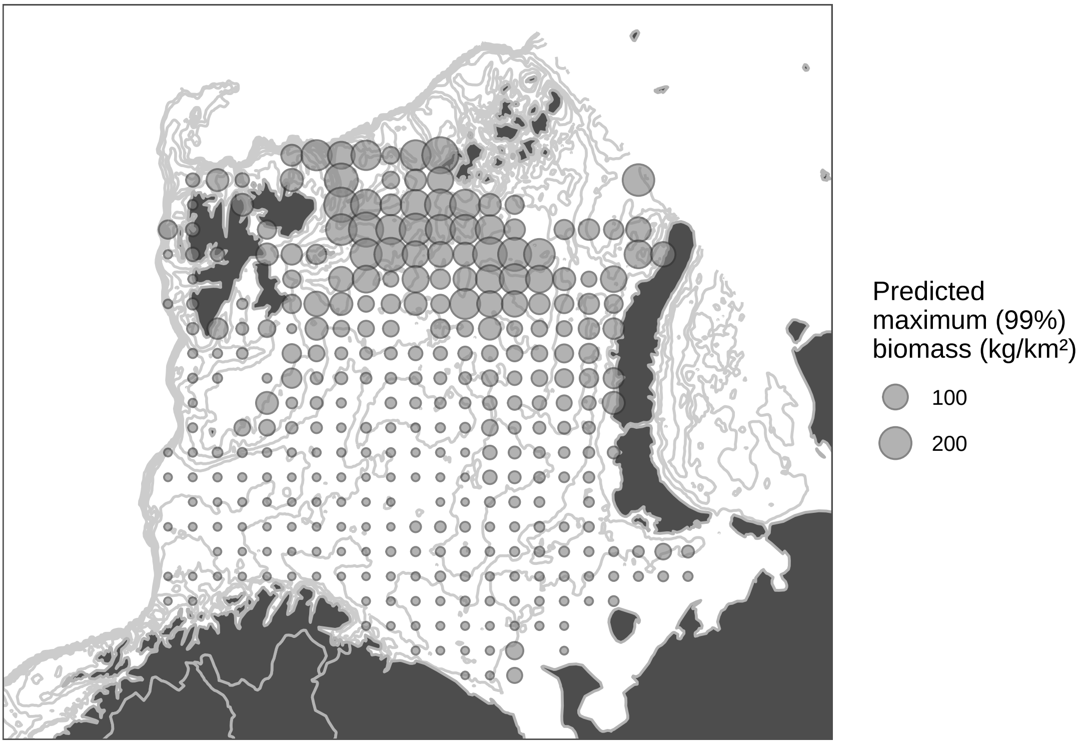


B

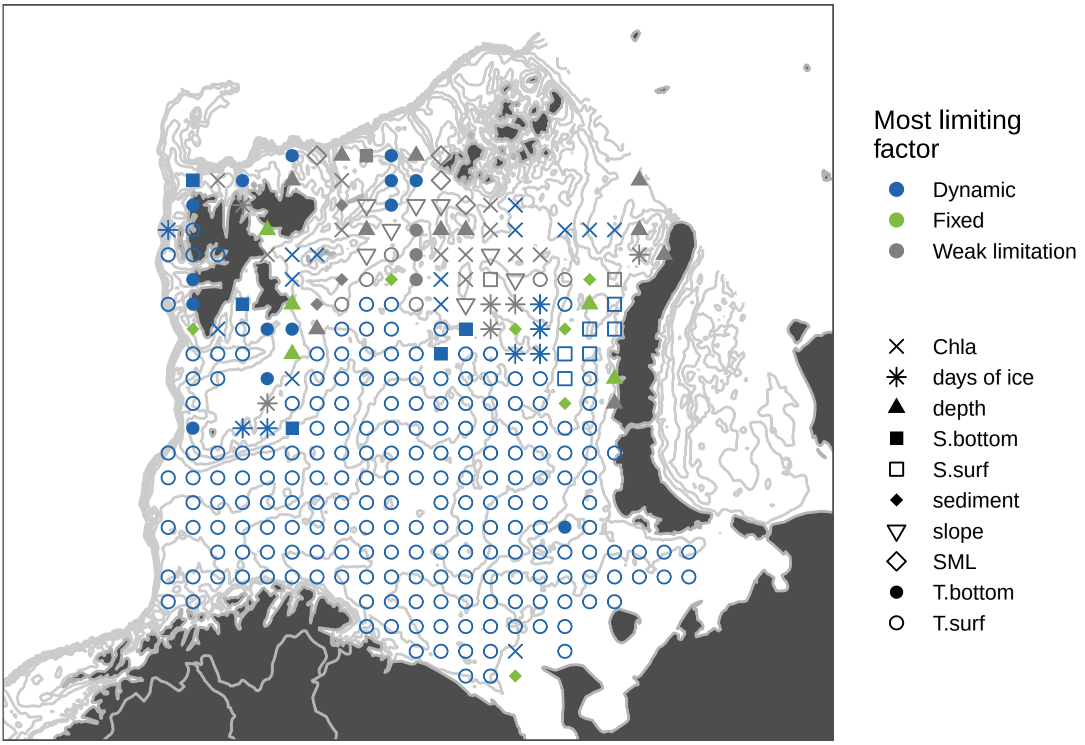

### Triglops pingelii

A

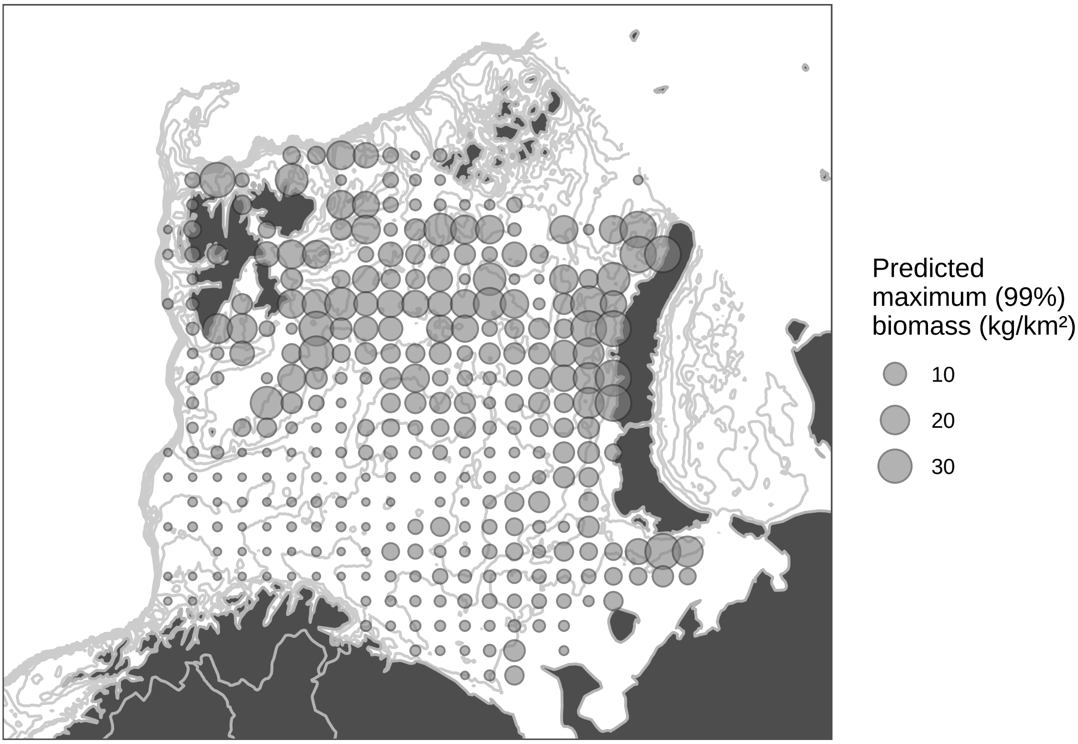


B

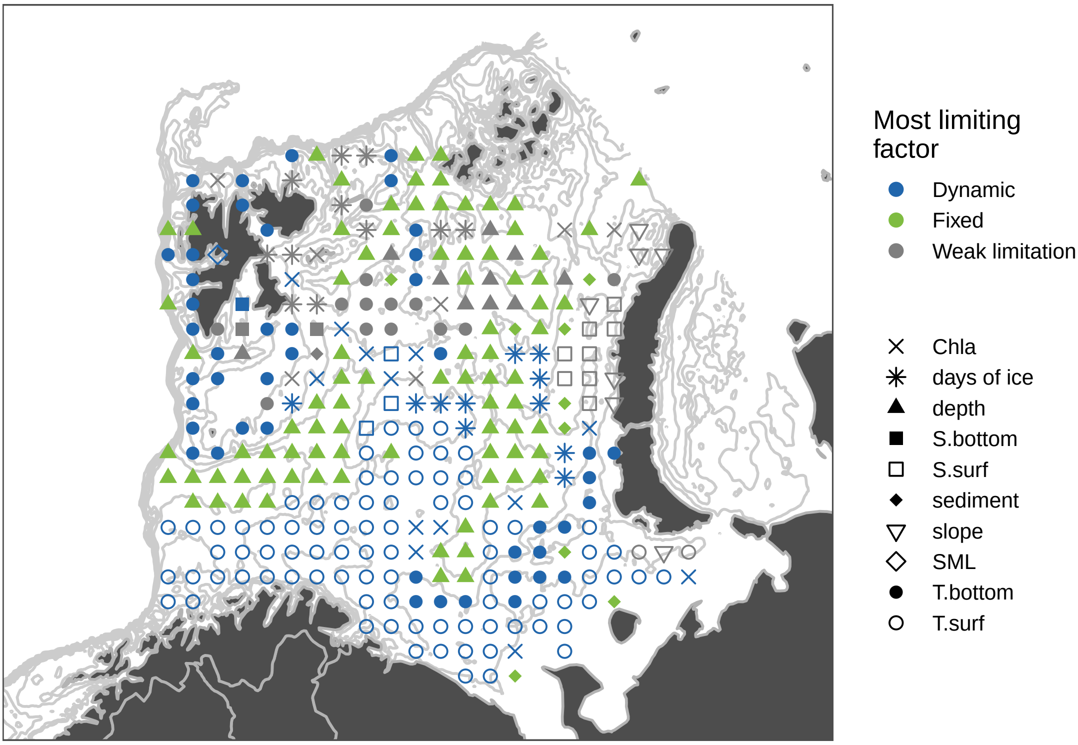

### Trisopterus esmarkii

A

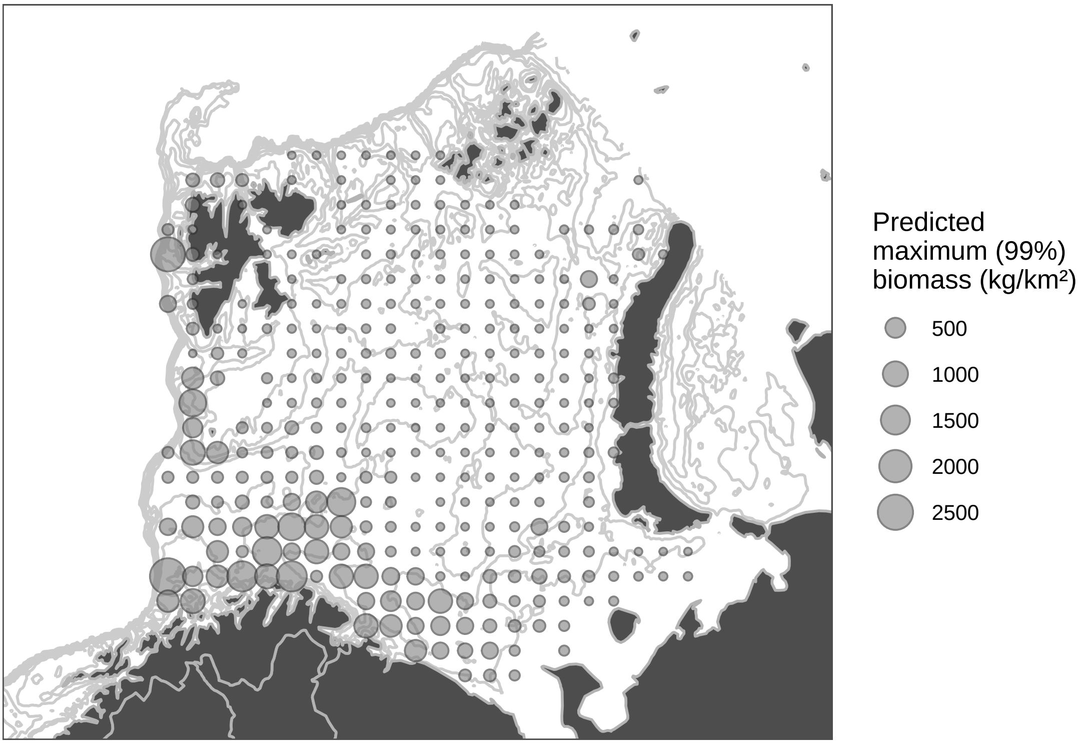


B

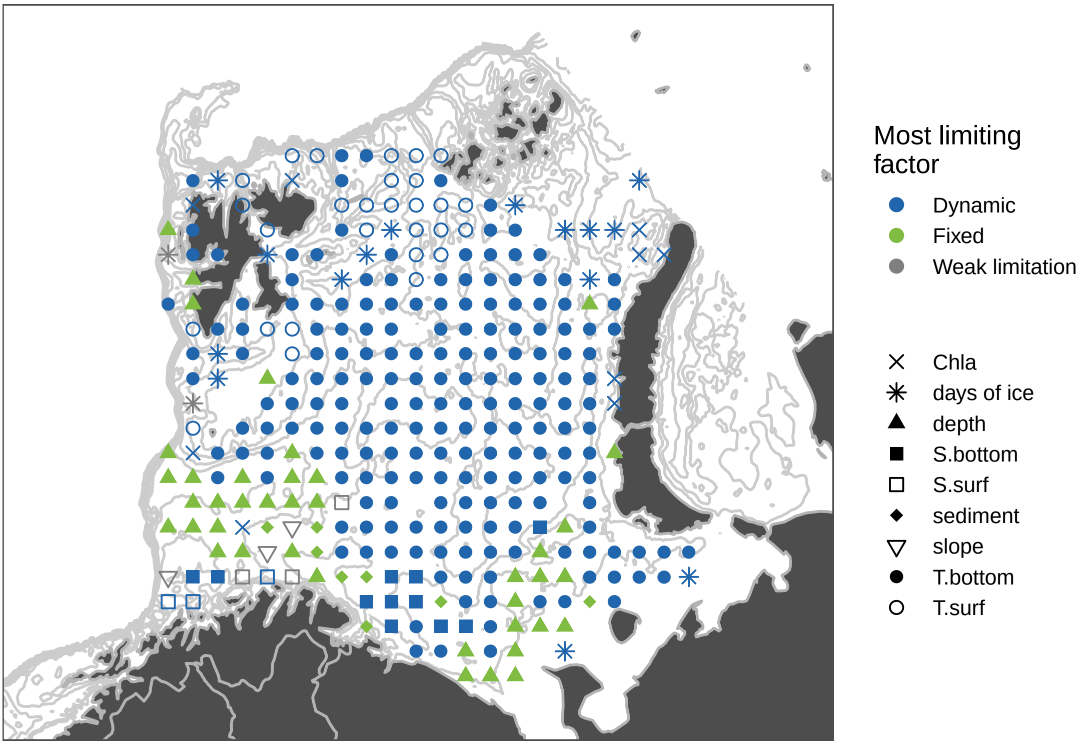

### Zoarcidae

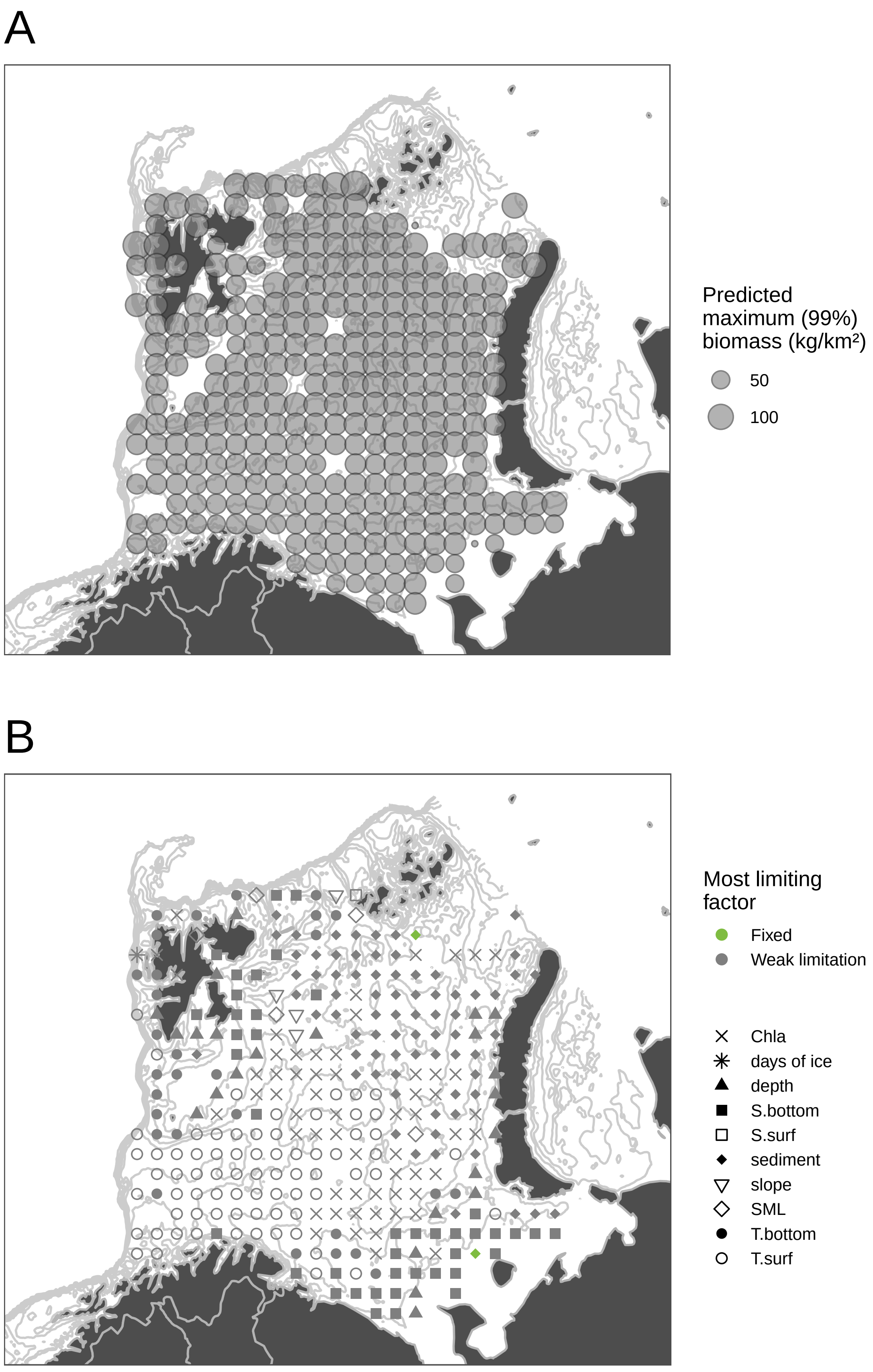
A
Predicted
maximum (99%)
biomass (kg/km²)
50
100
B
Most limiting
factor
Fixed
Weak limitation
Chla
days of ice
depth
S.bottom
S.surf
sediment
slope
SML
T.bottom
T.surf
